# A novel approach to map the causal impact of brain stimulation on semantic processing with language models

**DOI:** 10.1101/2025.05.06.652092

**Authors:** Andrea Bruera, Gesa Hartwigsen

**Affiliations:** Research Group Cognition and Plasticity, Max Planck Institute for Human Cognitive and Brain Sciences, Leipzig, Germany; Cognitive and Biological Psychology, Wilhelm Wundt Institute for Psychology, Leipzig University, Germany

**Keywords:** brain stimulation, TMS, semantics, semantic models, language models, causality, semantic similarity, surprisal

## Abstract

Non-invasive brain stimulation (NIBS) studies on semantic cognition hold the promise of revealing the functional relevance of brain areas through causal intervention. A primary challenge, however, is that findings are often interpreted through binary distinctions between sets of stimuli (e.g. related/unrelated words, same/different semantic category). This approach ignores the analysis of individual words, which mirrors every-day language use and is crucial for understanding semantic cognition. In this work, we used semantic similarity, as measured by a language model, to investigate how Transcranial Magnetic Stimulation (TMS) effects on semantic cognition unfold at the level of individual words. We re-analyzed 5 publicly available TMS datasets, covering multiple stimulation sites and lexical semantics tasks. We propose a simple methodology that can straightforwardly be applied to any TMS experiment on semantic cognition, and showcase its potential to generate new insights. We modelled trial-level response times using the language model and computed the correlation between the two. We also repeated the analyses for two lower-level variables (word frequency and length). Importantly, for each dataset, we compared correlations for effective and control (sham or vertex) stimulation conditions. We found that, for the language model, correlation was almost always significantly different depending on the type of stimulation (effective or control). Our results provide evidence that the stimulation effect interacts with the meaning of individual words. However, a similar pattern emerged in some cases for word frequency and length, suggesting that the effects of TMS on cognition can be widespread, well beyond their intended functional target. Collectively, our results demonstrate that language models provide new insight into the impact of neurostimulation on semantic processing, complementing standard measures.

## Introduction

Semantic cognition is central to human communication and interaction in everyday life. The meaning of the words we use can be captured by dividing them into distinct categories: animate as opposed to inanimate; objects as opposed to actions; concrete as opposed to abstract words. Such binary distinctions among sets of words have proven to be an insightful window into the way our brain processes semantics (Binder et al., 2005; Caramazza & Shelton, 1998; Martin, 2007). Yet, words are represented and used in ways that go well beyond dichotomies and categories (Huth et al., 2012; Trott & Bergen, 2023; van Hoef et al., 2023). For instance, a cat and an elephant belong to the same category - animals. However, while I regularly interact with cats, I have never seen an elephant. The vast differences in real-life sensory and social experiences I have with the two dramatically change the way my brain stores and accesses the meaning of the word ‘cat’ as opposed to ‘elephant’ (Brysbaert et al., 2018; Charest et al., 2014; Renoult et al., 2019; Yonelinas, 2002). This impact is so pronounced that the fact that they belong, ontologically, to the same category may become almost irrelevant (Bozeat et al., 2002; Bruera & Poesio, 2025; Hoffman, 2018; Rogers et al., 2015).

In recent years, the cognitive neuroscience of language has started taking seriously the investigation of how our brains store and use individual concepts, as opposed to binary, mutually exclusive categories (Fernandino et al., 2022; Huth et al., 2016; Mitchell et al., 2008; Pereira et al., 2018). Despite decades of effort, it is a longstanding, unresolved question how semantic knowledge - which has been shown to involve disparate pieces of information (e.g. sensory, motor, social, abstract) across distant areas of the brain - is retrieved, brought together and used in natural language and behaviour (Binder et al., 2009; Calzavarini, 2024; Lambon Ralph et al., 2017). This puzzle still eludes our understanding, among other reasons, because it has been extremely hard to obtain consistent and reliable evidence about the causal role of specific areas: which brain area is playing a necessary role, and when (Drijvers et al., 2025; Jackson, 2021; Schroën et al., To address the issue of causality, non-invasive brain stimulation (NIBS) is a key technique in the cognitive neuroscientist’s toolbox (Bergmann et al., 2016). In particular, transcranial magnetic stimulation (TMS) has been used for more than two decades now to transiently modulate brain function and probe the relevance of specific brain areas for distinct cognitive operations (Bergmann & Hartwigsen, 2021; Pitcher et al., 2021). In a typical cognitive experiment, healthy participants repeat the same task with effective and ineffective stimulation of a given area. This allows the researcher to probe the question of what would be the effect on a subject’s behaviour during a cognitive task if the regular functioning brain area *X* was altered. Relative to the study of patients with brain lesions, TMS-induced perturbations allow for relatively precisely localized stimulation effects, targeting individual brain regions and allowing to formulate very specific research questions about the relevance of specific brain areas for different cognitive sub-processes at different time points (Hallett, 2007; Hartwigsen, 2015; Jefferies, 2013; Valero-Cabré et al., 2017).

To address theoretical questions in semantics, TMS has been used in a variety of ways. First, to complement results in stroke patients, either trying to induce a category-specific deficit (Cattaneo et al., 2010; Pobric et al., 2010; Thompson et al., 2017) or to solve conflicting reports (Devlin et al., 2003; Finocchiaro et al., 2015). Secondly, to test existing models of semantics in the brain - such as the so-called hub-and-spoke and the controlled semantic cognition theories of semantic processing (Chiou et al., 2018; Jefferies, 2013) or the embodied cognition framework (Pulvermüller et al., 2005). Third, framing TMS perturbation in a network-based approach, to assess its effects beyond the stimulated brain region (Binney & Lambon Ralph, 2015; Hartwigsen et al., 2016). Finally, to understand the exact role in semantic processing of specific areas (e.g. (Davey et al., 2015; Gatti et al., 2020; Pobric et al., 2008)).

However, the impact of TMS studies on semantics is significantly limited by the focus on the comparisons between dichotomous categories or conditions (such as semantically related/unrelated, same/different semantic category), missing on the uniqueness of the meaning of individual words. Reaching this level of concept-specific granularity in our understanding of the causal promise of TMS bears exciting potentials: not only revealing previously unattainable evidence about semantic cognition, but also bringing this stimulation approach closer to everyday language use.

In this work, we re-analyzed five existing TMS datasets on semantic cognition to reveal how stimulation affected the processing of individual words (i.e. words in isolation, without any larger linguistic context). To ensure reliability of our results, we gathered as many publicly available datasets as possible. We could cover two languages (Italian and German), seven brain areas associated with semantic processing and five different semantic tasks, involving both comprehension and production tasks. The main characteristics of each dataset can be visualized in Figure 1. As a measure of cognitive processing related to processing of individual words, we used response times (RTs), that is the most commonly used index of cognitive effort in TMS studies (Devlin et al., 2003; Jefferies, 2013; Papeo et al., 2013; Qu et al., 2022).

**Figure 1.**
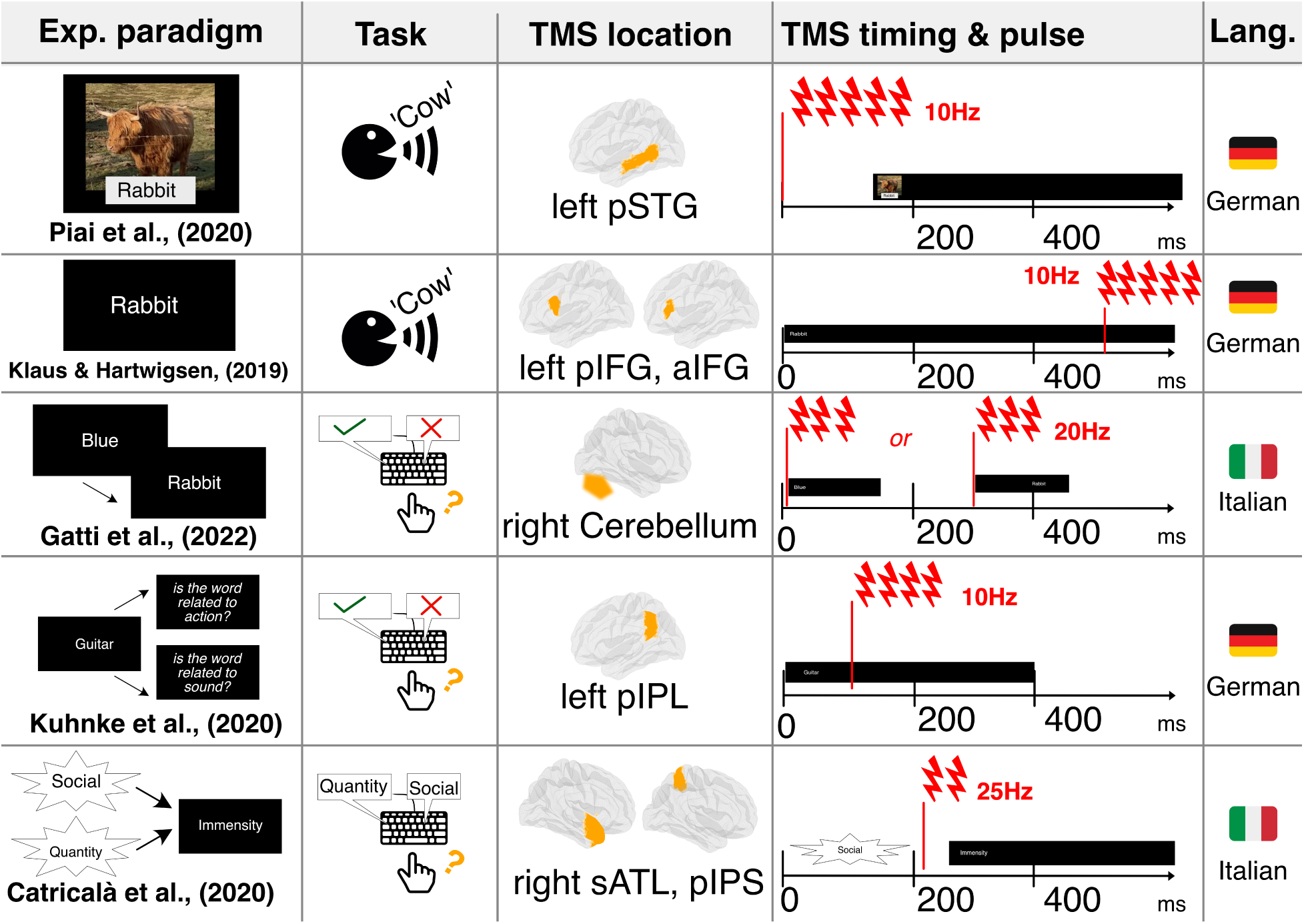
Main details of the used datasets.

To quantify the effects of individual word meaning on each trial, we adopted an information-based framework - that is commonplace in functional neuroimaging and electrophysiological approaches (Kriegeskorte & Bandettini, 2007; Naselaris et al., 2011) but not in TMS studies, at least for language (Romei et al., 2016). We measured the correlation between cognitive effort and different types of information (semantic, frequency and orthographic information), and probed whether this correlation was affected by TMS. We also used a simple procedure (hierarchical bootstrapping with subsampling - Saravanan et al. (2020)) to provide standardized measurements of correlation across studies and conditions, to even out differences in sample sizes that could bias the summary statistics (Minarik et al., 2016; Poldrack et al., 2017). The key advantages of this approach are twofold.

First, since it is based on correlation, it allows us to use continuous measurements for each type of information, thus recognizing and accounting for the specificity and uniqueness of the individual words in each trial (Bruffaerts et al., 2019; Gao et al., 2023).

Second, it operates at a more abstract level than simple RTs, which is by contrast commonplace in TMS research (Hartwigsen, 2015; Qu et al., 2022). A difference in RTs across conditions between effective and control stimulation is traditionally considered to be - in the context of a TMS study on semantic cognition - a measure of a semantic effect. However, such differences could in principle be explained by different factors (i.e. distinct types of information). By formulating explicit, competing models of cognitive processing, and evaluating how their fit with cognitive effort is affected by neurostimulation, an information-based approach enables to directly test whether the TMS effect on cognitive effort can be attributed to modulation of semantic information processing (Kriegeskorte & Douglas, 2018; Lopopolo et al., 2024; Naselaris & Kay, 2015; Peelen & Downing, 2023).

Importantly, this also allows to neutralize the effect of specific experimental parameters such as the stimulation timing, or the frequency and the number of pulses, that are known to have low-level effects on behavioural measures like RTs (Beynel et al., 2019; Hartwigsen & Silvanto, 2023; Pell et al., 2011).

For semantic information, we modelled the cognitive effort induced by each trial with a language model, or distributional semantic model (Boleda, 2020; Connell & Lynott, 2024; Lenci, 2018) - belonging to a family of computational models that have been shown to excel at capturing cognitive semantic processing (Goldstein et al., 2024; Huth et al., 2016; Lenci et al., 2022; Schrimpf et al., 2021; Tuckute et al., 2024). For each trial, we made the simple assumption that the cognitive effort should be related to the semantic similarity between the concepts involved in it - an expectation justified by a large body of previous work (Bhatia & Aka, 2022; Wingfield & Connell, 2022). We also implemented in the same way the two control models, capturing lower levels of processing: one using word frequency (i.e. statistical information), the other using word length (i.e. orthography).

Then, separately for each study, model and stimulation condition, we computed the correlation between RTs and each model (language model, word frequency, word length). Finally, we tested whether the effect of TMS on the fit between RTs and each type of information was significant: if that were the case, we interpreted as indicating that processing of that information was affected by stimulation. For a simplified visualization of our approach, see Figure 2A.

**Figure 2.**
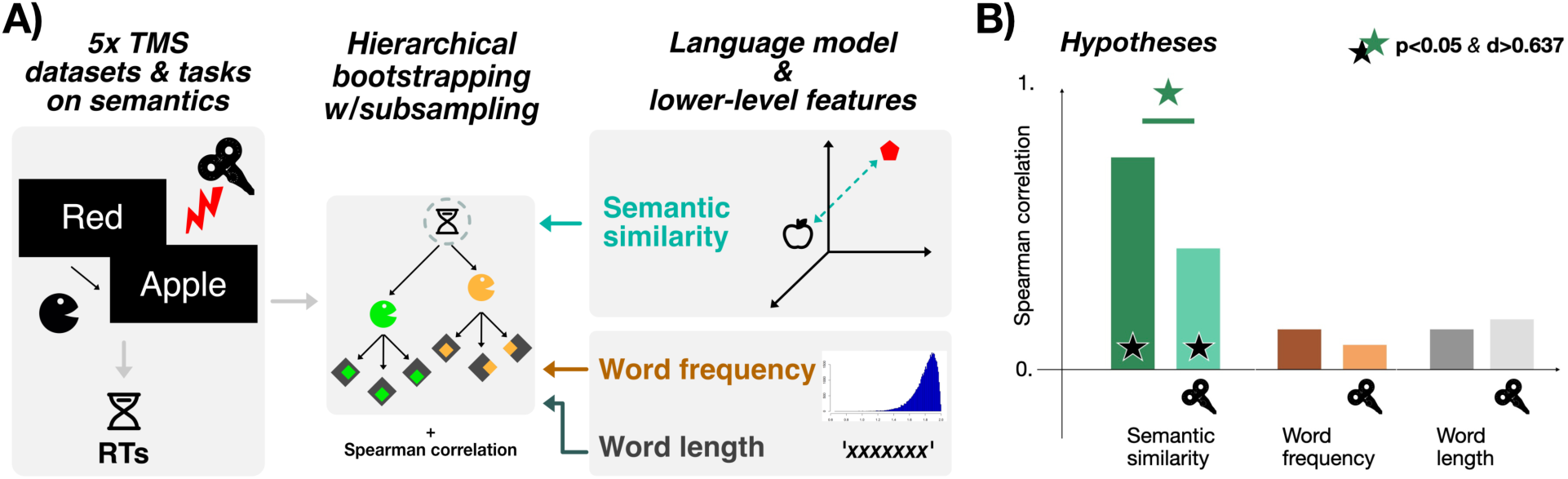
Methodology used to measure brain stimulation effects on semantic cognition (A), and experimental hypotheses that we expected to hold across studies (B).

Inspired by previous neurostimulation studies (e.g., Gatti et al. (2022)), we tested two hypotheses (see Figure 2B). First, the semantic similarity model should show above-chance correlation with cognitive effort for all tasks, in the condition where no effective stimulation was applied - indicating that individual word-level semantics was effectively captured in regular, unaltered cognition; regarding word frequency and length we had no specific expectation. Second, TMS should induce a significant modulation of the fit only for the semantic similarity model, indicating that the TMS effect was specific to semantics - and, that the directionality of this effect should be a reduction in correlation after stimulation (see Gatti et al. (2022)).

## Results

For simplicity, we divided the presentation of the results in four sections; first reporting the two datasets on language production (Klaus & Hartwigsen, 2019; Piai et al., 2020); then the two datasets on semantic judgments without priming (Gatti et al., 2022; Kuhnke et al., 2020); and finally, the semantic priming study (Catricalà et al., 2020).

We added an additional set of analyses since, contrary to our expectations, the effect of TMS turned out to be significant also on non-semantic variables for some datasets. Therefore, we repeated the analyses after removing the variance that could be explained by them (both word length and word frequency) from the RTs. This allowed us to measure the specific effect of TMS on semantics.

We report the full set of statistics in the main text (Spearman correlation, p-value, effect size, 95% CIs) only if two conditions were fulfilled: a p-value smaller than 0.05 *and* an effect size larger than *d>* 0.632, which corresponds to the 25th percentile value estimated from significant results across the cognitive neuroscience literature - that we interpret as indicating a small effect. We report the full set of statistics in the Supplementary Materials (Additional Results section, Tables Note that if the correlation between a model and the RTs is not statistically significant and the data do not show at least a small effect in the control condition (sham/vertex), the comparisons between conditions are not interpretable because of the lack of reliability of the model’s ability to explain the behavioural data. In such cases, we do not report statistics in the main text.

Also, while in the text for simplicity we refer to semantic (dis)similarity, word frequency and word length, note that the actual implementation of the models’ measures was adapted so that the expected correlation with cognitive effort (indexed by RTs) would be positive: therefore, in practice, we computed the correlation between RTs and semantic dissimilarity (1-similarity; the less similar two words, the larger the cognitive effort); RTs and negative word frequency (the less frequent a word, the larger the cognitive effort); RTs and word length.

### TMS increases sensitivity to semantic (dis)similarity in semantic production tasks

We start from the picture-word interference task of Piai et al. (2020) (Figure 3, left). Semantic dissimilarity correlated significantly with RTs for both vertex (*ρ* = 0.257, *p* = 0.004, *d* = 6.16, 95%*CIs* = (0.255, 0.260) and left pSTG (*ρ* = 0.293, *p* = 0.004, *d* = 7.12, 95%*CIs* = (0.290, 0.296)), while this was not the case for word frequency (vertex: *ρ* = 0.0459, left pSTG: *ρ* = 0.0331) nor word length (vertex: *ρ* = 0.0588, left pSTG: *ρ* = 0.0384). Similarly, the effect of TMS was significant and showed at least a small effect only for semantic dissimilarity (*d* = *≠*0.85, 95%*CIs* = (*≠*0.93*, ≠*0.78), *p* = 0.003), while it had a negligible effect for word frequency, however with the opposite direction as semantic similarity (*d* = 0.288, *p* = 0.003). The same happened with word length (*d* = 0.471, *p* = 0.003). Contrary to our hypothesis, effective TMS increased the correlation between semantic dissimilarity and RTs - suggesting that after stimulation, the semantic (dis)similarity between the trial-relevant items plays a more important role in determining the amount of cognitive effort involved in solving the task.

**Figure 3.**
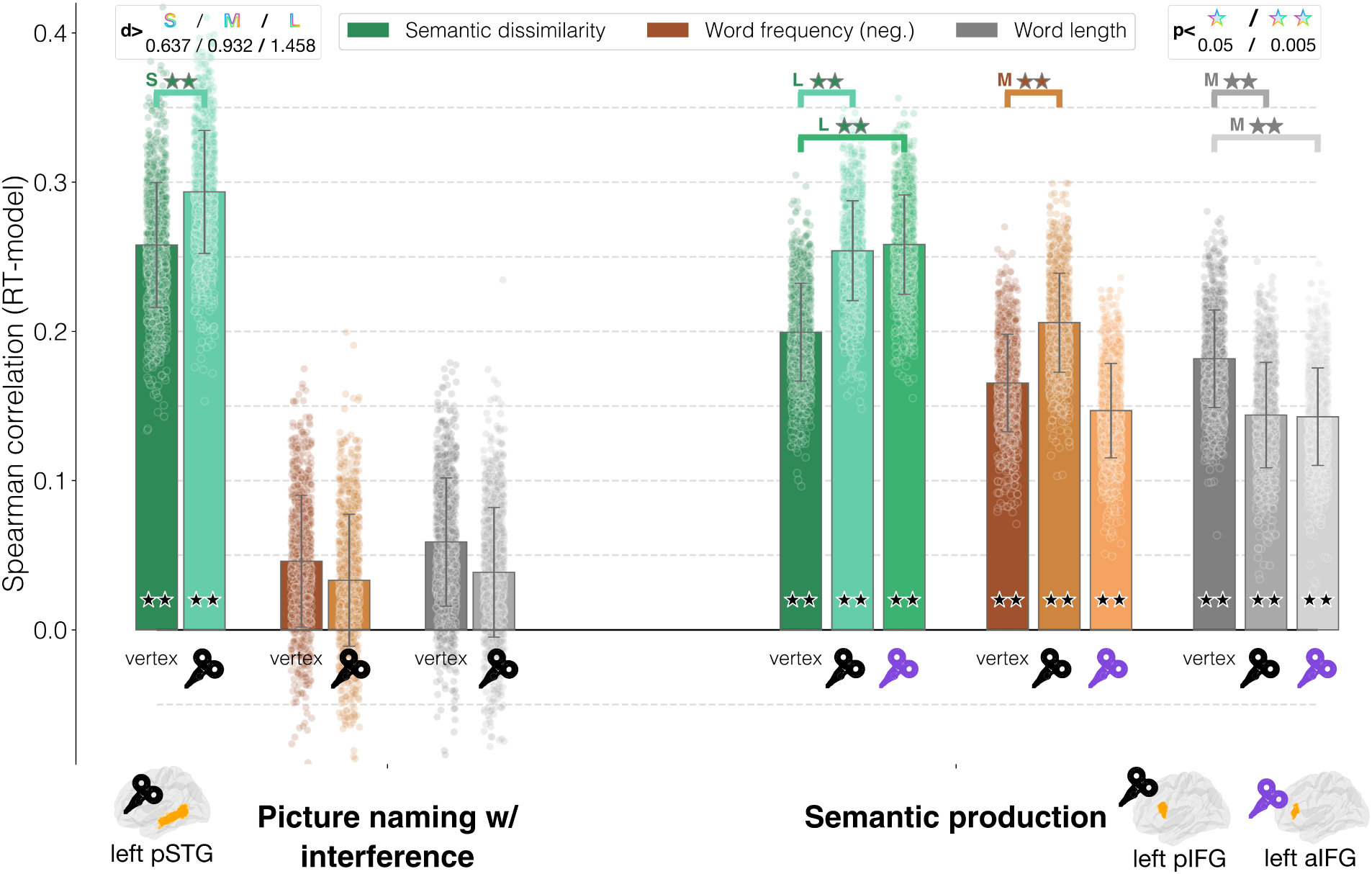
Semantic production tasks - main results. The bar heights correspond to the average Spearman correlation between each model and the RTs across all bootstrap iterations; error bars represent the standard deviation; the dots corresponds to the Spearman correlation for each individual iteration.

We found different results for the semantic production task (Klaus and Hartwigsen (2019); Figure 3, right). Here, all models - both semantic and non-semantic - was significantly correlated with RTs, in all conditions. The highest correlation value in the control condition was found for semantic dissimilarity (vertex: *ρ* = 0.199, *p* = 0.0043, *d* = 6.08, 95%*CIs* = (0.197, 0.201); pIFG: *ρ* = 0.254, *p* = 0.0043, *d* = 7.59, 95%*CIs* = (0.251, 0.256); aIFG: *ρ* = 0.258, *p* = 0.0043, *d* = 7.76, 95%*CIs* = (0.256, 0.26)).

The second highest correlation for vertex was found for word length (vertex: *ρ* = 0.181, *p* = 0.0043, *d* = 5.56, 95%*CIs* = (0.179, 0.183); pIFG: *ρ* = 0.143, *p* = 0.0043, *d* = 4.07, 95%*CIs* = (0.141, 0.146); aIFG: *ρ* = 0.142, *p* = 0.0043, *d* = 4.36, 95%*CIs* = (0.14, 0.144);); followed by word frequency (vertex: *ρ* = 0.165, *p* = 0.0043, *d* = 5.06, 95%*CIs* = (0.163, 0.167); pIFG: *ρ* = 0.205, *p* = 0.0043, *d* = 6.2, 95%*CIs* = (0.203, 0.207); aIFG: *ρ* = 0.146, *p* = 0.0043, *d* = 4.63, 95%*CIs* = (0.144, 0.148)). When looking at differences among conditions, all comparisons were statistically significant, showing at least a medium effect. However, the directionality of the effects varied noticeably, showing an interaction between stimulation and type of information being processed: For semantic dissimilarity, TMS consistently increased correlation between RTs and semantic similarity, coherently with the results reported above on picture-word interference (pIFG: *d* = *≠*1.647, 95%*CIs* = (*≠*1.74*, ≠*1.55), *p* = 0.0032; aIFG: *d* = *≠*1.777, 95%*CIs* = (*≠*1.88*, ≠*1.68), *p* = 0.0032). For word length, TMS had the opposite effect, always decreasing correlations - thus indicating that the same stimulation can affect the processing of different pieces of information in different ways (pIFG: *d* = 1.108, 95%*CIs* = (1.03, 1.19), *p* = 0.0032; aIFG: *d* = 1.186, 95%*CIs* = (1.11, 1.27), *p* = 0.0032). This was even more strongly evident for word frequency, where the effect diverged depending on the target area: for pIFG TMS only, correlation with word frequency increased (pIFG: *d* = *≠*1.22, 95%*CIs* = (*≠*1.31*, ≠*1.15), *p* = 0.0032), while results went in the other direction and were only close to having a small effect for the aIFG (aIFG: *d* = 0.574, 95%*CIs* = (0.51, 0.64), *p* = 0.01).

Overall, the results obtained for language production provided clear evidence for a significant effect on semantics. However, they contradicted our hypothesis that semantic (dis)similarity should be decreased as TMS should induce noise in the system. Rather, our results show that TMS increased the amount of variance explained by this value and this effect. Also, this was not specifically affecting semantic processing, but also lower level processes (recognition, as captured by word frequency, and orthography, *via*word length).

### Semantic judgments show that TMS and task interact

Results for semantic judgment tasks are shown in Figure 4. On the left side, we report correlations for the relatedness judgment task of Gatti et al. (2022). We confirmed the original study’s results, which also motivated some of our hypotheses. We found that semantic dissimilarity was significantly correlated with RTs (vertex: *ρ* = 0.214, *p* = 0.0044, *d* = 4.76, 95%*CIs* = (0.211, 0.216); right Cerebellum: *ρ* = 0.159, *p* = 0.0044, *d* = 3.65, 95%*CIs* = (0.157, 0.162)). Moreover, TMS significantly affected semantic processing, by reducing the correlation between semantics and RTs after stimulation (*d* = 1.184, 95%*CIs* = (1.1, 1.27), *p* = 0.003). Word length was significantly correlated with RTs in both conditions (vertex: *ρ* = 0.107, *p* = 0.0314, *d* = 2.58, 95%*CIs* = (0.104, 0.11); right Cerebellum: *ρ* = 0.0975, *p* = 0.042, *d* = 2.373, 95%*CIs* = (0.094, 0.1)); however, the effect of TMS on processing of orthographic information was negligible (*d* = 0.43, 95%*CIs* = (0.37, 0.5), *p* = 0.0032). Finally, correlation between word frequency and RTs was never statistically significant (vertex: *ρ* = 0.0745; right Cerebellum: *ρ* = 0.0838). In this respect, the results from this dataset fit our hypotheses: semantic (dis)similarity explained RTs during the task significantly better than chance, and such semantic information was selectively targeted by stimulation.

**Figure 4.**
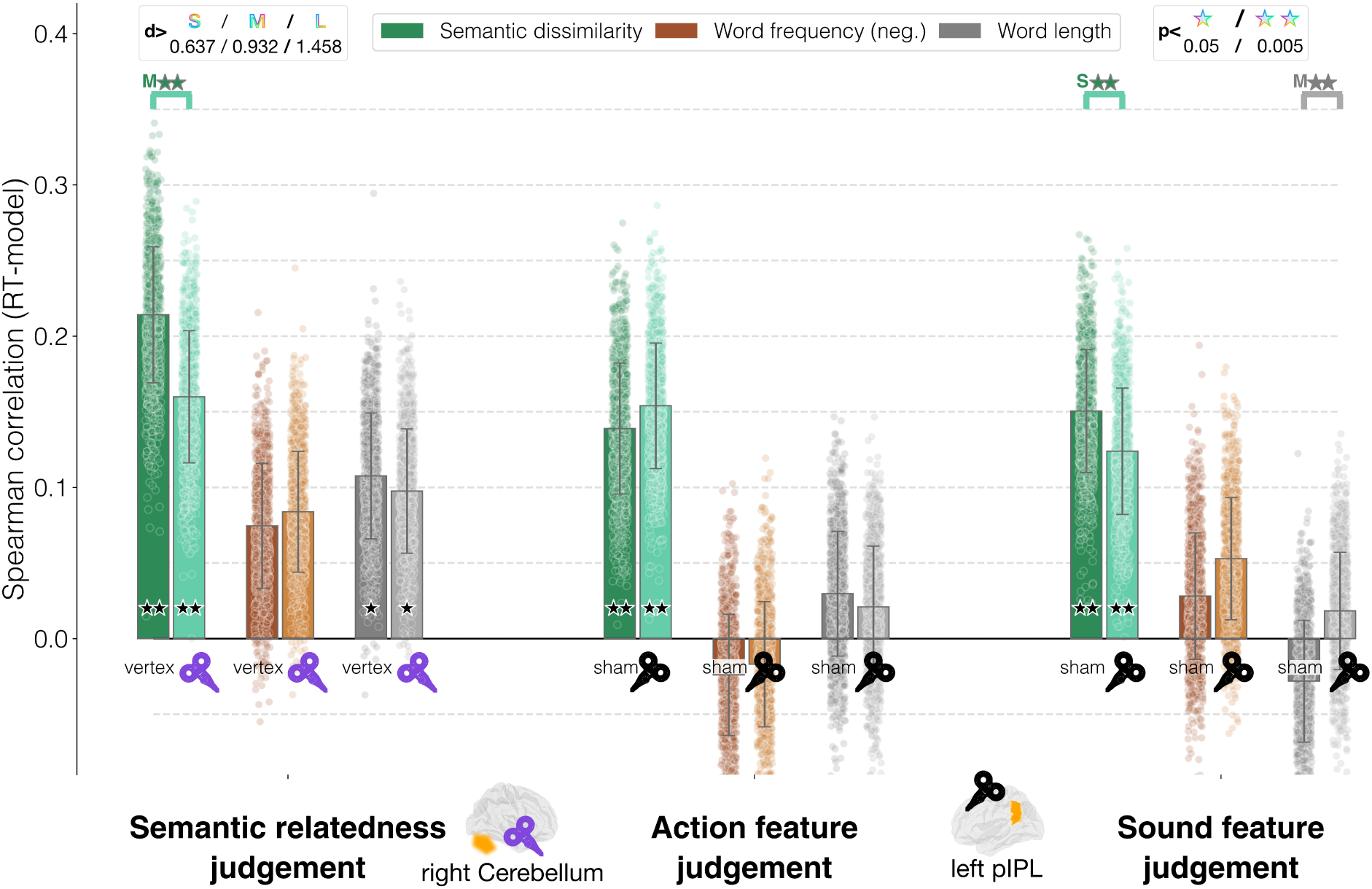
Semantic judgment tasks - main results. The bar heights correspond to the average Spearman correlation between each model and the RTs across all bootstrap iterations; error bars represent the standard deviation; the dots corresponds to the Spearman correlation for each individual iteration.

A similar picture emerged for the sound and action judgments dataset of Kuhnke et al. (2020), with the added value of showing a critical interaction between stimulation and task. More specifically, for the action judgment task (Figure 4, middle), only semantic dissimilarity was significantly correlated with RTs (sham: *ρ* = 0.138, *p* = 0.0044, *d* = 3.207, 95%*CIs* = (0.136, 0.141); left pIPL: *ρ* = 0.153, *p* = 0.0044, *d* = 3.706, 95%*CIs* = (0.151, 0.156)), and neither of the lower-level features did so (word frequency - sham: *ρ* = *≠*0.024; left pIPL: *ρ* = *≠*0.0169; word length - sham: *ρ* = 0.029; left pIPL: *ρ* = 0.021). However the effect of TMS on semantics during the action task, as measured by the effect size, was negligible (*d* = *≠*0.353, *p* = 0.003).

In contrast, for the sound task, the results matched the expected effects (Figure 5, right side): semantic dissimilarity was not only significantly correlated with RTs (sham: *ρ* = 0.15, *p* = 0.0044, *d* = 3.69, 95%*CIs* = (0.148, 0.153); left pIPL: *ρ* = 0.123, *p* = 0.0044, *d* = 2.96, 95%*CIs* = (0.121, 0.126)) but was also significantly affected by TMS, causing the expected decrease in correlation after stimulation (*d* = 0.644, 95%*CIs* = (0.58, 0.71), *p* = 0.003). Neither word frequency nor word length show significant correlations in the sham condition (word frequency: *ρ* = 0.028; word length: *ρ* = *≠*0.0282)

**Figure 5.**
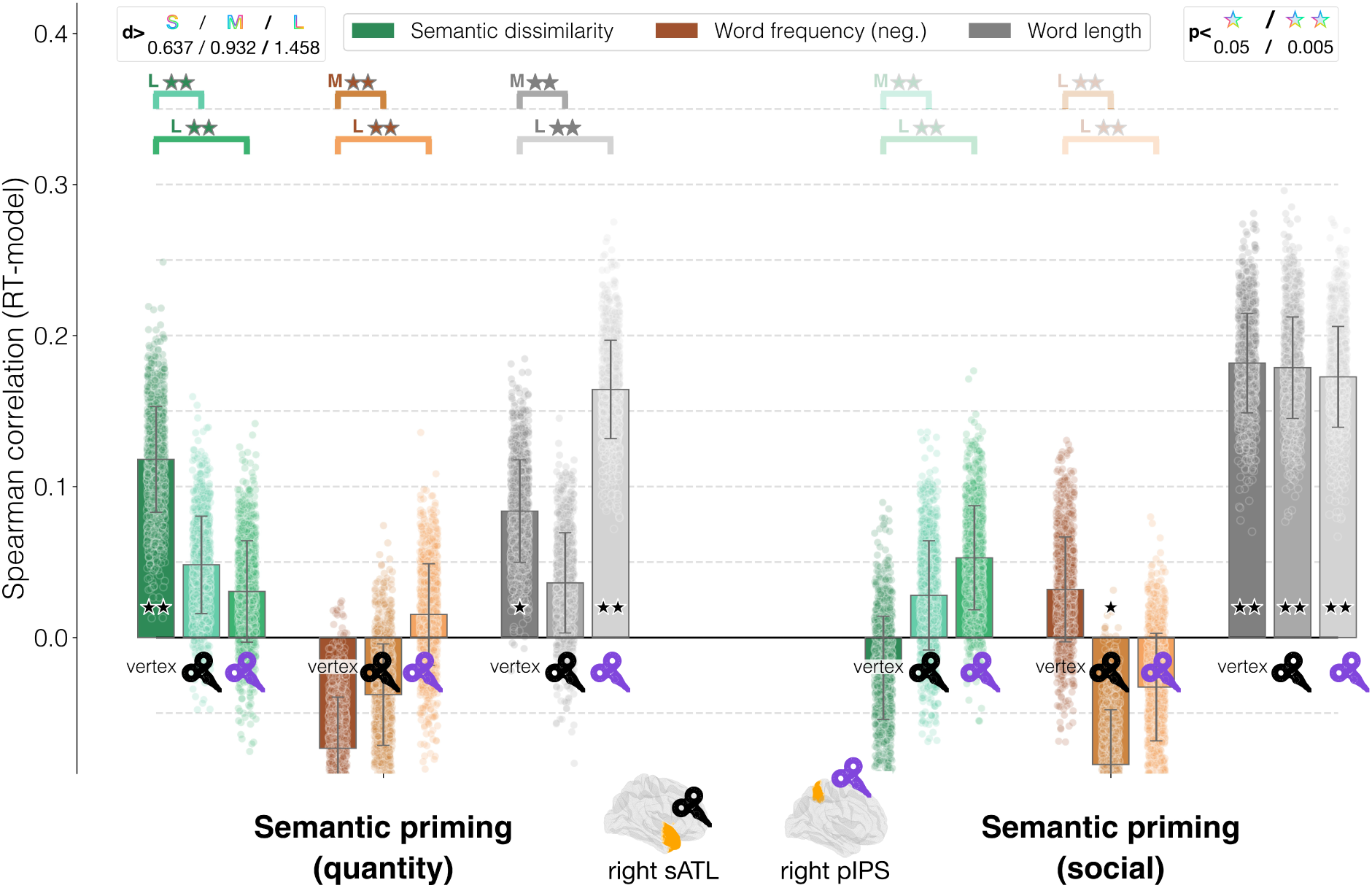
Semantic priming tasks - main results. The bar heights correspond to the average Spearman correlation between each model and the RTs across all bootstrap iterations; error bars represent the standard deviation; the dots corresponds to the Spearman correlation for each individual iteration.

### Language models capture TMS effects on semantic priming for quantity words

Finally, the results for the priming experiment are reported in Figure 5. Starting from semantic priming on quantity concepts (Figure 5, left side), semantic dissimilarity emerged as the closest match to RTs (vertex: *ρ* = 0.117, *p* = 0.0044, *d* = 3.36, 95%*CIs* = (0.115, 0.12)), followed by word length (vertex: *ρ* = 0.0837, *p* = 0.024, *d* = 2.46, 95%*CIs* = (0.0816, 0.085)). Word frequency, on the contrary, did not correlate significantly with RTs (vertex: *ρ* = *≠*0.0732). Stimulation had a strong effect on semantic processing, both for right pIPS and right sATL. In both cases, TMS decreased the match between semantic dissimilarity and RTs (right pIPS: *d* = 2.54, *p* = 0.0032, 95%*CIs* = (2.42, 2.68); right sATL: *d* = 2.073, *p* = 0.0032, 95%*CIs* = (1.96, 2.18)). This is consistent with the results reported in Figure 4 - in all these datasets the experimental task required subjects to provide semantic judgments. For word length, again consistently with the results of Figure 4, the effect of TMS depended on the target: when stimulating the right sATL, the correlation was significantly decreased while for the right pIPS, orthographic information was significantly correlated with RTs (right pIPS: *d* = *≠*2.419, *p* = 0.0032, 95%*CIs* = (*≠*2.54*, ≠*2.3); right sATL: *d* = 1.41, *p* = 0.0032, 95%*CIs* = (1.33, 1.5)). This aligned only partially with our hypotheses: the TMS effect on semantics matched our expectations, and revealed that not only the right pIPS, but also the right sATL played a role in semantic processing of quantity words. However, in contrast to our expectations, orthographic information was also significantly affected by TMS, although in partially different directions.

For social concepts (Figure 5, right side), our analyses revealed a different picture: variation in RTs seemed to be largely due to word length, the only model that was significantly correlated with RTs (vertex: *ρ* = 0.181, *p* = 0.0044, *d* = 5.52, 95%*CIs* = (0.179, 0.183); right pIPS: *ρ* = 0.172, *p* = 0.0044, *d* = 5.17, 95%*CIs* = (0.17, 0.174); right sATL: *ρ* = 0.178, *p* = 0.0044, *d* = 5.3, 95%*CIs* = (0.176, 0.18)). However, TMS did not seem to have an effect on orthographic processing either (right pIPS: *d* = 0.275, *p* = 0.063, 95%*CIs* = (0.21, 0.34); right sATL: *d* = 0.0911, *p* = 0.336, 95%*CIs* = (0.03, 0.15)); we interpret this as suggesting that none of the models used was in fact able to capture the type of processing affected by TMS.

### TMS effects on semantic processing are still present after accounting for word length and frequency

In a final set of analyses, we tested whether semantic (dis)similarity could still explain a significant portion of variance in RTs after removing the variance explained by lower-level variables (word frequency and worth length). In this way, we probed whether the effect of TMS on semantic processing was specific to semantics, and not confounded by recognition or orthographic effects (see Figure 6).

**Figure 6.**
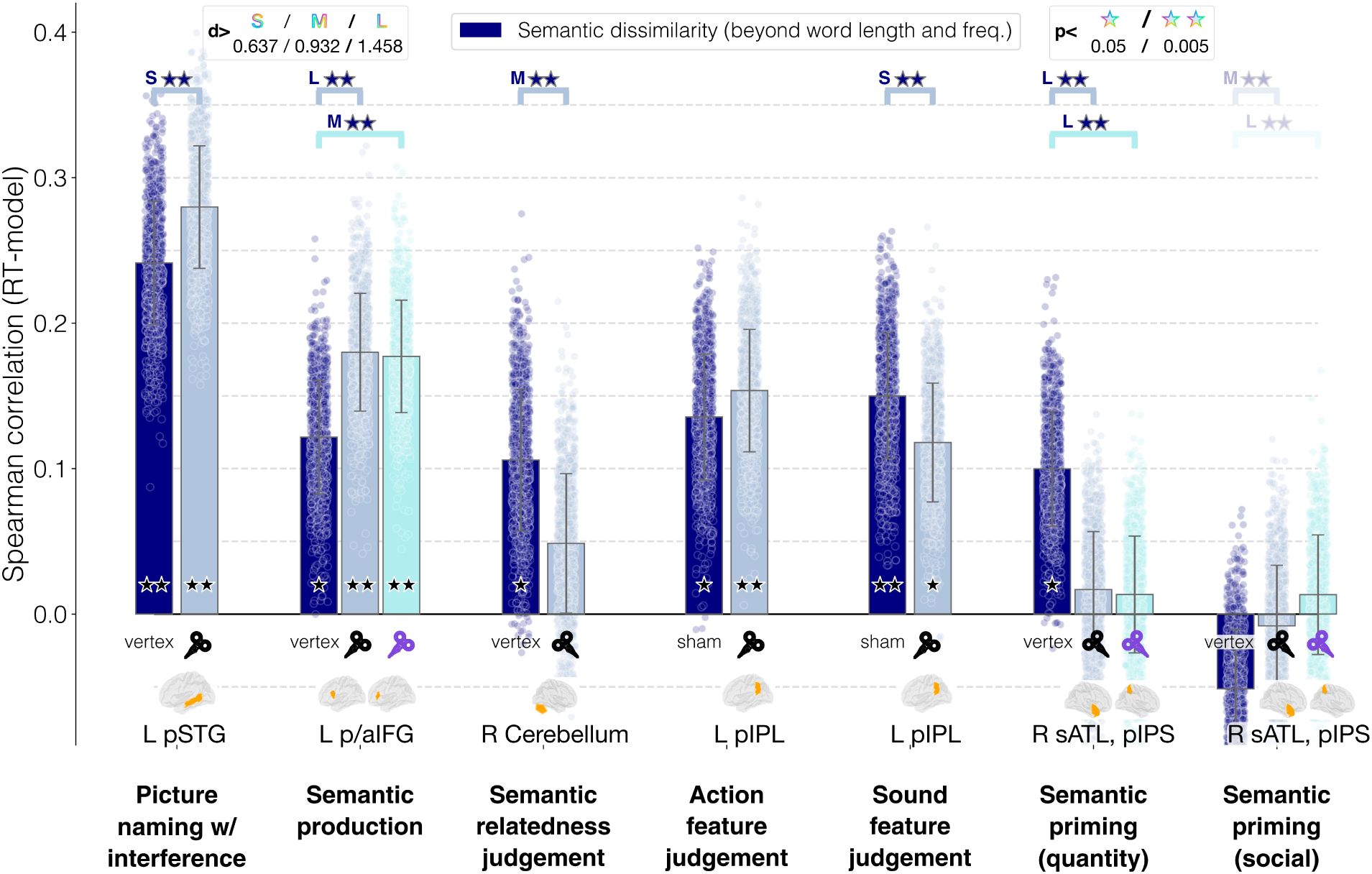
Semantic similarity results after residualization of word length and frequency. The bar heights correspond to the average Spearman correlation between PPMI dissimilarity and RTs across all bootstrap iterations; error bars represent the standard deviation; the dots corresponds to the Spearman correlation for each individual iteration.

To remove the variance explainable by word frequency and length alone, we adapted a cross-validated confound removal procedure validated by Snoek et al. (2019). For each subject within every iteration of the hierarchical bootstrap subsampling procedure, we trained a linear regression model on the left-out trials, learning to predict the RTs from word length and log-transformed word frequency. Then, we predicted the RTs selected for the current subsampling iteration using the newly trained model, and transformed the original RTs to the residual errors (i.e. the difference between the predicted and the true RTs; also called ‘residualization’ Rao et al. (2017)). This procedure ensures an unbiased removal of confounding information - due to training the residualization model with a dedicated, left-out portion of the data (Snoek et al., 2019).

We report here the results only for the semantic similarity model; in the Supplementary Materials (section Additional Results) it is possible to see the results also for word length and word frequency.

As expected, correlation values numerically decreased, which is likely explained by the known correlation between word length and word frequency (Piantadosi et al., 2011) as well as word frequency and semantic similarity (Burdick et al., 2018; Zhou et al., 2022).

Nevertheless, all the results reported above were confirmed. For language production, semantic dissimilarity explained best RTs for word naming with interference (vertex: *ρ* = 0.241, *p* = 0.0044, *d* = 5.69, 95%*CIs* = (0.238, 0.243)), and TMS significantly increased the way in which this type of semantic information was used, with a small effect size (*d* = *≠*0.909, *p* = 0.0032, 95%*CIs* = (*≠*0.98*, ≠*0.84)). For semantic production, we replicated the original pattern of results reported above - correlations were significantly above chance in all cases (vertex: *ρ* = 0.121, *p* = 0.0085, *d* = 3.105, 95%*CIs* = (0.119, 0.124)), and TMS again increased the correlation between semantic dissimilarity and RTs (pIFG: *d* = *≠*1.464, *p* = 0.0032, 95%*CIs* = (*≠*1.55*, ≠*1.38); aIFG: *d* = *≠*1.425, *p* = 0.0032, 95%*CIs* = (*≠*1.51*, ≠*1.34)).

For semantic judgment tasks too, the semantic effect of TMS on RTs was left untouched by residualization, consistently decreasing correlation between semantic dissimilarity and RTs. After removing the variance explained by lower-level variables, correlation between semantic dissimilarity and RTs in the semantic relatedness judgment task was significantly above chance (vertex: *ρ* = 0.105, *p* = 0.05, *d* = 2.17, 95%*CIs* = (0.102, 0.108)), as was the difference between effective stimulation of the cerebellum and vertex (*d* = 1.18, *p* = 0.0032, 95%*CIs* = (1.1, 1.27)). For the sound and action judgment dataset and the priming study, results followed the same pattern as above - TMS had a slightly detrimental effect on correlations between RTs and semantic similarities for sound feature judgments (vertex: *ρ* = 0.149, *p* = 0.0043, *d* = 3.475, 95%*CIs* = (0.147, 0.152)) and semantic priming with quantity concept words (vertex: *ρ* = 0.099, *p* = 0.031, *d* = 2.52, 95%*CIs* = (0.097, 0.102)), but the semantic effects remained well above significance (sound feature judgment: *d* = 0.762, *p* = 0.0032, 95%*CIs* = (0.69, 0.83); quantity priming, right pIPS: *d* = 2.16, *p* = 0.0032, 95%*CIs* = (2.06, 2.28); quantity priming, right sATL: *d* = 2.08, *p* = 0.0032, 95%*CIs* = (1.98, 2.2)). As before residualization, semantic effects on TMS for action feature judgment did not reach significance (*d* = *≠*0.425, *p* = 0.0032), and for social priming semantic similarity did not show significant correlation with RTs (*ρ* = *≠*0.051) .

## Discussion

In this study, we show that language models provide new insight into TMS-induced modulations of semantic cognition during word production and comprehension.

Our main finding was that semantic similarity could be used to successfully capture how RTs are modulated by non-invasive brain stimulation in trial- and word- specific semantic processing. We found that TMS affected semantic processing at the level of individual words, as measured with the language model, across almost all areas, tasks and datasets, with only minor exceptions (again social concept priming, and action feature judgment). Overall, these findings validate our approach. Importantly, they confirm that language models, whose ability to predict brain responses to language is by now well established, can be used to capture the subtle variations in behavioural measures induced by TMS - opening up the possibility of future experiments where word-specific modulations are exploited together with non-invasive brain stimulation.

However, importantly, two findings went against our predictions - highlighting the inherent challenges when looking at such fine-grained information using TMS. First, a significant stimulation effect was found to affect also lower-level, non-semantic processing for at least two tasks (semantic production and quantity semantic priming). This indicates that even if the effects of TMS are intended to operate on one single, cognitive level (semantics), they can in fact spread to other levels of processing. We note that similar effects have been demonstrated at the cortical level: despite stimulation targeting a single area, its effects can be widespread on the cortex (Beynel et al., 2020; Numssen et al., 2023; Schuler & Hartwigsen, 2024). Such converging evidence confirms the need for going beyond simple comparisons of RTs across binary conditions. What is needed is the comparison of explicit competing models, capturing trial-by-trial variation in the stimuli, whose relative ability to explain the RTs can be assessed and interpreted in the context of general theoretical models of linguistic and semantic processing (Kriegeskorte & Douglas, 2018; Lopopolo et al., 2024; Naselaris & Kay, 2015; Peelen & Downing, 2023). Secondly, semantic priming involving social semantic concepts could not be captured by semantic similarity; this may, however, be due to limitations in the model (see below for a discussion of this point).

Based on our results, one might ask whether the effect on semantic processing were in fact determined by lower-level variables, instead of semantic information. To test that this was not the case, we ran a post-hoc analysis, removing the variance in the RTs that could be explained by word frequency and length before looking at correlations between RTs and the semantic model. The final results confirmed the significance of all the TMS effects on semantic processing that we found in the original analyses, beyond what could be explained by lower-level variables. The only difference to report is an overall numerical decrease in correlation values - which was expected since during language processing, variables at different levels tend to be correlated with each other (Burdick et al., 2018; Piantadosi et al., 2011; Zhou et al., 2022).

The second finding that went against our expectations was that effective stimulation did not always decrease the correlation between RTs and the semantic model. This finding was consistent only for language comprehension (semantic judgments) tasks. The opposite pattern (increase in correlations between the semantic model and RTs after stimulation) emerged for language production tasks, where control regions were stimulated. This implies that, during or after stimulation, the processing of a trial became more or less cognitively demanding, in proportion to the semantic similarity of its items.

While we acknowledge that it is impossible to solve this puzzle in the context of the current paper, we interpret our findings in the light of existing neural noise frameworks for TMS effects (Miniussi et al., 2013). We propose that this mechanism could be hypothesized to reflect the two sides of a possible TMS effect. TMS might either result in a sharpening of the stimulus sensitivity by adding an “optimal level of noise for task performance” (see Bergmann and Hartwigsen (2021)) and thereby increasing the correlation between RTs and a type of information. Alternatively, the TMS-induced noise may be detrimental, resulting in a desensitization to stimulus properties and a decrease in correlation. This is particularly relevant for future studies. It does not only provide a testable hypothesis regarding the functional effects of TMS on complex cognitive processes like language, but also holds the promise of shedding light onto the mechanistic neural effects of the causal links behind semantic cognition.

In the following paragraphs, we discuss our findings more in depth in light of our hypotheses and previous studies.

### Hypothesis 1: semantic similarity explains RTs in almost all behavioural tasks

Our results indicate that - with the only exception of semantic priming with social concepts - semantic (dis)similarity as computed from a language model is capable of reliably capturing the cognitive effort associated with different semantic tasks, as measured in reaction times. This aligns with previous studies on semantic priming (Jones et al., 2006) and semantic relatedness judgments (Gatti et al., 2022). Nevertheless, our results move beyond theses initial findings and provide important new insight into semantic processing.

First, our results generalize to a much wider range of tasks where reaction times are used to capture semantic processing, spanning both language comprehension and production. Previous studies largely focused on comparing semantic similarity measures from language models to explicit word ratings (e.g. the model-based similarity vs the average human similarity rating for the pair of words ‘concepts’ and ‘object’ – Banks et al. (2021), Connell and Ramscar (2001), Lenci et al. (2022), and Wingfield and Connell (2022)). Here, by contrast, we show that language models are able to go one step further - significantly explaining trial-by-trial response times, a measure of implicit cognitive effort (Baayen & Milin, 2010), across a range of different tasks.

Moreover, we analyzed various datasets with an approach that strictly matched the final computation of statistics, ensuring a straightforward side-by-side interpretation of the general results. We adopted a bootstrap approach based on subsampling, providing a standardized evaluation of the amount of variance that could be explained by semantic similarity for each task. Such standardized scores are useful for planning future TMS studies on semantic processing as they offer a set of testable hypotheses with respect to the effects of TMS on semantic processing - both in terms of expected baseline values, as well as directionality of the effect (e.g. increase/decrease with stimulation at a certain site).

Our results allow researchers to choose a task whose cognitive effort can be successfully captured by semantic similarity. We found that, overall, RTs in production tasks (picture-word interference, semantic production) and at least some semantic judgments (semantic relatedness, sound features) are solidly captured by a semantic similarity measure. Priming, by contrast, seemed to be slightly trickier to explain using semantic similarity; however, there may be dataset-specific reasons for this pattern of results.

In particular, the only task where the correlation between language models and RTs failed to exceed the chance level was semantic priming with social concepts. Social semantic information is unique - partly because it seems to be, to some extent, segregated from general semantic knowledge (Binney & Ramsey, 2020; Olson et al., 2013), and partly because it tends to vary strongly across individuals depending on life experiences (Mazzuca et al., 2020). Consequently, it is possible that language models - whose core principle, when it comes to lexical semantics, is abstraction over many contexts of occurrence of the same word - may fail, in the absence of subject-specific information, to capture social semantic knowledge (for recent attempts at overcoming such limitations, see Bruera and Poesio (2024) and Johns (2021)).

### Hypothesis 2a: TMS effects on semantic processing are reliably found across studies, but their directionality is an open question

We found that TMS significantly modulated the amount of variance in RTs that could be explained by semantic (dis)similarity as computed from a language model. Since language models capture a broad, generic but deep knowledge about word meaning, we interpret this as indication that TMS causally affects the way the brain makes use of semantic knowledge. Importantly, we also show that this effect went beyond what could be explained by other lower-level, non-semantic variables.

This was true for all datasets, although not for all tasks within each dataset. Namely, action feature judgments and social concept priming did not show a significant effect of stimulation on semantic processing. Regarding the latter, semantic similarity – as computed with a language model – did not seem to explain RTs at all, (see previous section for a discussion). For action feature judgments, two possible explanations should be taken into account. First, the absence of a significant modulation may be explained by the way subjects ‘solve’ the task - hence, with the way we modelled it: semantic similarity may be a good model only for one feature (sound, where we found a significant effect), but not the other (action). Alternatively, the targeted pIPL may differentially retrieve and use sensorimotor relative to auditory information – which would reflect different processing strategies for different semantic modalities. The current analyses do not allow for a distinction between the two explanations, since we simply re-analyzed the dataset, and could not design ad-hoc experimental manipulations to decide between the two. Nevertheless, a clear picture can be drawn at a more general level: there is an interplay between TMS effects on semantics and the individual tasks.

A more pressing open issue is the question why TMS increased the fit between language model dissimilarity and response times for some tasks and decreased it for others.

A tempting explanation would be to interpret such results in light of the inhibition/excitation (also facilitation) dichotomy that is routinely used to frame TMS effects, but this has also been strongly criticized (Hartwigsen & Silvanto, 2023; Hussain & Freedberg, 2024). From this perspective, a higher correlation between RTs and semantic similarity could indicate facilitation of semantic information in the stimulated brain areas; a lower correlation would indicate, by contrast, inhibition, potentially due to an increase in noise.

However, such an interpretation is not warranted given our current knowledge of TMS effects on non-motor areas (Hussain & Freedberg, 2024). Previous work has emphasized the state dependency of TMS effects, demonstrating that the same protocol can have differential effects depending on the individual brain state (e.g., task versus rest) or level of baseline performance - (see Bradley et al. (2022) and Silvanto et al. (2007)).

Secondly, it is not clear whether the effect of TMS on the stimulated neurons is best described as noise injection, temporary silencing (a so-called ‘virtual lesion’), or something else (Ruzzoli et al., 2010). An additional issue that is relevant here is that the vast majority of studies investigating how TMS works at the level of neurons have looked at the motor cortex only. Because of the complexity of the cortex, it is not obvious that other areas will respond in the same way as the motor ones (Siebner et al., 2022). With the current analyses and without any neuroimaging data, we cannot distinguish the different possible explanations.

Nevertheless, our results show that, as expected, all semantics-related areas that were stimulated were also affected by TMS as compared to their non-stimulated state. Some of the observed differences in our results may reflect the differential role of the targeted areas in semantic processing. Stimulation of areas associated with semantic control, the left IFG (Jackson, 2021) and the left pSTG (Piai et al., 2020), increased the correlation between RTs and semantic similarity; while stimulating areas not involved in semantic control but representation or integration (left pIPL, right Cerebellum, right ATL, right pIPS) resulted in decreased correlations. A second option could be that stimulation during semantic production tasks (incidentally involving the IFG and the pSTG) has a different effect than stimulation during semantic comprehension. Differentiating between these explanations would require a controlled study that should keep all other parameters (e.g., stimulation protocol, timing, brain region) constant.

Nevertheless, our computational modeling approach complements the focus on brain areas with a discussion at the level of information processing. While such a perspective is not common in TMS studies yet (notable exceptions being reviewed in Miniussi et al. (2013)), it is well established in neuroimaging and electrophysiological studies (Kriegeskorte & Bandettini, 2007; Naselaris et al., 2011). In this sense, the question is not so much whether TMS helped or impaired semantic processing. Rather, we ask whether TMS triggered, at a computational level, a process that increased (higher correlation with RTs) or decreased (lower correlation with RTs) the importance of semantic (or statistical, or orthographic) information in determining RTs.

Under such a framework, the three main observations from our analyses were as follows: first, the TMS effect on semantic information processing was statistically significant. Secondly, the modulatory effect interacted with individual word meaning, as captured by language models. Third, it could be isolated from other levels of linguistic processing. The increase of correlation between semantic similarity and RTs after TMS could be reformulated as follows: the easier the items were to process originally, the easier they will become after TMS; and vice versa for harder items, that become even harder as a consequence of TMS. It is evident that this is not compatible with an inhibitory/facilitatory account of TMS: this implies that TMS both helps (in the case of easier trials) and disrupts (in the case of tougher trials) processing, depending on the interaction with processing difficulty.

Another mechanistic explanation would be that TMS increased or decreased the stimulus-sensitivity of the cognitive processing at hand. In some cases (semantic production, when stimulating semantic control areas) TMS increased *stimulus-sensitivity* during semantic processing - sharpening the effect of semantic similarity.In other cases, by contrast, TMS made cognitive processing more generic, less affected by the semantic representations of the words. In the larger context of how exactly TMS affects information processing, this can be related to the existing literature on so-called discrimination sensitivity (Miniussi et al., 2013). In short, the relationship between the input sensory strength and the sensitivity in behavioural responses (e.g. in an auditory discrimination task) can be modelled by a sigmoid function, which is s-shaped: lower strengths are compressed to 0 (i.e. not discriminated), and higher input strength is compressed to 1 (i.e. easily discriminable). Accordingly Miniussi et al. (2013) propose that TMS *shifts* the sigmoid function along the x axis, making it easier or tougher to discriminate at a fixed input sensory strength. We propose that in our tasks, where discrimination is not really involved, the sigmoid function may not have been shifted by TMS but *its slope may have been altered* (Han & Moraga, 1995). The sigmoid function thus becomes either steeper in its central part (increasing stimulus-sensitivity, hence correlation) or, on the contrary, more horizontal (decreasing stimulus-sensitivity and correlation). Again, while it’s impossible to validate whether this explanation is correct, it can motivate future studies, and allows to place the effects found here in the broader context of mechanistic models of brain stimulation.

### Hypothesis 2b: TMS effects on semantics-related brain areas can spread to lower-level cognitive processes

A final, important, result of our analyses was unexpected. We reasoned that TMS effects should be confined to semantic processing since we included studies with semantic tasks and target areas associated with semantic processing (rather than lower-level processes). Yet, we found that in some cases, TMS significantly affected processing of both word frequency and length.

For word frequency, this was the case only for the left IFG (see Figure 3). This is actually not surprising, since the IFG has been found to be sensitive to word frequency (e.g. Hauk et al. (2008) and Sánchez et al. (2023)). However, it shows that it may be premature to interpret effects on RTs as being exclusively semantic in nature. Looking deeper, however, the pattern of results could actually provide original insight with respect to the respective roles of the anterior and posterior portions of the left IFG, since they were both stimulated separately during a semantic production task.

Interestingly, the two show a clear dissociation. When looking at the cumulative effect on processing of word frequency (Figure 3), stimulation over the posterior IFG significantly increased correlations, while the opposite happened for the anterior part. When distinguishing between written and uttered words, the effect seems to be confined to production (see Supplementary Materials, Figures C3, C4). Consequently, we may conclude that stimulation of the left posterior IFG increased stimulus sensitivity for word frequency. This not only confirms its role in phonological processing, but also contributes to the discussion on its functional role (cf. Fiebach et al. (2002), Klaus and Hartwigsen (2019), and Sánchez et al. (2023)).

For word length, an even lower-level variable, we found that TMS unexpectedly had an effect for the left IFG, right sATL and right pIPS. In the left IFG, the effect was consistent across but subregions, resulting in correlation decreases between RTs and word length. Again, the effect was present when either considering both the written and the uttered words’ lengths together, or the uttered word in isolation, but not when considering only the length of the written word (Supplementary Materials, Figures C3, C4). This confirms the role of the left IFG at lower linguistic levels like orthography and phonology (Indefrey, 2011; Pammer et al., 2004). Also, this effect is interestingly opposed in directionality to that for word frequency: the same stimulation can increase stimulus sensitivity for one piece of information, and decrease it for another. In this sense, tracking the interactions between stimulation and computational processes can provide new insight for cognitive models of information processes during language processing (de Zubicaray, Regarding the right sATL and the right pIPS, the effect of stimulation on processing of word length was significant only for quantity concept priming. It was found to act especially on the target word (Supplementary Materials, Figures C14), and it acted in different directions across the two areas (decrease in correlation between RTs and word length for the sATL, increase for the pIPS). This is relatively surprising. Both areas are usually associated with higher-cognitive functions (semantic, non-linguistic processing for the right ATL (Lambon Ralph et al., 2017; Rice et al., 2018) and numerosity and spatial processing for the right IPS (Binkofski et al., 2016; Dehaene et al., 2005)) and only recent reports suggest they may both play a role in word processing (for the right ATL: Cope et al. (2020), for the right IPS: Santacroce et al. (2024)).

Finally, the discrepancy in results across types of priming remains puzzling: one would assume that TMS should have equally affected the processing of word length, given that they were part of the same experiment. While we are not able to explain this effect, we hypothesize that this is an after-effect of our finding that semantic similarity significantly explains priming RTs only for quantity (where also word length is significantly affected by TMS) but not for social concepts: it is possible that participants used different strategies for the two, and this significantly altered the way in which information was processed across the right pIPS and the right sATL, explaining the difference in the way TMS affected them.

In short, the above discussed results suggest that lower-level and higher-level processing interact and modulatory TMS effects may not be constrained to either level, even when higher-level areas are targeted. This is not surprising: language processing in the brain is a complex mesh of interrelated processes happening concurrently, interacting with each other and where information flows in multiple directions at the same time (Balota et al., 2004; Christiansen & Chater, 2008; Hickok & Poeppel, 2007; Levelt, 1999).

In summary, our approach shows that framing the modulatory effects of TMS on semantic processing in terms of information processing - by comparing the fit between competing models and cognitive processing - can unveil linguistic processes affected by TMS at different levels. This has not only the potential to advance our understanding of the mechanistic impact of non-invasive brain stimulation (Hartwigsen & Silvanto, 2023), but also to sharpen the definition of current models of language processing.

## Materials and Methods

### Datasets

Included datasets were chosen based on the following five criteria: 1. TMS study with a language task ; 2. including at least one TMS and one sham or other control condition; 3. a focus on lexical semantics (i.e. the meaning of individual words, not words in context); 4. reporting a significant stimulation effect on semantic processing; and 5. publicly and fully available dataset according to the FAIR Open Data principles (Wilkinson et al., 2016). First, we screened the publications of a recent review (Qu et al., 2022), identifying 3 papers that gave full public access to the datasets (Klaus & Hartwigsen, 2019; Kuhnke et al., 2020; Piai et al., 2020). Then, we expanded our search to Google Scholar, looking for papers respecting our criteria and containing one of the following combinations: ‘TMS’+’semantic’, ‘brain stimulation’+’semantic’, ‘TMS’+‘concepts’, ‘brain stimulation’+‘concepts’. We found two additional papers (Catricalà et al., 2020; Gatti et al., 2022). In the following, we provide short descriptions of the main characteristics of each dataset; a comparative visual overview of the is also given in Figure 1 (for detailed information, please refer to the subsection “Datasets” of the Supplementary Materials).

The first dataset employed as a task picture naming with interference (Piai et al., 2020). TMS was applied on left mid-to-posterior superior temporal gyrus (pSTG), an area reported to play an important role in semantic control, and particularly in resolving semantic interference (i.e. correctly naming a picture despite the presence of a distracting word; Jackson (2021)). Stimulation was online, consisting of burst of five pulses at 10 Hz delivered at each picture onset. TMS intensity was set to 90% of the individual motor threshold of the left primary motor hand area. The authors reported a significant facilitatory effect on RTs (i.e. lower RTs for TMS vs vertex) for a subset of the overall trials (cases where task difficulty was higher, and both the picture and the word referred to the same concept).

The second dataset was described in Klaus and Hartwigsen (2019). The experiment consisted of two production tasks - one phonological and one semantic; given our focus, we only analyzed the RTs for the semantic one. TMS was applied at two sites: the posterior and anterior portions of the inferior frontal gyrus (pIFG and aIFG respectively). Such regions were chosen because of their putative selective engagement for, respectively, phonological and semantic processing (Poldrack et al., 1999). TMS intensity was set to 90% of the motor threshold for the right hand region of the primary motor cortex. The authors reported an inhibitory effect on semantic processing when stimulating the aIFG (i.e. higher RTs for TMS vs vertex).

The third dataset was first analyzed in Gatti et al. (2022). The task was a semantic relatedness judgment between a noun and an adjective. Online 20Hz triple-pulse TMS was applied either at the noun or adjective onset, in a counterbalanced fashion. The stimulated area was the right cerebellum, that has started to be recently associated with semantic processing (Murdoch, 2010). Results revealed that the TMS effect shifted from facilitation to inhibition in semantic processing as the similarity between nouns and adjectives increased.

The fourth dataset was first presented in Kuhnke et al., 2020. There were two different, but structurally identical semantic feature judgment tasks: auditory feature and action feature judgment (i.e. ‘is the following concept related to sound?’, ‘is the following concept related to action?’). TMS was delivered as four online 10Hz pulses 100ms after word onset. The brain area of interest was the posterior inferior parietal lobe (pIPL), found to be consistently activated by a large number of semantic processing tasks (Jackson, 2021). The authors found a significant effect on accuracy (importantly, not on RTs) for the action feature judgment task alone.

The final dataset was introduced in Catricalà et al. (2020). It tested two types of semantic priming - using primes referring to either social (i.e. ‘sociability’) or quantity (i.e. ‘immensity’) concepts. There were two stimulation sites of interest, the right superior anterior temporal lobe (sATL) - hypothesized to be selectively involved in social semantics - and the right intraparietal sulcus (pIPS) - associated with numerical information and quantification. Additionally, as control site the authors used the vertex. Double-pulse TMS at 25Hz was delivered between prime and target words, 10ms after the blank following the prime word. Intensity was set at 100% of the motor threshold. The authors found that stimulation for the regions of interest, as opposed to vertex, reduced the priming effect for social words in both pIPS and sATL, and for quantity words only in the pIPS condition.

Note that we always used the original exclusion criteria for trials and subjects reported in the individual papers. Following common procedures (Van Zandt, 2002), we transformed RTs to their logarithm before entering the analyses to ensure normality of their distributions.

### Measuring the correlation between models and response times

The goal of our study was to probe different models (semantic, frequency- and orthography- based) to find out whether the TMS effects found across five datasets, and attributed originally to semantics alone, on the basis of simple contrasts of conditions (i.e. direct comparisons of RTs across stimulation conditions), were actually due to a selective semantic effect.

The intuition behind our methodology is quite straightforward: for each trial we modelled the cognitive effort in terms of the three models (semantic similarity among the words involved, sum of word frequencies or word lengths). Then we used Spearman correlation (*ρ*) to measure the association between trial-level RTs and each of the three models. We ran the analyses separately for each subject, then averaging the *ρ* scores across subjects, thus accounting for subject-specific differences in reaction times. For each model, we then compared the correlations across effective stimulation and control conditions (sham/vertex). A significant difference among the two for a given model would indicate that the TMS effect acted on that level of cognitive processing (semantic, lexical/recognition, orthographic). As detailed above, we hypothesized that significant differences among TMS conditions should emerge only for the semantic model.

## Models

### Language model semantic similarity

As a model of lexical semantics, we used so-called language models, also known as distributional semantics models. These are computational models that try to capture the meanings of words by abstracting them over large amounts of texts, called corpora (Lenci, 2018). These models can be used to answer questions that can be either theoretical - e.g. how humans acquire conceptual knowledge in the wild? (Hosseini et al., 2024; Landauer & Dumais, 1997) - or practical - e.g. can we simulate the way humans use words in natural language for applied tasks like translation? (Bengio et al., 2003; Min et al., 2023).

Although each model can differ noticeably in its implementation, they all start from a basic principle: they create vector representations for word meanings by learning from patterns of word co-occurrences in corpora. For instance, they learn that not only ‘wine’ is similar to ‘glass’, because they are often found in the same sentences; but also that ‘wine’ is similar to ‘beer’, because they are both frequently found in the same sentence as ‘glass’.

From the point of view of semantics, it has been consistently shown that such vector representations - often called word vectors - capture quite well the way in which humans represent the corresponding concepts (Schrimpf et al., 2021; Wingfield & Connell, 2022).

Since an extremely large number of models exists, we selected the language model that best captured behavioural responses - as measured from an independent task and dataset, ensuring an unbiased evaluation.

First, we selected a number of candidate models that respected three criteria: having been previously used in the cognitive literature; differentially covering the three different types of architectures and approaches to language modelling across the years (as defined in Connell and Lynott (2024) and Lenci et al. (2022): count-based models, word embeddings, contextualized - both small and large - language models); being trained exclusively in the two languages present in our work (German and Italian). We thus restricted the set of candidates to Pointwise Positive Mutual Information - PPMI (Levy et al., 2015); fasttext (Bojanowski et al., 2017); GPT2-small (Radford et al., 2019); and Llama (Dubey et al., 2024).

We used the word vectors extracted from each language model to measure semantic similarity, by which we refer to the cosine similarity between two word vector semantic representations - this is the standard way of measuring semantic similarity in the field (Turney & Pantel, 2010). Conversely, when referring to ‘semantic dissimilarity’ we refer to 1-semantic similarity, which we assume to proportionally reflect processing effort (the more dissimilar two representations, the more demanding the processing and the longer the reaction times). We also considered so-called word surprisal, a measure of conditional word probability that has been recently shown to capture to a surprising extent semantic processing (Wilcox et al., 2023). We could compute surprisal measure only for PPMI, GPT2-small, and Llama since fasttext does not allow to do so.

Secondly, we singled out the language model able to best model behavioural response times when processing semantics. We used a behavioural task, lexical decision, which seemed an ideal choice for several reasons. First, it was not used in any of the included TMS datasets, ensuring an unbiased evaluation. Secondly, lexical decision is commonly thought to cover multiple processing levels that are relevant here - sub-lexical, lexical and semantic processing (Balota et al., 2007). Finally, previous work already established that language models are able to capture the cognitive effort involved in this task (Hollis & Westbury, 2016).

We thus used publicly available reaction times data during a lexical decision task. For German, we used the dataset presented in Schröter and Schroeder (2017), counting 1152 words, and we employed only the response times from young adults. For Italian, we used the dataset published with (Vergallito et al., 2020), comprising 1121 words rated by young adults.

Finally, we computed the extent to which each model captured lexical decision times in both languages using a correlation metric. The aim was to single out the model showing the highest correlation with the lexical decision response times for further analyses. Details of the vector extraction procedure, as well as the methods and results of the evaluation are reported in the Supplementary Materials (section Details and Evaluation of the Language Models; Figures and B1). These also include the full set of results for the second-best model, GPT2-small surprisal (Figures C14). The model that performed better turned out to be **PPMI**; therefore we used this model for all the analyses reported in the main paper.

### Word frequency

To test whether TMS selectively affected semantic processing, we also repeated the analyses using word frequency, a variable that is not semantic in nature. By contrast, it is related to a lower-level, statistical learning effect: words that have been encountered more often are easier to recognize (i.e. they have lower response times in a number of tasks - see Brysbaert et al. (2018) and Kuperman et al. (2024)).

For word frequency we used, as a measure of cognitive effort, the negative of the sum of the logarithm to the base 10 of the frequencies of all words involved in each trial (following Brysbaert et al. (2018)). The intuition was that the less frequent the words, the bigger the cognitive effort should be; because of this inverse relation, we used the negative of the sum. For the production tasks, we summed the log-transformed frequency of the visually presented and the uttered words. For the semantic judgments tasks, we summed the log-transformed frequency of the visually presented words - that in all cases, except sound and action feature judgments, were two. For the latter case we just took the log-transformed frequency of the word that appeared during each trial.

Summing the frequency values could lead, in principle, to missing important parts of information regarding the TMS effect - because it may have acted only on the first or the second word. To show that this was not the case, and that summing the frequency values reliably captured the overall effort of each trial, we repeated the analyses using either only the first word or the second word’s log-transformed frequencies. Results are reported in the Supplementary Materials (Figures C14). They confirm that, overall, the sum approach is the best frequency-based model, yielding in the vast majority of cases the highest correlation.

### Word length

Additionally, we ran the analyses for word length, a purely orthographic variable that has consistently been shown to affect word processing (see Barton et al. (2014)).

Similarly to word frequency, we assumed that the longer the words, the larger the cognitive effort should be. Again, as a measure of effort, we took the sum of all word lengths involved in each trial. For production tasks, the sum of visually presented and uttered words; for semantic judgment tasks, for sound- and action- feature judgments, the length of the word to be judged in each trial; for all other cases, the sum of the two visually presented words.

Likewise, we also re-ran the analyses using either the first word’s or the second word’s length - they are reported in the Supplementary Materials (Figures Again, we found that summing the lengths of all words in a trial provided the model with the best performance.

### Hierarchical bootstrapping with subsampling

The remaining challenge was to standardize correlation scores across studies, to allow easier interpretation of the results - which is not trivial, since evaluations are affected by experimental parameters like the number of subjects and trials and cannot be directly compared (Guerra et al., 2020; Hebart & Baker, 2018). To overcome this, we used hierarchical bootstrapping with subsampling, a simple but powerful non-parametric approach. This approach has been demonstrated to provide the same rate of false positives as a linear mixed model approach, and it accounts for each the hierarchical structure of the dataset (i.e. individual subjects are resampled and evaluated separately -Saravanan et al. (2020)). It relies on simulating N repetitions of each experiment using a fixed number of subjects and trials across all studies. In our case, *N_iterations_* = 1000; for each iteration, subjects and trials were randomly subsampled with fixed *N_subjects_* = 20, *N_trials_* = 25 from the full experiments. In practice, for each iteration in a dataset - and separately for each condition and model, we sampled 20 subjects, 25 trials for each of them, computed the Spearman correlation within each subjects, then averaged the individual scores. This allowed to exactly match the number of subjects and trials across the five datasets (note that, as in the original papers, we only used correct trials; hence the lower number of trials available). Because of this, the resulting average correlations between each model and the RTs stem from standardized experimental conditions and thus have the same scale (see Figure 2 for a simplified visualization of the methodology used for the analyses).

### Statistical significance testing, correction for multiple comparisons and effect sizes

We ran two types of statistical significance tests. First, we checked whether, for each model and each TMS/control condition, the distribution of 1000 scores obtained with hierarchical bootstrapping was significantly above or below chance (thus using a two-tailed approach). To do so, we used the procedure recommended in Saravanan et al. (2020), which is similar to the one typically used in permutation testing (Nichols & Holmes, 2002). We counted the number of iterations where the average correlation was below the value corresponding to the null hypothesis (*ρ* = 0), and divided this number by *N_iterations_* = 1000. This provides a one-sided p-value, a so-called *p_bootstrap_* (Saravanan et al., 2020) - the probability of finding a result below chance when repeating the experiment. In fact, we smoothed the p-value by adding 1 to both sides of the division - this avoids having 0 as a resulting p-value, as recommended in Phipson and Smyth (2010). To obtain a two-sided p-value, we followed the implementation of the python package scipy for two-tailed permutation testing (Virtanen et al., 2020): we repeated the same operation with the inverse direction (considering average iteration values above chance); then we took the minimum of the two, and multiplied it by two, thus arriving at the p-value. Since the t-statistic is affected by the number of observations, that here is ‘artificially’ set at *N_iterations_* = 1000, we do not report t-values but only Cohen’s *d*, which is independent of sample size.

Secondly, we compared control with effective stimulation(s), separately for each dataset and each model. We ran non-parametric, two-sample two-sided permutation tests. Here we report Cohen’s *d* effect sizes for paired data.

We corrected for multiple comparisons using the False Discovery Rate (FDR; Benjamini and Hochberg (1995)). We used an extremely strict approach, correcting all p-values referring to the comparisons across all TMS datasets together once (i.e. calling only one time scipy’s *false*_*discovery*_*control* function on all p-values – including comparisons against chance and comparisons between conditions). This rigorous correction for multiple comparisons ensures keeping the risk of incurring in false positives at a minimum.

We also report 95% confidence intervals, following recent recommendations (Cumming, 2014; Hentschke & Stüttgen, 2011). As a lower bound to consider a result statistically significant, we set the threshold at *p<* 0.05, however, to avoid false positives, we complement it with thresholds for effect sizes tailored to the cognitive neuroscience literature (Szucs & Ioannidis, 2017), following recent recommendations (Correll et al., 2020; Schäfer & Schwarz, 2019). As thresholds for low, medium and strong effects we selected the 25th percentile (*d* = 0.637), median (*d* = 0.932) and 75th percentile (*d* = 1.458) of the effect sizes for statistically significant results reported in a recent review on effect sizes in cognitive neuroscience (Szucs & Ioannidis, 2017). These values are more stringent than generic effect size thresholds (*low* = 0.2, *mid* = 0.5, *high* = 0.8; Calin-Jageman (2018)) - allowing to reduce the risk of false positives and highlighting which results can be confidently interpreted as reflecting an effect of TMS. To compute effect sizes and confidence intervals, we use the python package pingouin (Vallat, 2018).

## Acknowledgements

We would like to thank the members of the Cognition and Plasticity lab for feedback on earlier versions of these results.

## References

1. Baayen, R. H., & Milin, P. (2010). Analyzing reaction times. International journal of psychological research, 3 (2), 12–28.

2. Balota, D. A., Cortese, M. J., Sergent-Marshall, S. D., Spieler, D. H., & Yap, M. J. (2004). Visual word recognition of single-syllable words. Journal of experimental psychology: General, 133 (2), 283.

3. Balota, D. A., Yap, M. J., Hutchison, K. A., Cortese, M. J., Kessler, B., Loftis, B., Neely, J. H., Nelson, D. L., Simpson, G. B., & Treiman, R. (2007). The english lexicon project. Behavior research methods, 39, 445–459.

4. Banks, B., Wingfield, C., & Connell, L. (2021). Linguistic distributional knowledge and sensorimotor grounding both contribute to semantic category production. Cognitive Science, 45 (10), e13055.

5. Baroni, M., Bernardini, S., Ferraresi, A., & Zanchetta, E. (2009). The wacky wide web: A collection of very large linguistically processed web-crawled corpora. Language resources and evaluation, 43, 209–226.

6. Barton, J. J., Hanif, H. M., Eklinder Björnström, L., & Hills, C. (2014). The word-length effect in reading: A review. Cognitive neuropsychology, 31 (5-6), 378–412.

7. Bengio, Y., Ducharme, R., Vincent, P., & Jauvin, C. (2003). A neural probabilistic language model. Journal of machine learning research, 3 (Feb), 1137–1155.

8. Benjamini, Y., & Hochberg, Y. (1995). Controlling the false discovery rate: A practical and powerful approach to multiple testing. Journal of the Royal statistical society: series B (Methodological), 57 (1), 289–300.

9. Bergmann, T. O., & Hartwigsen, G. (2021). Inferring causality from noninvasive brain stimulation in cognitive neuroscience. Journal of cognitive neuroscience, 33 (2), 195–225.

10. Bergmann, T. O., Karabanov, A., Hartwigsen, G., Thielscher, A., & Siebner, H. R. (2016). Combining non-invasive transcranial brain stimulation with neuroimaging and electrophysiology: Current approaches and future perspectives. Neuroimage, 140, 4–19.

11. Beynel, L., Appelbaum, L. G., Luber, B., Crowell, C. A., Hilbig, S. A., Lim, W., Nguyen, D., Chrapliwy, N. A., Davis, S. W., Cabeza, R., et al. (2019). Effects of online repetitive transcranial magnetic stimulation (rtms) on cognitive processing: A meta-analysis and recommendations for future studies. Neuroscience & Biobehavioral Reviews, 107, 47–58.

12. Beynel, L., Powers, J. P., & Appelbaum, L. G. (2020). Effects of repetitive transcranial magnetic stimulation on resting-state connectivity: A systematic review. Neuroimage, 211, 116596.

13. Bhatia, S., & Aka, A. (2022). Cognitive modeling with representations from large-scale digital data. Current Directions in Psychological Science, 31 (3), 207–214.

14. Binder, J. R., Desai, R. H., Graves, W. W., & Conant, L. L. (2009). Where is the semantic system? a critical review and meta-analysis of 120 functional neuroimaging studies. Cerebral cortex, 19 (12), 2767–2796.

15. Binder, J. R., Westbury, C. F., McKiernan, K. A., Possing, E. T., & Medler, D. A. (2005). Distinct brain systems for processing concrete and abstract concepts. Journal of cognitive neuroscience, 17 (6), 905–917.

16. Binkofski, F. C., Klann, J., & Caspers, S. (2016). On the neuroanatomy and functional role of the inferior parietal lobule and intraparietal sulcus. In Neurobiology of language (pp. 35–47). Elsevier.

17. Binney, R. J., & Lambon Ralph, M. A. (2015). Using a combination of fmri and anterior temporal lobe rtms to measure intrinsic and induced activation changes across the semantic cognition network. Neuropsychologia, 76, 170–181.

18. Binney, R. J., & Ramsey, R. (2020). Social semantics: The role of conceptual knowledge and cognitive control in a neurobiological model of the social brain. Neuroscience & Biobehavioral Reviews, 112, 28–38.

19. Bojanowski, P., Grave, E., Joulin, A., & Mikolov, T. (2017). Enriching word vectors with subword information. Transactions of the association for computational linguistics, 5, 135–146.

20. Boleda, G. (2020). Distributional semantics and linguistic theory. Annual Review of Linguistics, 6 (1), 213–234.

21. Bommasani, R., Davis, K., & Cardie, C. (2020). Interpreting pretrained contextualized representations via reductions to static embeddings. Proceedings of the 58th Annual Meeting of the Association for Computational Linguistics, 4758–4781.

22. Bozeat, S., Lambon Ralph, M. A., Patterson, K., & Hodges, J. R. (2002). The influence of personal familiarity and context on object use in semantic dementia. Neurocase, 8 (1-2), 127–134.

23. Bradley, C., Nydam, A. S., Dux, P. E., & Mattingley, J. B. (2022). State-dependent effects of neural stimulation on brain function and cognition. Nature Reviews Neuroscience, 23 (8), 459–475.

24. Bruera, A., & Poesio, M. (2024). Family lexicon: Using language models to encode memories of personally familiar and famous people and places in the brain. PloS One, 19 (11), e0291099.

25. Bruera, A., & Poesio, M. (2025). Electroencephalography searchlight decoding reveals person-and place-specific responses for semantic category and familiarity. Journal of Cognitive Neuroscience, 37 (1), 135–154.

26. Bruffaerts, R., De Deyne, S., Meersmans, K., Liuzzi, A. G., Storms, G., & Vandenberghe, R. (2019). Redefining the resolution of semantic knowledge in the brain: Advances made by the introduction of models of semantics in neuroimaging. Neuroscience & Biobehavioral Reviews, 103, 3–13.

27. Brysbaert, M., Mandera, P., & Keuleers, E. (2018). The word frequency effect in word processing: An updated review. Current directions in psychological science, 27 (1), 45–50.

28. Burdick, L., Kummerfeld, J. K., & Mihalcea, R. (2018). Factors influencing the surprising instability of word embeddings. Proceedings of the 2018 Conference of the North American Chapter of the Association for Computational Linguistics: Human Language Technologies, Volume 1 (Long Papers), 2092–2102.

29. Calin-Jageman, R. J. (2018). The new statistics for neuroscience majors: Thinking in effect sizes. Journal of undergraduate neuroscience education, 16 (2), E21.

30. Calzavarini, F. (2024). Rethinking modality-specificity in the cognitive neuroscience of concrete word meaning: A position paper. Language, Cognition and Neuroscience, 39 (7), 815–837.

31. Caramazza, A., & Shelton, J. R. (1998). Domain-specific knowledge systems in the brain: The animate-inanimate distinction. Journal of cognitive neuroscience, 10 (1), 1–34.

32. Catricalà, E., Conca, F., Fertonani, A., Miniussi, C., & Cappa, S. F. (2020). State-dependent tms reveals the differential contribution of atl and ips to the representation of abstract concepts related to social and quantity knowledge. Cortex, 123, 30–41.

33. Cattaneo, Z., Devlin, J. T., Salvini, F., Vecchi, T., & Silvanto, J. (2010). The causal role of category-specific neuronal representations in the left ventral premotor cortex (pmv) in semantic processing. Neuroimage, 49 (3), 2728–2734.

34. Charest, I., Kievit, R. A., Schmitz, T. W., Deca, D., & Kriegeskorte, N. (2014). Unique semantic space in the brain of each beholder predicts perceived similarity. Proceedings of the National Academy of Sciences, 111 (40), 14565–14570.

35. Chiou, R., Humphreys, G. F., Jung, J., & Lambon Ralph, M. A. (2018). Controlled semantic cognition relies upon dynamic and flexible interactions between the executive ‘semantic control’and hub-and-spoke ‘semantic representation’systems. cortex, 103, 100–116.

36. Christiansen, M. H., & Chater, N. (2008). Language as shaped by the brain. Behavioral and brain sciences, 31 (5), 489–509.

37. Connell, L., & Lynott, D. (2024). What can language models tell us about human cognition? Current Directions in Psychological Science, 33 (3), 181–189.

38. Connell, L., & Ramscar, M. (2001). Using distributional measures to model typicality in categorization. Proceedings of the Annual Meeting of the Cognitive Science Society, 23 (23).

39. Cope, T. E., Shtyrov, Y., MacGregor, L. J., Holland, R., Pulvermüller, F., Rowe, J. B., & Patterson, K. (2020). Anterior temporal lobe is necessary for efficient lateralised processing of spoken word identity. Cortex, 126, 107–118.

40. Correll, J., Mellinger, C., McClelland, G. H., & Judd, C. M. (2020). Avoid cohen’s ‘small’,‘medium’, and ‘large’for power analysis. Trends in cognitive sciences, 24 (3), 200–207.

41. Cumming, G. (2014). The new statistics: Why and how. Psychological science, 25 (1), 7–29.

42. Davey, J., Cornelissen, P. L., Thompson, H. E., Sonkusare, S., Hallam, G., Smallwood, J., & Jefferies, E. (2015). Automatic and controlled semantic retrieval: Tms reveals distinct contributions of posterior middle temporal gyrus and angular gyrus. Journal of Neuroscience, 35 (46), 15230–15239.

43. Dehaene, S., Piazza, M., Pinel, P., & Cohen, L. (2005). Three parietal circuits for number processing. In The handbook of mathematical cognition (pp. 433–453). Psychology Press.

44. de Varda, A. G., Marelli, M., & Amenta, S. (2024). Cloze probability, predictability ratings, and computational estimates for 205 english sentences, aligned with existing eeg and reading time data. Behavior Research Methods, 56 (5), 5190–5213.

45. Devlin, J. T., Matthews, P. M., & Rushworth, M. F. (2003). Semantic processing in the left inferior prefrontal cortex: A combined functional magnetic resonance imaging and transcranial magnetic stimulation study. Journal of cognitive neuroscience, 15 (1), 71–84.

46. de Zubicaray, G. I. (2023). The neural organization of language production: Evidence from neuroimaging and neuromodulation. Language Production, 97–142.

47. Drijvers, L., Small, S. L., & Skipper, J. I. (2025). Language is widely distributed throughout the brain. Nature Reviews Neuroscience, 1–1.

48. Dubey, A., Jauhri, A., Pandey, A., Kadian, A., Al-Dahle, A., Letman, A., Mathur, A., Schelten, A., Yang, A., Fan, A., et al. (2024). The llama 3 herd of models. *arXiv preprint arXiv:2407.21783*.

49. Faaß, G., & Eckart, K. (2013). Sdewac–a corpus of parsable sentences from the web. Language Processing and Knowledge in the Web: 25th International Conference, GSCL 2013, Darmstadt, Germany, September 25-27, 2013. Proceedings, 61–68.

50. Fernandino, L., Tong, J.-Q., Conant, L. L., Humphries, C. J., & Binder, J. R. (2022). Decoding the information structure underlying the neural representation of concepts. Proceedings of the National Academy of Sciences, 119 (6), e2108091119.

51. Fiebach, C. J., Friederici, A. D., Müller, K., & Von Cramon, D. Y. (2002). Fmri evidence for dual routes to the mental lexicon in visual word recognition. Journal of cognitive neuroscience, 14 (1), 11–23.

52. Finocchiaro, C., Capasso, R., Cattaneo, L., Zuanazzi, A., & Miceli, G. (2015). Thematic role assignment in the posterior parietal cortex: A tms study. Neuropsychologia, 77, 223–232.

53. Gao, Z., Zheng, L., Gouws, A., Krieger-Redwood, K., Wang, X., Varga, D., Smallwood, J., & Jefferies, E. (2023). Context free and context-dependent conceptual representation in the brain. Cerebral Cortex, 33 (1), 152–166.

54. Gatti, D., Rinaldi, L., Marelli, M., & Vecchi, T. (2022). Cerebellar involvement in distributional semantic learning: Insights from a combined tms-computational approach. *Brain Stimulation: Basic*, Translational, and Clinical Research in Neuromodulation, 15 (4), 999–1001.

55. Gatti, D., Van Vugt, F., & Vecchi, T. (2020). A causal role for the cerebellum in semantic integration: A transcranial magnetic stimulation study. Scientific Reports, 10 (1), 18139.

56. Goldstein, A., Grinstein-Dabush, A., Schain, M., Wang, H., Hong, Z., Aubrey, B., Nastase, S. A., Zada, Z., Ham, E., Feder, A., et al. (2024). Alignment of brain embeddings and artificial contextual embeddings in natural language points to common geometric patterns. Nature communications, 15 (1), 2768.

57. Guerra, A., López-Alonso, V., Cheeran, B., & Suppa, A. (2020). Variability in non-invasive brain stimulation studies: Reasons and results. Neuroscience letters, 719, 133330.

58. Hallett, M. (2007). Transcranial magnetic stimulation: A primer. Neuron, 55 (2), 187–199.

59. Han, J., & Moraga, C. (1995). The influence of the sigmoid function parameters on the speed of backpropagation learning. International workshop on artificial neural networks, 195–201.

60. Hartwigsen, G. (2015). The neurophysiology of language: Insights from non-invasive brain stimulation in the healthy human brain. Brain and language, 148, 81–94.

61. Hartwigsen, G., & Silvanto, J. (2023). Noninvasive brain stimulation: Multiple effects on cognition. The Neuroscientist, 29 (5), 639–653.

62. Hartwigsen, G., Weigel, A., Schuschan, P., Siebner, H. R., Weise, D., Classen, J., & Saur, D. (2016). Dissociating parieto-frontal networks for phonological and semantic word decisions: A condition-and-perturb tms study. Cerebral cortex, 26 (6), 2590–2601.

63. Hauk, O., Davis, M. H., & Pulvermüller, F. (2008). Modulation of brain activity by multiple lexical and word form variables in visual word recognition: A parametric fmri study. Neuroimage, 42 (3), 1185–1195.

64. Hebart, M. N., & Baker, C. I. (2018). Deconstructing multivariate decoding for the study of brain function. Neuroimage, 180, 4–18.

65. Hentschke, H., & Stüttgen, M. C. (2011). Computation of measures of effect size for neuroscience data sets. European Journal of Neuroscience, 34 (12), 1887–1894.

66. Hickok, G., & Poeppel, D. (2007). The cortical organization of speech processing. Nature reviews neuroscience, 8 (5), 393–402.

67. Hoffman, P. (2018). An individual differences approach to semantic cognition: Divergent effects of age on representation, retrieval and selection. Scientific reports, 8 (1), 8145.

68. Hollis, G., & Westbury, C. (2016). The principals of meaning: Extracting semantic dimensions from co-occurrence models of semantics. Psychonomic bulletin & review, 23, 1744–1756.

69. Hosseini, E. A., Schrimpf, M., Zhang, Y., Bowman, S., Zaslavsky, N., & Fedorenko, E. (2024). Artificial neural network language models predict human brain responses to language even after a developmentally realistic amount of training. Neurobiology of Language, 5 (1), 43–63.

70. Hussain, S. J., & Freedberg, M. V. (2024). Debunking the myth of excitatory and inhibitory repetitive transcranial magnetic stimulation in cognitive neuroscience research. Journal of Cognitive Neuroscience, 1–14.

71. Huth, A. G., De Heer, W. A., Griffiths, T. L., Theunissen, F. E., & Gallant, J. L. (2016). Natural speech reveals the semantic maps that tile human cerebral cortex. Nature, 532 (7600), 453–458.

72. Huth, A. G., Nishimoto, S., Vu, A. T., & Gallant, J. L. (2012). A continuous semantic space describes the representation of thousands of object and action categories across the human brain. Neuron, 76 (6), 1210–1224.

73. Indefrey, P. (2011). The spatial and temporal signatures of word production components: A critical update. Frontiers in psychology, 2, 255.

74. Jackson, R. L. (2021). The neural correlates of semantic control revisited. NeuroImage, 224, 117444.

75. Jefferies, E. (2013). The neural basis of semantic cognition: Converging evidence from neuropsychology, neuroimaging and tms. Cortex, 49 (3), 611–625.

76. Johns, B. T. (2021). Distributional social semantics: Inferring word meanings from communication patterns. Cognitive Psychology, 131, 101441.

77. Jones, M. N., Kintsch, W., & Mewhort, D. J. (2006). High-dimensional semantic space accounts of priming. Journal of memory and language, 55 (4), 534–552.

78. Kiela, D., & Clark, S. (2014). A systematic study of semantic vector space model parameters. Proceedings of the 2nd workshop on continuous vector space models and their compositionality (CVSC), 21–30.

79. Klaus, J., & Hartwigsen, G. (2019). Dissociating semantic and phonological contributions of the left inferior frontal gyrus to language production. Human brain mapping, 40 (11), 3279–3287.

80. Kriegeskorte, N., & Bandettini, P. (2007). Analyzing for information, not activation, to exploit high-resolution fmri. Neuroimage, 38 (4), 649–662.

81. Kriegeskorte, N., & Douglas, P. K. (2018). Cognitive computational neuroscience. Nature neuroscience, 21 (9), 1148–1160.

82. Kuhnke, P., Beaupain, M. C., Cheung, V. K., Weise, K., Kiefer, M., & Hartwigsen, G. (2020). Left posterior inferior parietal cortex causally supports the retrieval of action knowledge. Neuroimage, 219, 117041.

83. Kuperman, V., Schroeder, S., & Gnetov, D. (2024). Word length and frequency effects on text reading are highly similar in 12 alphabetic languages. Journal of Memory and Language, 135, 104497.

84. Lambon Ralph, M. A., Jefferies, E., Patterson, K., & Rogers, T. T. (2017). The neural and computational bases of semantic cognition. Nature reviews neuroscience, 18 (1), 42–55.

85. Landauer, T. K., & Dumais, S. T. (1997). A solution to plato’s problem: The latent semantic analysis theory of acquisition, induction, and representation of knowledge. Psychological review, 104 (2), 211.

86. Lenci, A. (2018). Distributional models of word meaning. Annual review of Linguistics, 4 (1), 151–171.

87. Lenci, A., Sahlgren, M., Jeuniaux, P., Cuba Gyllensten, A., & Miliani, M. (2022). A comparative evaluation and analysis of three generations of distributional semantic models. Language resources and evaluation, 56 (4), 1269–1313.

88. Levelt, W. J. (1999). Models of word production. Trends in cognitive sciences, 3 (6), 223–232.

89. Levy, O., Goldberg, Y., & Dagan, I. (2015). Improving distributional similarity with lessons learned from word embeddings. Transactions of the association for computational linguistics, 3, 211–225.

90. Lopopolo, A., Fedorenko, E., Levy, R., & Rabovsky, M. (2024). Cognitive computational neuroscience of language: Using computational models to investigate language processing in the brain.

91. Mandera, P., Keuleers, E., & Brysbaert, M. (2017). Explaining human performance in psycholinguistic tasks with models of semantic similarity based on prediction and counting: A review and empirical validation. Journal of Memory and Language, 92, 57–78.

92. Martin, A. (2007). The representation of object concepts in the brain. Annu. Rev. Psychol., 58 (1), 25–45.

93. Mazzuca, C., Majid, A., Lugli, L., Nicoletti, R., & Borghi, A. M. (2020). Gender is a multifaceted concept: Evidence that specific life experiences differentially shape the concept of gender. Language and Cognition, 12 (4), 649–678.

94. Min, B., Ross, H., Sulem, E., Veyseh, A. P. B., Nguyen, T. H., Sainz, O., Agirre, E., Heintz, I., & Roth, D. (2023). Recent advances in natural language processing via large pre-trained language models: A survey. ACM Computing Surveys, 56 (2), 1–40.

95. Minarik, T., Berger, B., Althaus, L., Bader, V., Biebl, B., Brotzeller, F., Fusban, T., Hegemann, J., Jesteadt, L., Kalweit, L., et al. (2016). The importance of sample size for reproducibility of tdcs effects. Frontiers in human neuroscience, 10, 453.

96. Miniussi, C., Harris, J. A., & Ruzzoli, M. (2013). Modelling non-invasive brain stimulation in cognitive neuroscience. Neuroscience & Biobehavioral Reviews, 37 (8), 1702–1712.

97. Mitchell, T. M., Shinkareva, S. V., Carlson, A., Chang, K.-M., Malave, V. L., Mason, R. A., & Just, M. A. (2008). Predicting human brain activity associated with the meanings of nouns. science, 320 (5880), 1191–1195.

98. Murdoch, B. E. (2010). The cerebellum and language: Historical perspective and review. Cortex, 46 (7), 858–868.

99. Nair, S., & Resnik, P. (2023). Words, subwords, and morphemes: What really matters in the surprisal-reading time relationship? Findings of the Association for Computational Linguistics: EMNLP 2023, 11251–11260.

100. Naselaris, T., & Kay, K. N. (2015). Resolving ambiguities of mvpa using explicit models of representation. Trends in cognitive sciences, 19 (10), 551–554.

101. Naselaris, T., Kay, K. N., Nishimoto, S., & Gallant, J. L. (2011). Encoding and decoding in fmri. Neuroimage, 56 (2), 400–410.

102. Nichols, T. E., & Holmes, A. P. (2002). Nonparametric permutation tests for functional neuroimaging: A primer with examples. Human brain mapping, 15 (1), 1–25.

103. Numssen, O., van der Burght, C. L., & Hartwigsen, G. (2023). Revisiting the focality of non-invasive brain stimulation–implications for studies of human cognition. Neuroscience & Biobehavioral Reviews, 149, 105154.

104. Olson, I. R., McCoy, D., Klobusicky, E., & Ross, L. A. (2013). Social cognition and the anterior temporal lobes: A review and theoretical framework. Social cognitive and aJective neuroscience, 8 (2), 123–133.

105. Pammer, K., Hansen, P. C., Kringelbach, M. L., Holliday, I., Barnes, G., Hillebrand, A., Singh, K. D., & Cornelissen, P. L. (2004). Visual word recognition: The first half second. Neuroimage, 22 (4), 1819–1825.

106. Papeo, L., Pascual-Leone, A., & Caramazza, A. (2013). Disrupting the brain to validate hypotheses on the neurobiology of language. Frontiers in Human Neuroscience, 7, 148.

107. Peelen, M. V., & Downing, P. E. (2023). Testing cognitive theories with multivariate pattern analysis of neuroimaging data. Nature human behaviour, 7 (9), 1430–1441.

108. Pell, G. S., Roth, Y., & Zangen, A. (2011). Modulation of cortical excitability induced by repetitive transcranial magnetic stimulation: Influence of timing and geometrical parameters and underlying mechanisms. Progress in neurobiology, 93 (1), 59–98.

109. Pereira, F., Lou, B., Pritchett, B., Ritter, S., Gershman, S. J., Kanwisher, N., Botvinick, M., & Fedorenko, E. (2018). Toward a universal decoder of linguistic meaning from brain activation. Nature communications, 9 (1), 963.

110. Phipson, B., & Smyth, G. K. (2010). Permutation p-values should never be zero: Calculating exact p-values when permutations are randomly drawn. Statistical applications in genetics and molecular biology, 9 (1).

111. Piai, V., Nieberlein, L., & Hartwigsen, G. (2020). Effects of transcranial magnetic stimulation over the left posterior superior temporal gyrus on picture-word interference. Plos one, 15 (11), e0242941.

112. Piantadosi, S. T., Tily, H., & Gibson, E. (2011). Word lengths are optimized for efficient communication. Proceedings of the National Academy of Sciences, 108 (9), 3526–3529.

113. Pitcher, D., Parkin, B., & Walsh, V. (2021). Transcranial magnetic stimulation and the understanding of behavior. Annual Review of Psychology, 72 (1), 97–121.

114. Pobric, G., Jefferies, E., & Lambon Ralph, M. A. (2010). Category-specific versus category-general semantic impairment induced by transcranial magnetic stimulation. Current biology, 20 (10), 964–968.

115. Pobric, G., Mashal, N., Faust, M., & Lavidor, M. (2008). The role of the right cerebral hemisphere in processing novel metaphoric expressions: A transcranial magnetic stimulation study. Journal of cognitive neuroscience, 20 (1), 170–181.

116. Poldrack, R. A., Baker, C. I., Durnez, J., Gorgolewski, K. J., Matthews, P. M., Munafò, M. R., Nichols, T. E., Poline, J.-B., Vul, E., & Yarkoni, T. (2017). Scanning the horizon: Towards transparent and reproducible neuroimaging research. Nature reviews neuroscience, 18 (2), 115–126.

117. Poldrack, R. A., Wagner, A. D., Prull, M. W., Desmond, J. E., Glover, G. H., & Gabrieli, J. D. (1999). Functional specialization for semantic and phonological processing in the left inferior prefrontal cortex. Neuroimage, 10 (1), 15–35.

118. Pulvermüller, F., Hauk, O., Nikulin, V. V., & Ilmoniemi, R. J. (2005). Functional links between motor and language systems. European Journal of Neuroscience, 21 (3), 793–797.

119. Qu, X., Wang, Z., Cheng, Y., Xue, Q., Li, Z., Li, L., Feng, L., Hartwigsen, G., & Chen, L. (2022). Neuromodulatory effects of transcranial magnetic stimulation on language performance in healthy participants: Systematic review and meta-analysis. Frontiers in Human Neuroscience, 16, 1027446.

120. Radford, A., Wu, J., Child, R., Luan, D., Amodei, D., Sutskever, I., et al. (2019). Language models are unsupervised multitask learners. OpenAI blog, 1 (8), 9.

121. Rao, A., Monteiro, J. M., Mourao-Miranda, J., Initiative, A. D., et al. (2017). Predictive modelling using neuroimaging data in the presence of confounds. NeuroImage, 150, 23–49.

122. Renoult, L., Irish, M., Moscovitch, M., & Rugg, M. D. (2019). From knowing to remembering: The semantic–episodic distinction. Trends in cognitive sciences, 23 (12), 1041–1057.

123. Rice, G. E., Caswell, H., Moore, P., Hoffman, P., & Lambon Ralph, M. A. (2018). The roles of left versus right anterior temporal lobes in semantic memory: A neuropsychological comparison of postsurgical temporal lobe epilepsy patients. Cerebral Cortex, 28 (4), 1487–1501.

124. Rogers, T. T., Patterson, K., Jefferies, E., & Lambon Ralph, M. A. (2015). Disorders of representation and control in semantic cognition: Effects of familiarity, typicality, and specificity. Neuropsychologia, 76, 220–239.

125. Romei, V., Thut, G., & Silvanto, J. (2016). Information-based approaches of noninvasive transcranial brain stimulation. Trends in Neurosciences, 39 (11), 782–795.

126. Rosch, E., & Mervis, C. B. (1975). Family resemblances: Studies in the internal structure of categories. Cognitive psychology, 7 (4), 573–605.

127. Rosch, E. H. (1973). Natural categories. Cognitive psychology, 4 (3), 328–350.

128. Ruzzoli, M., Marzi, C. A., & Miniussi, C. (2010). The neural mechanisms of the effects of transcranial magnetic stimulation on perception. Journal of Neurophysiology, 103 (6), 2982–2989.

129. Sánchez, A., Carreiras, M., & Paz-Alonso, P. M. (2023). Word frequency and reading demands modulate brain activation in the inferior frontal gyrus. Scientific Reports, 13 (1), 17217.

130. Santacroce, F., Cachia, A., Fragueiro, A., Grande, E., Roell, M., Baldassarre, A., Sestieri, C., & Committeri, G. (2024). Human intraparietal sulcal morphology relates to individual differences in language and memory performance. Communications Biology, 7 (1), 520.

131. Saravanan, V., Berman, G. J., & Sober, S. J. (2020). Application of the hierarchi cal bootstrap to multi-level data in neuroscience. Neurons, behavior, data analysis and theory, 3 (5), https–nbdt.

132. Schäfer, T., & Schwarz, M. A. (2019). The meaningfulness of effect sizes in psychological research: Differences between sub-disciplines and the impact of potential biases. Frontiers in psychology, 10, 813.

133. Schrimpf, M., Blank, I. A., Tuckute, G., Kauf, C., Hosseini, E. A., Kanwisher, N., Tenenbaum, J. B., & Fedorenko, E. (2021). The neural architecture of language: Integrative modeling converges on predictive processing. Proceedings of the National Academy of Sciences, 118 (45), e2105646118.

134. Schroën, J. A., Gunter, T. C., Numssen, O., Kroczek, L. O., Hartwigsen, G., & Friederici, A. D. (2023). Causal evidence for a coordinated temporal interplay within the language network. Proceedings of the National Academy of Sciences, 120 (47), e2306279120.

135. Schröter, P., & Schroeder, S. (2017). The developmental lexicon project: A behavioral database to investigate visual word recognition across the lifespan. Behavior Research Methods, 49, 2183–2203.

136. Schuler, A.-L., & Hartwigsen, G. (2024). The potential of interleaved tms-fmri for linking stimulation-induced changes in task-related activity with behavioral modulations. Brain Stimulation.

137. Siebner, H. R., Funke, K., Aberra, A. S., Antal, A., Bestmann, S., Chen, R., Classen, J., Davare, M., Di Lazzaro, V., Fox, P. T., et al. (2022). Transcranial magnetic stimulation of the brain: What is stimulated?–a consensus and critical position paper. Clinical Neurophysiology, 140, 59–97.

138. Silvanto, J., Muggleton, N. G., Cowey, A., & Walsh, V. (2007). Neural adaptation reveals state-dependent effects of transcranial magnetic stimulation. European Journal of Neuroscience, 25 (6), 1874–1881.

139. Smith, N. J., & Levy, R. (2013). The effect of word predictability on reading time is logarithmic. Cognition, 128 (3), 302–319.

140. Snoek, L., MiletiÊ, S., & Scholte, H. S. (2019). How to control for confounds in decoding analyses of neuroimaging data. Neuroimage, 184, 741–760.

141. Szucs, D., & Ioannidis, J. P. (2017). Empirical assessment of published effect sizes and power in the recent cognitive neuroscience and psychology literature. PLoS biology, 15 (3), e2000797.

142. Thompson, H., Davey, J., Hoffman, P., Hallam, G., Kosinski, R., Howkins, S., Wooffindin, E., Gabbitas, R., & Jefferies, E. (2017). Semantic control deficits impair understanding of thematic relationships more than object identity. Neuropsychologia, 104, 113–125.

143. Trott, S., & Bergen, B. (2023). Word meaning is both categorical and continuous. Psychological Review, 130 (5), 1239.

144. Tuckute, G., Kanwisher, N., & Fedorenko, E. (2024). Language in brains, minds, and machines. Annual Review of Neuroscience, 47 (2024), 277–301.

145. Turney, P. D., & Pantel, P. (2010). From frequency to meaning: Vector space models of semantics. Journal of artificial intelligence research, 37, 141–188.

146. Valero-Cabré, A., Amengual, J. L., Stengel, C., Pascual-Leone, A., & Coubard, O. A. (2017). Transcranial magnetic stimulation in basic and clinical neuroscience: A comprehensive review of fundamental principles and novel insights. Neuroscience & Biobehavioral Reviews, 83, 381–404.

147. Vallat, R. (2018). Pingouin: Statistics in python. J. Open Source Softw., 3 (31), 1026.

148. Van Zandt, T. (2002). Analysis of response time distributions. *Stevens’* handbook of experimental psychology, 4, 461–516.

149. van Hoef, R., Connell, L., & Lynott, D. (2023). The effects of sensorimotor and linguistic information on the basic-level advantage. Cognition, 241, 105606.

150. Vergallito, A., Petilli, M. A., & Marelli, M. (2020). Perceptual modality norms for 1,121 italian words: A comparison with concreteness and imageability scores and an analysis of their impact in word processing tasks. Behavior Research Methods, 52 (4), 1599–1616.

151. Virtanen, P., Gommers, R., Oliphant, T. E., Haberland, M., Reddy, T., Cournapeau, D., Burovski, E., Peterson, P., Weckesser, W., Bright, J., et al. (2020). Scipy 1.0: Fundamental algorithms for scientific computing in python. Nature methods, 17 (3), 261–272.

152. VuliÊ, I., Ponti, E. M., Litschko, R., Glavaö, G., & Korhonen, A. (2020). Probing pretrained language models for lexical semantics. Proceedings of the 2020 Conference on Empirical Methods in Natural Language Processing (EMNLP), 7222–7240.

153. Westera, M., Gupta, A., Boleda, G., & Padó, S. (2021). Distributional models of category concepts based on names of category members. Cognitive Science, 45 (9), e13029.

154. Wilcox, E. G., Pimentel, T., Meister, C., Cotterell, R., & Levy, R. P. (2023). Testing the predictions of surprisal theory in 11 languages. Transactions of the Association for Computational Linguistics, 11, 1451–1470.

155. Wilkinson, M. D., Dumontier, M., Aalbersberg, I. J., Appleton, G., Axton, M., Baak, A., Blomberg, N., Boiten, J.-W., da Silva Santos, L. B., Bourne, P. E., et al. (2016). The fair guiding principles for scientific data management and stewardship. Scientific data, 3 (1), 1–9.

156. Wingfield, C., & Connell, L. (2022). Understanding the role of linguistic distributional knowledge in cognition. Language, Cognition and Neuroscience, 37 (10), 1220–1270.

157. Yonelinas, A. P. (2002). The nature of recollection and familiarity: A review of 30 years of research. Journal of memory and language, 46 (3), 441–517.

158. Zhou, K., Ethayarajh, K., Card, D., & Jurafsky, D. (2022). Problems with cosine as a measure of embedding similarity for high frequency words. Proceedings of the 60^th^ Annual Meeting of the Association for Computational Linguistics (Volume 2: Short Papers), 401–423.

