## Supplementary Materials for "A novel approach to map the causal impact of brain stimulation on semantic processing with language models"

#### Author Note

Andrea Bruera https://orcid.org/0000-0003-2484-2483

Gesa Hartwigsen https://orcid.org/0000-0002-8084-1330

Correspondence concerning this article should be addressed to Andrea Bruera, Max Planck Institute for Human Cognitive and Brain Sciences, Stephanstrasse 1A, 04103 Leipzig, Germany.

### Appendix A

#### Datasets

##### *Picture naming with interference*

This dataset was originally described in Piai et al. (2020). The experiment was carried out in German. The sample size in terms of subjects was 24; each subject completed 174 trials per condition (TMS and vertex). The task was picture naming with interference. During each trial, subjects first saw a picture depicting a concrete entity, accompanied by a word that could be either matched, semantically related or unrelated to the image. Then they had to name the referent of the picture. In total, 58 pictures from 14 concrete semantic categories were used. TMS was applied on left mid-to-posterior superior temporal gyrus (pSTG), an area reported to play an important role in semantic control, and particularly in resolving semantic interference (i.e. correctly naming a picture despite the presence of a distracting word; Jackson (2021)). Stimulation was online, consisting of burst of five pulses at 10 Hz delivered at each picture onset. TMS intensity was set to 90% of the individual motor threshold of the left primary motor hand area. The authors reported a significant facilitatory effect on RTs (i.e. lower RTs for TMS vs vertex) for a subset of the overall trials (cases where task difficulty was higher, and both the picture and the word referred to the same concept).

For language models, we modelled the cognitive processing by measuring the semantic dissimilarity between the word appearing below the image and the word uttered by the subject. We assumed that these measures should index a participant's effort at carrying out the task (the larger the semantic distance between the word to be uttered and the one appearing on screen, the harder the interference, hence the task requirements).

##### *Semantic production*

The second dataset was described in Klaus and Hartwigsen (2019). The experiment was carried out in German as well. The sample size of subjects was 24; each subject completed 159 trials per condition (two TMS conditions and vertex). The experiment consisted of two production tasks - one phonological and one semantic; given our focus, we only analyzed the RTs for the semantic one. Each trial consisted of a concrete word presented on screen, and the subject's task was to produce a member of the same semantic category (e.g. input: 'cat', acceptable output: 'dog'). TMS was applied at two sites: the posterior and anterior portions of the inferior frontal gyrus (pIFG and aIFG respectively). Such regions were chosen because of their putative selective engagement for, respectively, phonological and semantic processing (Poldrack et al., 1999). TMS intensity was set to 90% of the motor threshold for the right hand region of the primary motor cortex. The authors reported an inhibitory effect on semantic processing when stimulating the aIFG (i.e. higher RTs for TMS vs vertex).

As a measure of cognitive effort for language models, we used the semantic dissimilarity between visually presented and uttered word. We assume, therefore, that, given a word as stimulus (e.g. 'cat'), it is easier to produce a member of the same category that is more similar to the stimulus (e.g. 'dog') rather than one which is less similar (e.g. 'parrot').

***Semantic relatedness judgment***

The third dataset was first analyzed in Gatti et al. (2022). The experiment took place in Italian. The sample size in terms of subjects was 40, with each one of them completing 70 trials per condition (TMS and vertex). During each trial participant saw two words (one noun and one adjective; e.g. 'blue', 'apricot') presented in rapid serial visual presentation, and then had to judge whether the two were semantically related. Words were semantically related in half of the trials. Online 20Hz triple-pulse TMS was applied either at the noun or adjective onset, in a counterbalanced fashion. The stimulated area was the right cerebellum, that has started to be recently associated with semantic processing (Murdoch, 2010). Results revealed that the TMS effect shifted from facilitation to inhibition in semantic processing as the similarity between nouns and adjectives increased.

Here modelling of cognitive processing was straightforward: for language models, we took the semantic dissimilarity between the adjective and the noun as the index of cognitive effort (i.e. the closer two words in meaning, the easier it is to judge them as being similar).

##### ***Sound and action feature judgment***

This dataset was first presented in Kuhnke et al., 2020. The language of the experiment was once again German. The number of subjects was 26, that completed 52 trials per task and condition (sound and action feature judgment; TMS and sham). As anticipated, there were two different, but structurally identical tasks: auditory feature and action feature judgment (i.e. 'is the following concept related to sound?', 'is the following concept related to action?'). The experiment was divided in blocks, whose task was previously declared (i.e. sound or action). In each trial, a word referring to a concrete object was presented and participants had to indicate whether it was related to the block's feature. TMS was delivered as four online 10Hz pulses 100ms after word onset. The brain area of interest was the posterior inferior parietal lobe (pIPL), found to be consistently activated by a large number of semantic processing tasks (Jackson, 2021). The authors found a significant effect on accuracy (importantly, not on RTs) for the action feature judgment task alone.

Here, since each trial involved only one word (i.e. the word to be judged as either sound-related or action-related), we used a different approach, inspired by the concept of semantic prototype (E. H. Rosch, 1973). We started from the original intuition of E. Rosch and Mervis (1975): the difficulty of categorizing a concept, as measured in reaction times, depends on the similarity between the concept and the prototype for the category to be judged. For instance, for the category 'furniture', 'chair' is easier to judge as belonging to 'furniture' than 'radio'. This is because the underlying process would correspond to trying to match the concept and the prototype, and the closer the match -i.e. the similarity- between the two, the easier it is to provide the answer. We extend by analogy this reasoning to our case. We assume that the effort (i.e. response times) when judging whether a concept was sound- or action- related can be modelled in terms of semantic similarity to a sound- and action- prototype (i.e. the centroid, or average representation, of the concepts referring to something that makes sound and can be interacted with the hands).

We created such prototype by averaging the word vectors for the words in the set of stimuli that were both sound- and action- related. This has been shown to be the best way to produce a prototype with language models (Westera et al., 2021). Then, for each trial, we modelled the underlying cognitive processing by measuring the semantic dissimilarity between the trial's word and the sound-and-action prototype (something similar, although without the *ad-hoc* creation of a prototype vector, had been done in Connell and Ramscar (2001)).

##### *Semantic priming*

The final dataset was introduced in Catricalà et al. (2020). It was run in Italian, testing two types of semantic priming - using primes referring to either social (i.e. 'sociability') or quantity (i.e. 'immensity') concepts. Each trial involved a judgment task, whose response time was measured and analyzed. Subjects were asked whether the target word was either social or quantity-related; successful semantic priming should lead to reduced reaction times when the target word was preceded by its category as a prime. Following the original paper, we ran the analyses on the subset of subjects for which the original results were reported - i.e. 22 subjects for social and 20 for quantity concept priming. There were two stimulation sites of interest, the right superior anterior temporal lobe (sATL) - hypothesized to be selectively involved in social semantics - and the right intraparietal sulcus (pIPS) - associated with numerical information and quantification. Additionally, as control site the authors used the vertex. For each condition, there were 56 trials for each priming type (social and quantity), involving 28 congruent (e.g. social prime, social target) and 28 incongruent (e.g. social prime, quantity target) trials. Double-pulse TMS at 25Hz was delivered between prime and target words, 10ms after the blank following the prime word. Intensity was set at 100% of the motor threshold. The authors found that stimulation for the regions of interest, as opposed to vertex, reduced the priming effect for social words in both pIPS and sATL, and for quantity words only in the pIPS condition.

To model the cognitive processing carried out by participants, we used the well-established approach introduced in Jones et al. (2006) - consisting of measuring the semantic dissimilarity between prime and target words. The assumption was that a lower dissimilarity (higher similarity) between prime and target should make the task easier, hence reducing reaction times.

### Appendix B

#### Details and evaluation of the language models

We report here the fundamental details of the procedure used to create and extract vectors and surprisal scores from the PPMI language model, since it is a model created from scratch. For the pretrained models (fasttext, GPT2-small, Llama) we only report the vector extraction and surprisal computation procedures; for training details see the original papers.

**PPMI model creation.** We follow standard practice in the field (Kiela & Clark, 2014; Lenci et al., 2022; Levy et al., 2015). We use the Wacky wide web corpora (Baroni et al., 2009), that are available in both Italian and German and are the most commonly used datasets in the cognitive literature (Gatti et al., 2020; Lenci et al., 2022; Mandera et al., 2017). For German, we used sDeWac, a cleaned version of the original DeWac corpus, counting around 900M tokens (Faaß & Eckart, 2013). For Italian, we used the basic ItWac corpus, counting around 1.5B tokens.

We compute co-occurrences among content words (nouns, verbs, adjectives, adverbs) in each of the corpora within a symmetrical window of 10 words around each word. Then, we transform the counts according to the PPMI procedure (Levy et al., 2015). This provides us with a square matrix, where each line corresponds to the PPMI-reweighted co-occurrences for word $x$ with all other words - this constitutes also, in practical terms, the PPMI word vector. Since the dimensionality of the word vectors plays a key role in the amount of semantic information contained in them (Kiela & Clark, 2014), we sort the column words by their absolute frequency, and extract word vectors in increasingly bigger dimensionality, from 100 dimensions to 500000, following the recommendations Kiela and Clark (2014) (i.e. the initial 100 dimensions correspond to the 100 most common words in the corpus, and the final 500000 dimensions correspond to the 500000 most common words).

We follow different strategies to compute **word similarity** depending on the family (static vs contextualized):

- **Static LMs (fasttext and PPMI)**: we extract the previously created word vectors for each word; then compute cosine similarities among each pair of words $x$ and $y$ appearing in a trial. For PPMI we repeat this procedure multiple times, each time increasing the dimensionality (i.e. the number of words included in the columns of the PPMI-transformed co-occurence matrix) up until a limit of 500000 words, sorted them by absolute frequency in the corpus, as recommended in Kiela and Clark (2014);
- **Contextualized LMs (GPT2-small, Llama)**: we experiment with two methodologies typically used in the literature - extracting the LM's activations for the word in isolation, and so-called representation pooling (i.e. encoding 10 sentences where a word is mentioned, then averaging the activations at each layer for the subword tokens of the relevant word - Bommasani et al. (2020) and Vulić et al. (2020)). We found that the first option provided better results, so we extracted the hidden activations across all layers using the words in isolation. Then, separately for each layer, we computed the cosine similarities among each pair of words $x$ and $y$ appearing in a trial.

Similarly for **word surprisal** (note that fasttext does not allow to compute word surprisal):

- **Static LM (PPMI)**: we use, as is standard in the field, a n-gram measure of association between words - i.e. the negative logarithm of the probability of finding word $y$ in linear order after word $x$ in the Wac corpus, considering at most $n = 5$ words after $x$ - technically, we thus measure 5-gram word surprisal (de Varda et al., 2024; Smith & Levy, 2013).
- **Contextualized LM (GPT2-small, Llama)**: First, we encode with the LM word $x$ followed by word $y$. Then, we use the most common implementation of word

surprisal in the field (Nair & Resnik, 2023), corresponding to the negative logarithm of the sum of the probabilities for each subword token composing word $y$.

We report here the results for the evaluation of the language models, that we carried out on datasets involving an independent behavioural task involving semantics (lexical decision), so as to find out in an unbiased way the best model to capture how cognitive effort is reflected in RTs during semantic processing. We ran the analyses both in Italian and German; in both cases, the best performing model is PPMI, both in Italian and German - that reaches a *plateau* in performance at a dimnesionality of around 100000.

For German, we used the dataset presented in Schröter and Schroeder (2017), counting 1152 words, and we employed only the response times from young adults. For Italian, we used the dataset published with (Vergallito et al., 2020), comprising 1121 words rated by young adults.

**Figure B1**

*Model evaluation with lexical decision RTs in Italian. The x axis corresponds to vector dimensionality for PPMI, and to* $layer * 10000$ *for contextualized language models (e.g. for* $layer = 12$*,* $x = 120000$*).*

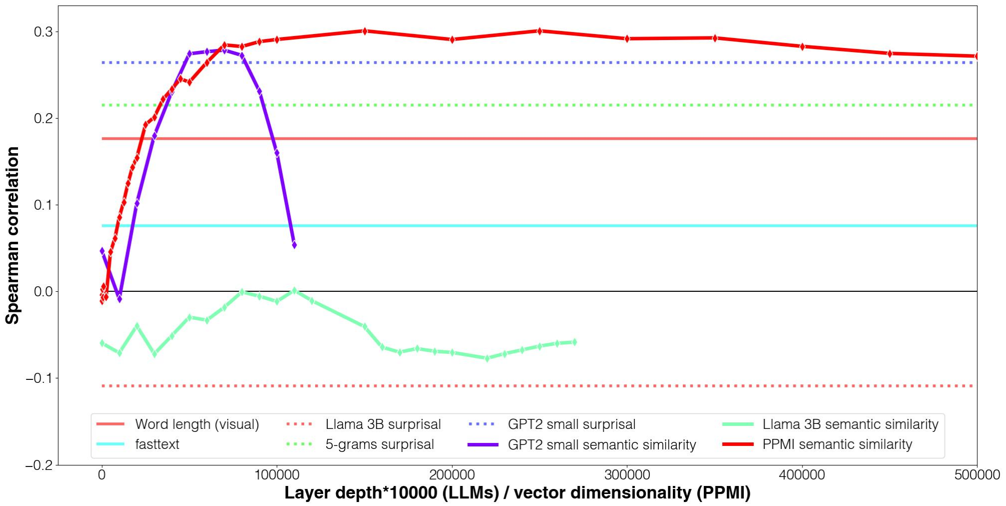

**Figure B2**

*Model evaluation with lexical decision RTs in German. The x axis corresponds to vector dimensionality for PPMI, and to* $layer * 10000$ *for contextualized language models (e.g. for* $layer = 12$, $x = 120000$*).*

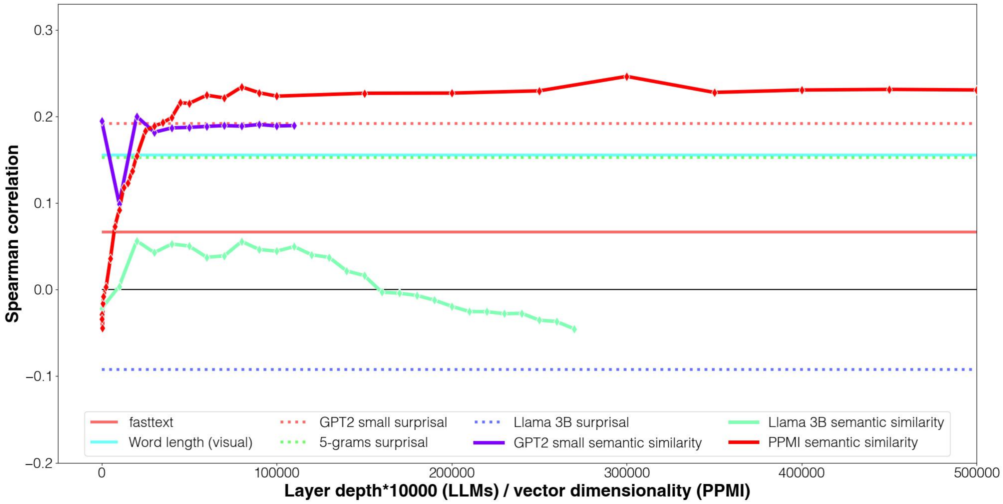

### Appendix C

#### Additional results

In the following we complement the results reported in the main text by reporting additional results for:

- the second-best model, i.e. **word surprisal values extracted from GPT2-small**;
- **word length, separately for the first and the second word** appearing in each trial (with the exception of semantic feature judgment, where there was only one word appearing);
- **word frequency, separately for the first and the second word** appearing in each trial (with the exception of semantic feature judgment, where there was only one word appearing);

Additionally, we report the full set of results **before and after removing the variance explainable by word length and word frequency** with residualization.

First we report the plots; then the table containing the full set of statistics, both when running statistical tests against the null hypothesis and when comparing across conditions.

**Figure C1**

*Additional results for* ***picture naming with interference, no residualization***

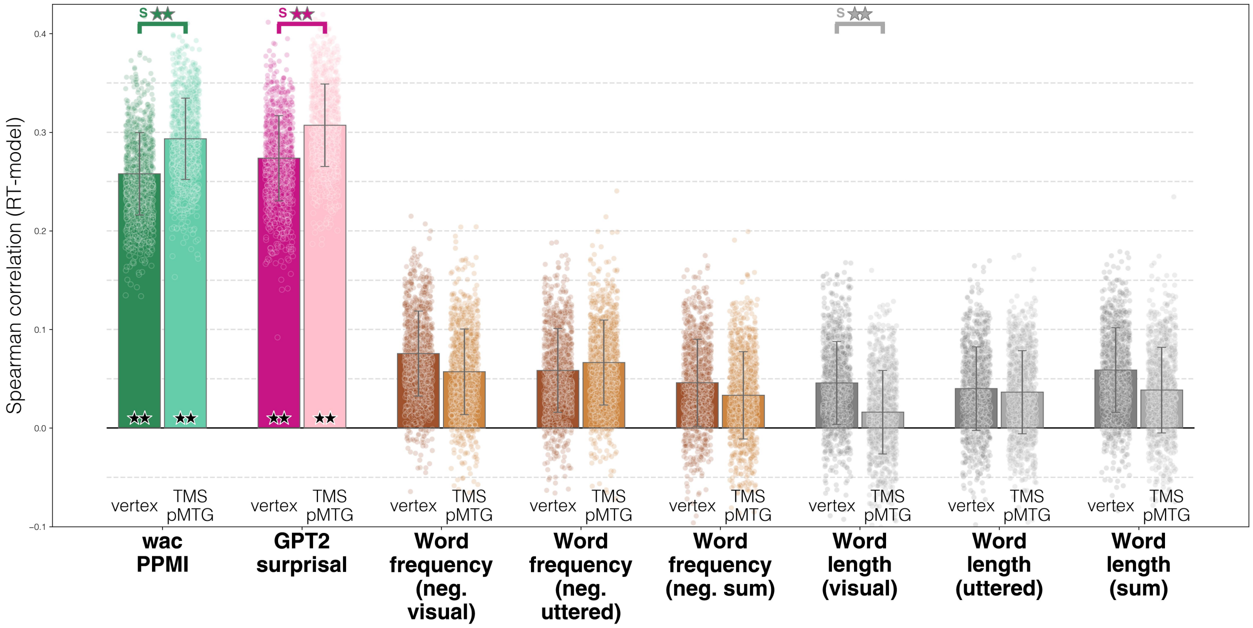


**Figure C2**

*Additional results for* ***picture naming with interference, after residualization*** *of word length and frequency*

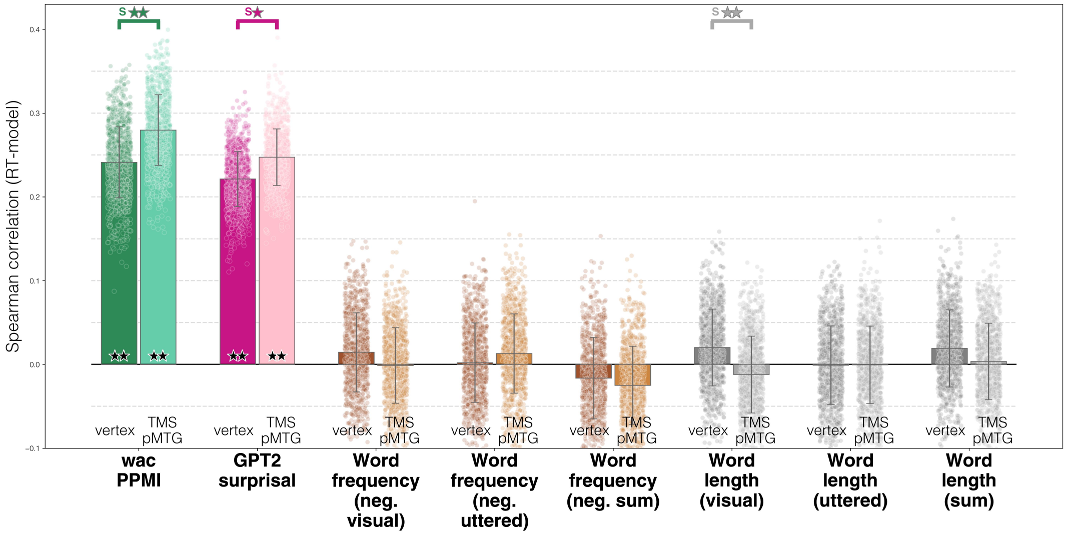

**Figure C3**

*Additional results for* ***semantic production, no residualization***

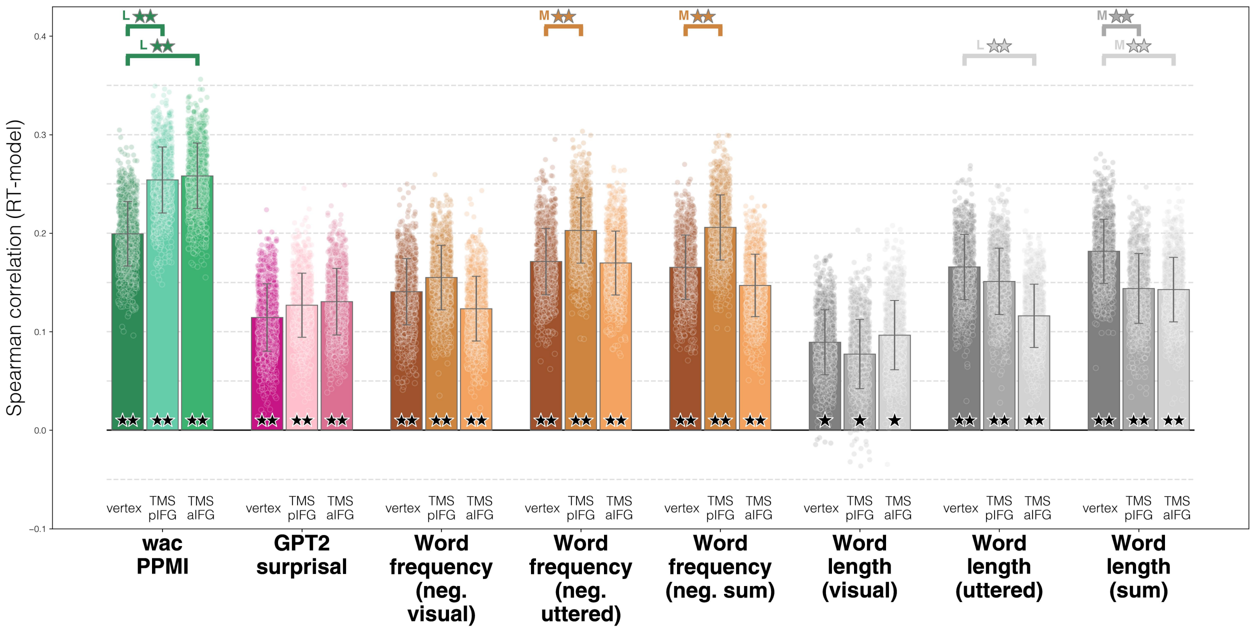


**Figure C4**

*Additional results for* ***semantic production, after residualization*** *of word length and frequency*

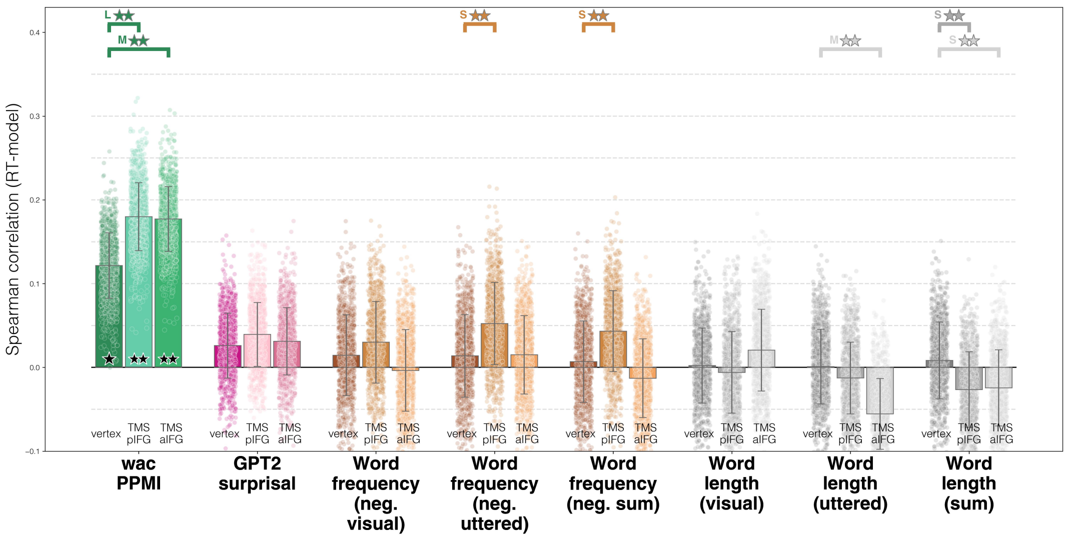

**Figure C5**

*Additional results for* ***semantic relatedness judgment, no residualization***

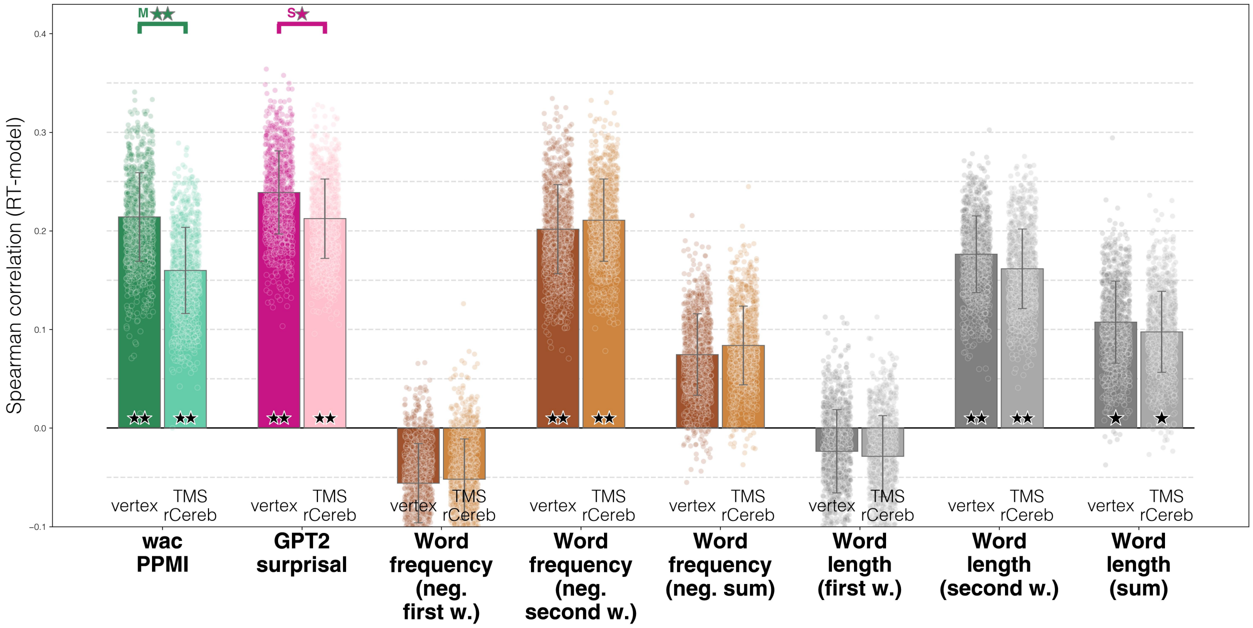


**Figure C6**

*Additional results for* ***semantic relatedness judgment, after residualization*** *of word length and frequency*

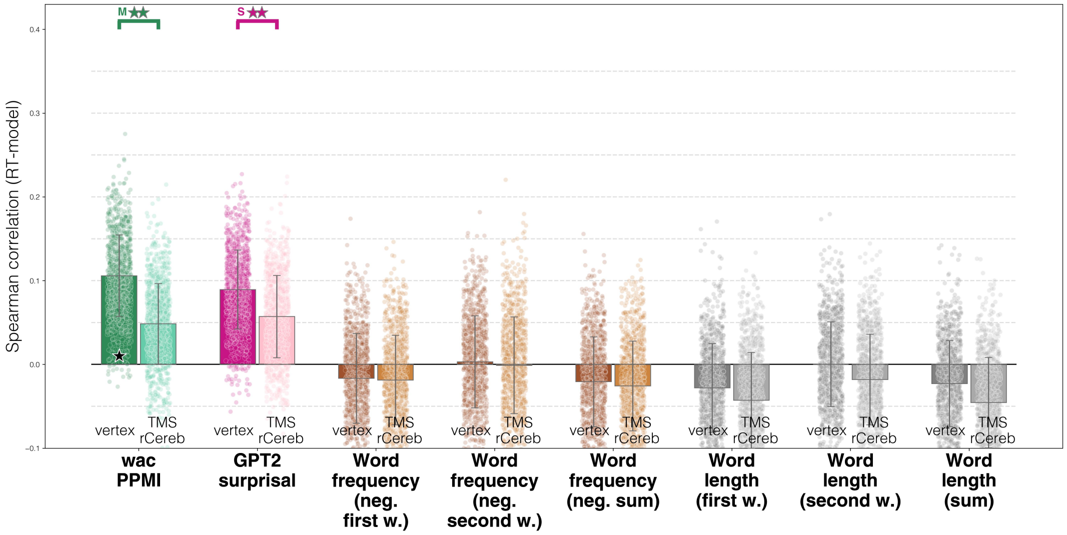

**Figure C7**

*Additional results for* ***action feature judgment, no residualization***

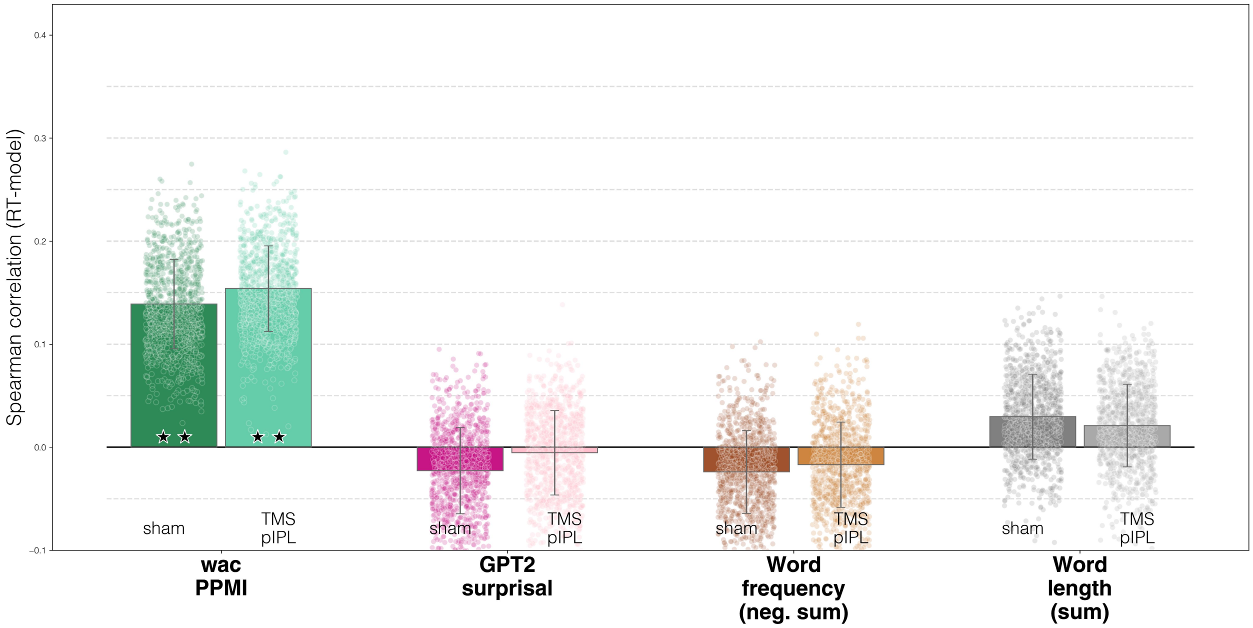


**Figure C8**

*Additional results for action feature judgment, after residualization of word length and frequency*

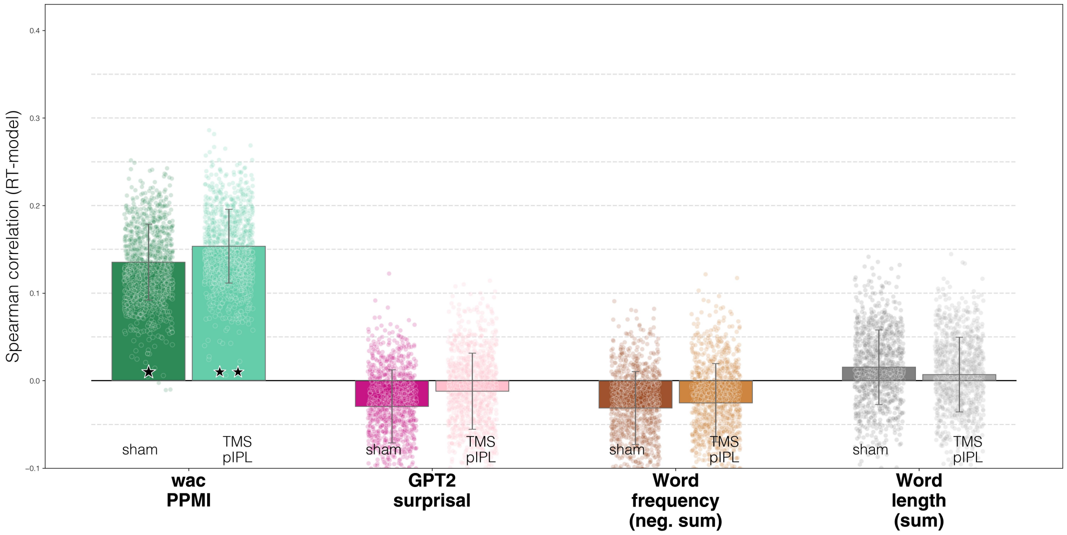

**Figure C9**

*Additional results for* ***sound feature judgment, no residualization***

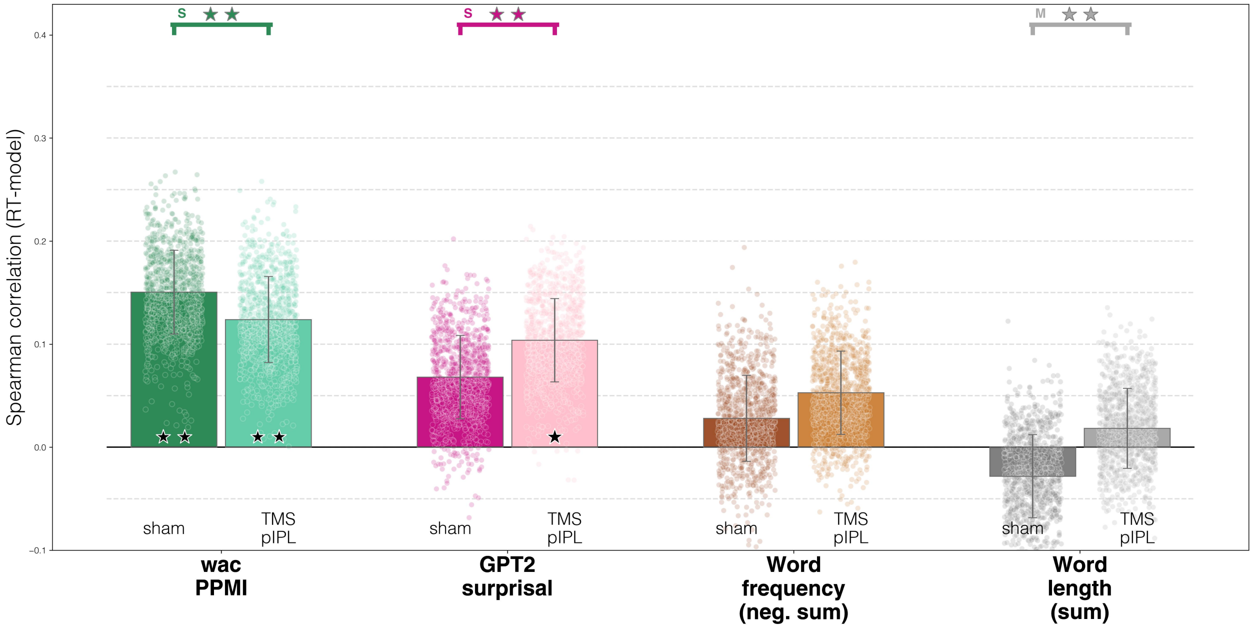


**Figure C10**

*Additional results for* ***sound feature judgment, after residualization*** *of word length and frequency*

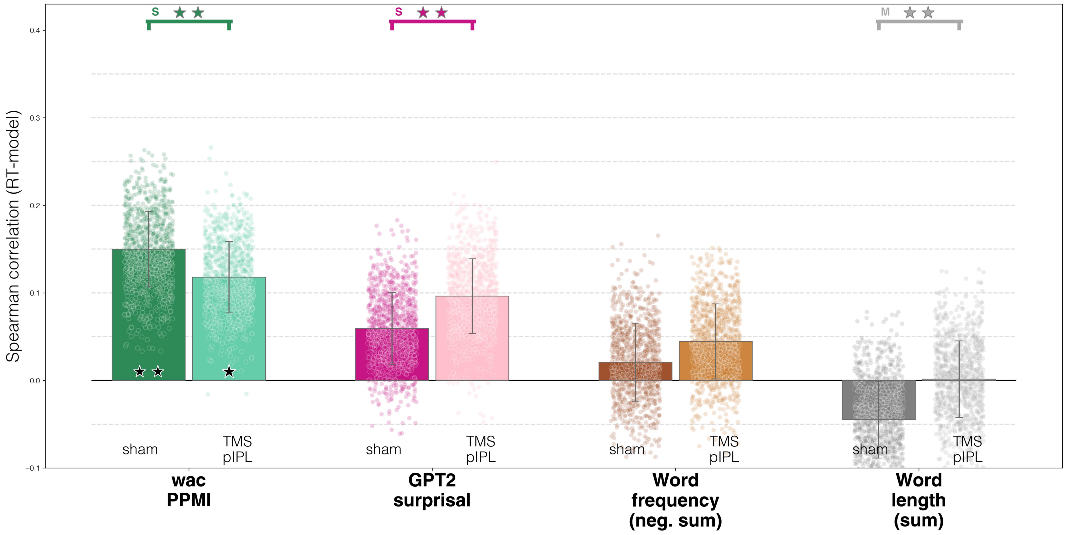

**Figure C11**

*Additional results for* ***quantity semantic priming, no residualization***

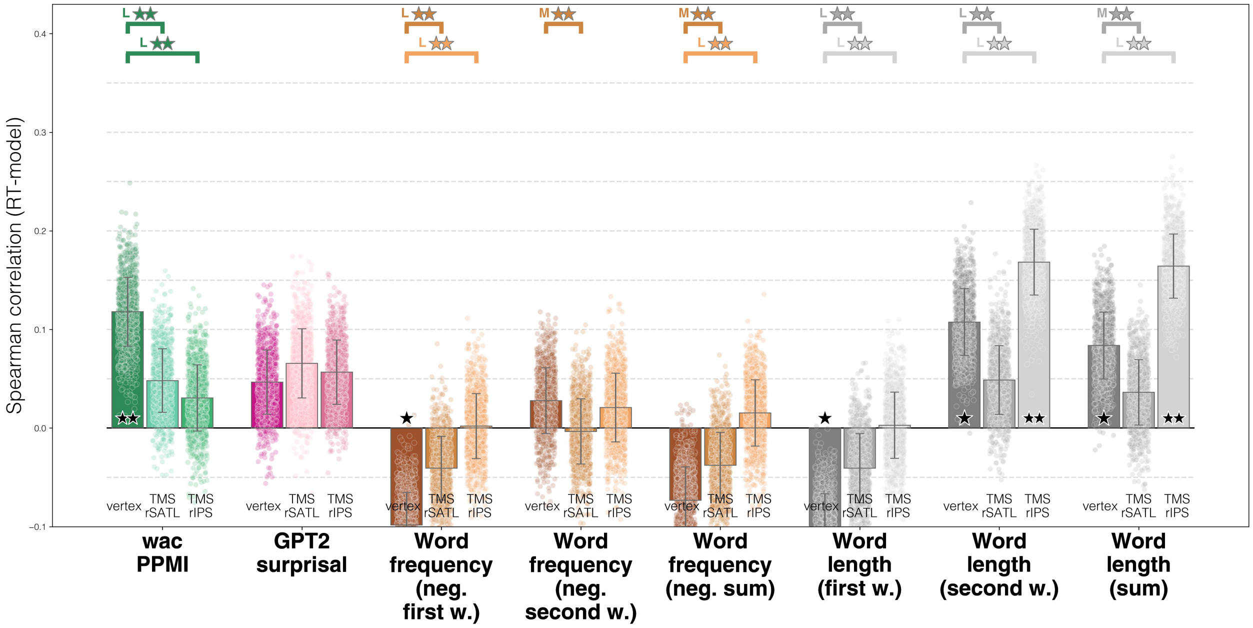


**Figure C12**

*Additional results for* ***quantity semantic priming, after residualization*** *of word length and frequency*

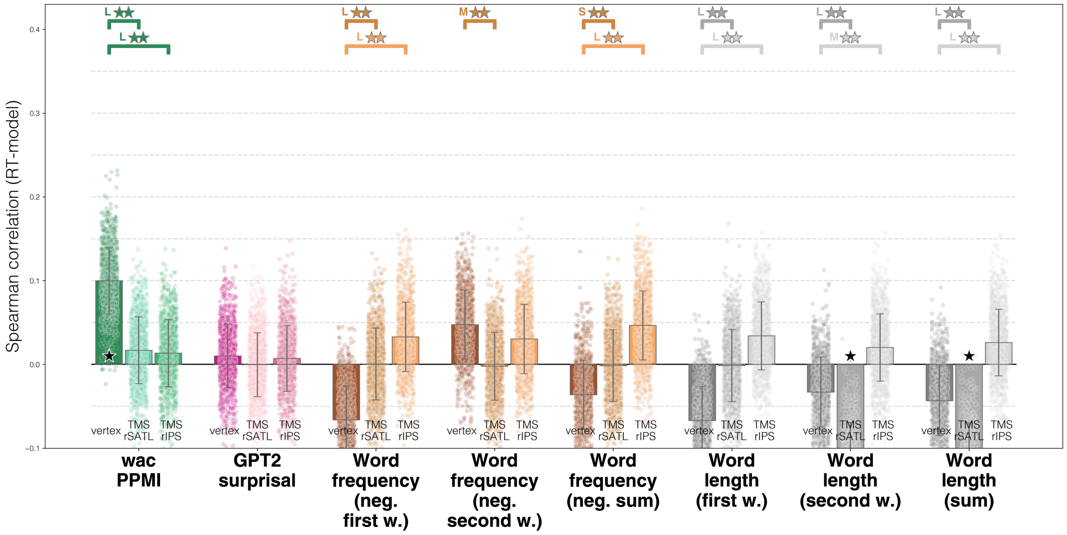

#### Figure C13

*Additional results for* ***social semantic priming, no residualization***

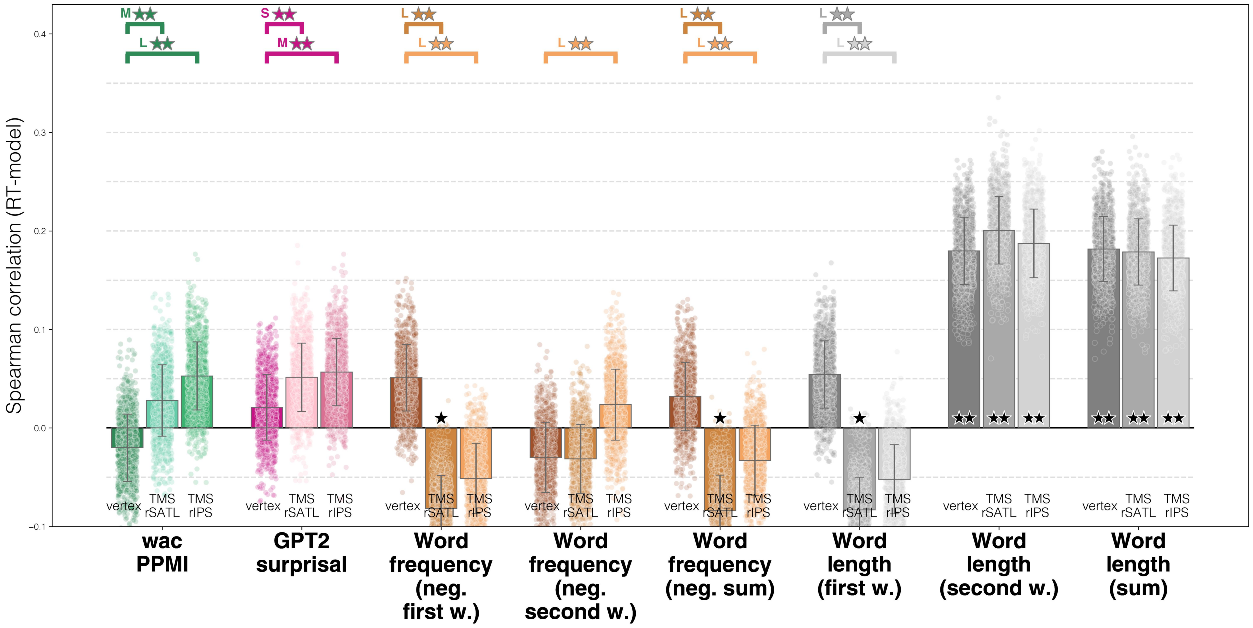


#### Figure C14

*Additional results for* ***social semantic priming, after residualization*** *of word length and frequency*

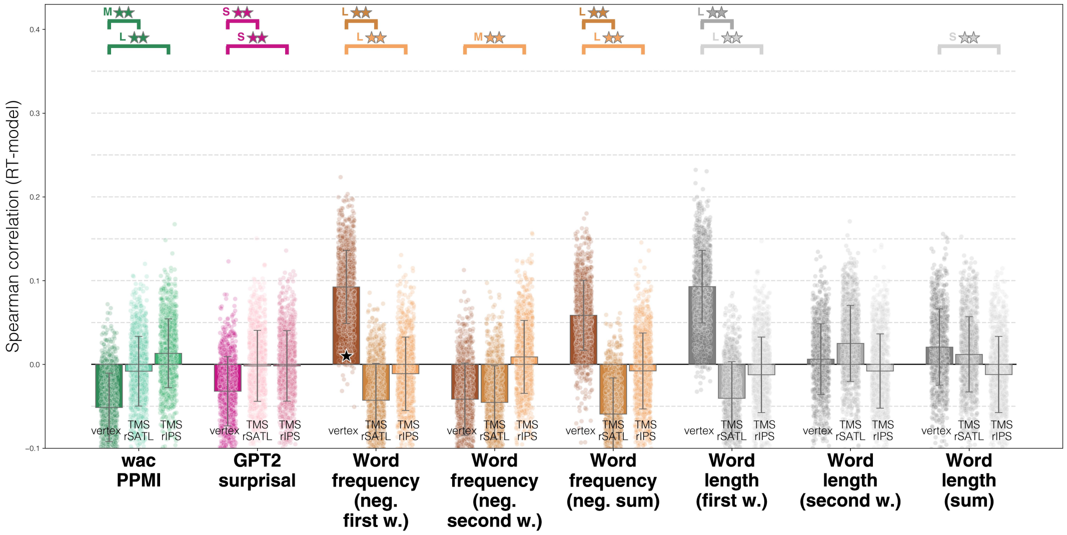

**Figure C15**

*Full stats against null hypothesis #1*

wac_corrected_baseline_p-vals

| approach | dataset | condition | model | ci_min | ci_max | perms_avg | t_value | raw_p | fdr_corrected_p |
|---|---|---|---|---|---|---|---|---|---|
| **residualize** | action feature judgment | sham | wac$PPMI | 0.13273035005277492 | 0.13810149630542504 | 0.13541592317909998 | 3.1252792 | 0.007992007992007992 | 0.01658881658881659 |
| **residualize** | action feature judgment | TMS$pIPL | wac$PPMI | 0.1509876363664787 | 0.15620928676952126 | 0.15359846156799997 | 3.646407 | 0.001998001998001998 | 0.004434871486894609 |
| **residualize** | action feature judgment | sham | GPT2$surprisal | -0.03197886104601806 | -0.026809831353341935 | -0.02939434619968 | -0.7049216 | 0.48951048951048953 | 0.635040635040635 |
| **residualize** | action feature judgment | TMS$pIPL | GPT2$surprisal | -0.014826750295980295 | -0.009441018812075704 | -0.012133884554028 | -0.2792809 | 0.7812187812187812 | 0.8797302404340527 |
| **residualize** | action feature judgment | sham | Word$frequency$(neg. sum) | -0.03382661740840826 | -0.028671344492551737 | -0.03124898095048 | -0.7513984 | 0.45554445554445555 | 0.6032036928588652 |
| **residualize** | action feature judgment | TMS$pIPL | Word$frequency$(neg. sum) | -0.02811532092368997 | -0.022572096280310025 | -0.025343708601999998 | -0.5667529 | 0.5794205794205795 | 0.7200566423867395 |
| **residualize** | action feature judgment | sham | Word$length$(sum) | 0.01272262706054926 | 0.01798966235871874 | 0.015356144709634 | 0.36141154 | 0.7272727272727273 | 0.8361458900381057 |
| **residualize** | action feature judgment | TMS$pIPL | Word$length$(sum) | 0.004325394698081103 | 0.009617005630650897 | 0.006971200164366 | 0.16330734 | 0.8771228771228772 | 0.9382038574239131 |
| **residualize** | sound feature judgment | sham | wac$PPMI | 0.1472392683415169 | 0.1525868087224831 | 0.149913038532 | 3.475133 | 0.001998001998001998 | 0.004434871486894609 |
| **residualize** | sound feature judgment | TMS$pIPL | wac$PPMI | 0.11531709208800323 | 0.12038129273999679 | 0.117849192414 | 2.8847077 | 0.005994005994005994 | 0.012577586348078151 |
| **residualize** | sound feature judgment | sham | GPT2$surprisal | 0.05685610062960739 | 0.061949899363288624 | 0.059402999996448005 | 1.4456152 | 0.15984015984015984 | 0.26118562288775055 |
| **residualize** | sound feature judgment | TMS$pIPL | GPT2$surprisal | 0.093618348575507 | 0.09891634377929301 | 0.0962673461774 | 2.2524414 | 0.03196803196803197 | 0.060175118998648414 |
| **residualize** | sound feature judgment | sham | Word$frequency$(neg. sum) | 0.017996978983021786 | 0.023523695494218212 | 0.02076033723862 | 0.46564326 | 0.6713286713286714 | 0.7932006455083379 |
| **residualize** | sound feature judgment | TMS$pIPL | Word$frequency$(neg. sum) | 0.0419279448459499 | 0.04723615442189008 | 0.04458204963391999 | 1.0411134 | 0.3116883116883117 | 0.45682561713096065 |
| **residualize** | sound feature judgment | sham | Word$length$(sum) | -0.0474856123218358 | -0.04204492715554419 | -0.044765269738689994 | -1.0199379 | 0.2997002997002997 | 0.4460655623446321 |
| **residualize** | sound feature judgment | TMS$pIPL | Word$length$(sum) | -0.0013294755257670435 | 0.004098274853055042 | 0.0013843996636439995 | 0.03161751 | 0.961038961038961 | 0.9928894634776987 |
| **residualize** | semantic production | vertex | wac$PPMI | 0.23867014488767593 | 0.24392662732032402 | 0.24129838610399998 | 5.690432 | 0.001998001998001998 | 0.004434871486894609 |
| **residualize** | semantic production | TMS$pMTG | wac$PPMI | 0.27714958980993226 | 0.2823709095700678 | 0.27976024969 | 6.6418915 | 0.001998001998001998 | 0.004434871486894609 |
| **residualize** | semantic production | vertex | GPT2$surprisal | 0.2192624519882775 | 0.22333151538972248 | 0.221296983689 | 6.741664 | 0.001998001998001998 | 0.004434871486894609 |
| **residualize** | semantic production | TMS$pMTG | GPT2$surprisal | 0.24519890940392397 | 0.249393639276076 | 0.24729627433999998 | 7.3080187 | 0.001998001998001998 | 0.004434871486894609 |
| **residualize** | semantic production | vertex | Word$frequency$(neg.$visual) | 0.011436118250680966 | 0.01729718585689904 | 0.014366652053790003 | 0.30385396 | 0.7592407592407593 | 0.8625693832794424 |
| **residualize** | semantic production | TMS$pMTG | Word$frequency$(neg.$visual) | -0.004073407553298247 | 0.001508356384014446 | -0.0012825255846419003 | -0.028482664 | 0.997002997002997 | 0.999000999000999 |
| **residualize** | semantic production | vertex | Word$frequency$(neg.$uttered) | -0.0008915719090691189 | 0.004994741391407119 | 0.002051584741169 | 0.043204814 | 0.999000999000999 | 0.999000999000999 |
| **residualize** | semantic production | TMS$pMTG | Word$frequency$(neg.$uttered) | 0.010044247406808003 | 0.015906947446931996 | 0.01297559742687 | 0.2743568 | 0.7872127872127872 | 0.8813111087163565 |
| **residualize** | semantic production | vertex | Word$frequency$(neg. sum) | -0.019348039555024257 | -0.013328129433779739 | -0.016338084494401998 | -0.33643195 | 0.7132867132867133 | 0.8271368271368271 |
| **residualize** | semantic production | TMS$pMTG | Word$frequency$(neg. sum) | -0.027831845827637952 | -0.022056173217266044 | -0.024944009522451998 | -0.5353647 | 0.5754245754245755 | 0.7197493060685244 |
| **residualize** | semantic production | vertex | Word$length$(visual) | 0.017328107745561563 | 0.023017240116498434 | 0.020172673931029998 | 0.43954515 | 0.6413586413586414 | 0.7744708122066614 |
| **residualize** | semantic production | TMS$pMTG | Word$length$(visual) | -0.014964218441954962 | -0.009283221456625036 | -0.012123719949289999 | -0.26454368 | 0.7992007992007992 | 0.889191733122369 |
| **residualize** | semantic production | vertex | Word$length$(uttered) | -0.003915274594184674 | 0.0019015427392846745 | -0.0010068659274499998 | -0.021457165 | 0.967032967032967 | 0.9928894634776987 |
| **residualize** | semantic production | TMS$pMTG | Word$length$(uttered) | -0.0034460588055185816 | 0.002288989059288581 | -0.0005785348731150001 | -0.012504853 | 0.999000999000999 | 0.999000999000999 |
| **residualize** | semantic production | vertex | Word$length$(sum) | 0.016320592377508415 | 0.02203599284968158 | 0.019178292613594997 | 0.4159579 | 0.6813186813186813 | 0.802534888424459 |
| **residualize** | semantic production | TMS$pMTG | Word$length$(sum) | 0.0005727890597155994 | 0.006233046053084401 | 0.0034029175564000004 | 0.07452489 | 0.9590409590409591 | 0.9928894634776987 |
| **residualize** | semantic production | vertex | wac$PPMI | 0.11916446935949149 | 0.12401763117272852 | 0.12159105026611 | 3.1057243 | 0.003996003996003996 | 0.008524808524808525 |
| **residualize** | semantic production | TMS$pIFG | wac$PPMI | 0.17744067887496587 | 0.18246170522903413 | 0.179951192052 | 4.4427137 | 0.001998001998001998 | 0.004434871486894609 |
| **residualize** | semantic production | TMS$aIFG | wac$PPMI | 0.17471296155728303 | 0.17950762451271698 | 0.177110293035 | 4.579012 | 0.001998001998001998 | 0.004434871486894609 |
| **residualize** | semantic production | vertex | GPT2$surprisal | 0.02364578712804944 | 0.02844877757446656 | 0.026047282351258 | 0.6722593 | 0.4775224775224775 | 0.6284500482452701 |
| **residualize** | semantic production | TMS$pIFG | GPT2$surprisal | 0.036960763721558314 | 0.04168738997197768 | 0.039324076846767995 | 1.0313199 | 0.31368631368631367 | 0.4580058724545416 |
| **residualize** | semantic production | TMS$aIFG | GPT2$surprisal | 0.028653212906043748 | 0.03362325744873626 | 0.031138235177390002 | 0.77664006 | 0.4115884115884116 | 0.56245533825605 |
| **residualize** | semantic production | vertex | Word$frequency$(neg.$visual) | 0.011580702452916062 | 0.017565297510363932 | 0.014572999981639998 | 0.30185634 | 0.7512487512487512 | 0.8585700014271442 |
| **residualize** | semantic production | TMS$pIFG | Word$frequency$(neg.$visual) | 0.02699660777339641 | 0.033028776618959595 | 0.030012692196178 | 0.61676186 | 0.5714285714285714 | 0.7170868347338936 |
| **residualize** | picture naming w/ interf | TMS$aIFG | Word$frequency$(neg.$visual) | -0.006670940764647392 | -0.0006305977429686069 | -0.0036507692538079992 | -0.07492191 | 0.9630369630369631 | 0.9928894634776987 |
| **residualize** | picture naming w/ interf | vertex | Word$frequency$(neg.$uttered) | 0.010758929497687742 | 0.016875091339284255 | 0.013817010418485999 | 0.2800407 | 0.7772227772227772 | 0.8778045483927838 |
| **residualize** | picture naming w/ interf | TMS$pIFG | Word$frequency$(neg.$uttered) | 0.0493105962767819 | 0.055407993671608105 | 0.052359294974195 | 1.0644747 | 0.30569430569430567 | 0.4514869745638976 |
| **residualize** | picture naming w/ interf | TMS$aIFG | Word$frequency$(neg.$uttered) | 0.01214727942510075 | 0.017947868010407254 | 0.015047573717754002 | 0.32157367 | 0.7692307692307693 | 0.8713410483321988 |
| **residualize** | picture naming w/ interf | vertex | Word$frequency$(neg. sum) | 0.00384178150359154 | 0.00985523392872846 | 0.00684850771616 | 0.14117512 | 0.8811188811188811 | 0.9398601398601399 |
| **residualize** | picture naming w/ interf | TMS$pIFG | Word$frequency$(neg. sum) | 0.04028361456872661 | 0.04627169292149339 | 0.04327765374511 | 0.8959057 | 0.3436563436563437 | 0.49610539836103745 |
| **residualize** | picture naming w/ interf | TMS$aIFG | Word$frequency$(neg. sum) | -0.015815705051216777 | -0.009983728479323224 | -0.01289971676527 | -0.2741893 | 0.8091908091908092 | 0.8951048951048951 |
| **residualize** | picture naming w/ interf | vertex | Word$length$(visual) | -0.0005054527004422792 | 0.005042607364558279 | 0.002268577332058 | 0.05068722 | 0.977022977022977 | 0.9971990873847638 |
| **residualize** | picture naming w/ interf | TMS$pIFG | Word$length$(visual) | -0.008980651595468223 | -0.0029404161712717784 | -0.00596053388337 | -0.122325614 | 0.8891108891108891 | 0.9431452525375177 |
| **residualize** | picture naming w/ interf | TMS$aIFG | Word$length$(visual) | 0.01766255230190745 | 0.023711842077332554 | 0.02068719718962 | 0.4239194 | 0.6713286713286714 | 0.7932006455083379 |
| **residualize** | picture naming w/ interf | vertex | Word$length$(uttered) | -0.0018225136312936853 | 0.0037086887251536853 | 0.00094308754693 | 0.02113579 | 0.983016983016983 | 0.9980099056616207 |
| **residualize** | picture naming w/ interf | TMS$pIFG | Word$length$(uttered) | -0.015332912442824625 | -0.010019063465792774 | -0.0126759879543087 | -0.29570502 | 0.7172827172827173 | 0.8271368271368271 |
| **residualize** | picture naming w/ interf | TMS$aIFG | Word$length$(uttered) | -0.05810153134810279 | -0.05286834076873521 | -0.055484936058419 | -1.3143003 | 0.17182817182817184 | 0.2772353696723445 |
| **residualize** | picture naming w/ interf | vertex | Word$length$(sum) | 0.005485512408303747 | 0.011172503912596252 | 0.00832900816045 | 0.1815502 | 0.8691308691308691 | 0.9374894768153195 |
| **residualize** | picture naming w/ interf | TMS$pIFG | Word$length$(sum) | -0.029281450321697156 | -0.023689109967482844 | -0.02648528014459 | -0.5870797 | 0.5774225774225774 | 0.7199034731502264 |
| **residualize** | picture naming w/ interf | TMS$aIFG | Word$length$(sum) | -0.02730895082574263 | -0.021676174598797376 | -0.024492562712270002 | -0.5390112 | 0.6053946053946054 | 0.7427205382476949 |
| **residualize** | semantic relatedness judg. | vertex | wac$PPMI | 0.09737563774312052 | 0.10227828516239948 | 0.09982696145276 | 2.5240808 | 0.015984015984015984 | 0.031638464628155345 |
| **residualize** | semantic relatedness judg. | TMS$rSATL | wac$PPMI | 0.014400392862609222 | 0.01933445320523078 | 0.01686742303392 | 0.42377016 | 0.6593406593406593 | 0.7862944508907243 |
| **residualize** | semantic relatedness judg. | TMS$rIPS | wac$PPMI | 0.010897132950253135 | 0.015871482315822867 | 0.013384307633038 | 0.3335383 | 0.7152847152847153 | 0.8271368271368271 |
| **residualize** | semantic relatedness judg. | vertex | GPT2$surprisal | 0.0076761719163537115 | 0.01238744359043029 | 0.010031807753392 | 0.2639533 | 0.8011988011988012 | 0.889191733122369 |
| **residualize** | semantic relatedness judg. | TMS$rSATL | GPT2$surprisal | -0.0027069789605097894 | 0.0020183635087897894 | -0.00034430772586 | -0.009032329 | 0.985014985014985 | 0.9980099056616207 |
| **residualize** | semantic relatedness judg. | TMS$rIPS | GPT2$surprisal | 0.004583020778536543 | 0.009433748487355455 | 0.007008384632945999 | 0.17910063 | 0.8771228771228772 | 0.9382038574239131 |
| **residualize** | semantic relatedness judg. | vertex | Word$frequency$(neg.$first w.) | -0.06873087309522996 | -0.06370768047597 | -0.06621927678559998 | -1.6341453 | 0.1038961038961039 | 0.18052535699594524 |
| **residualize** | semantic relatedness judg. | TMS$rSATL | Word$frequency$(neg.$first w.) | -0.0020415911329527145 | 0.0032769870535927148 | 0.0006176979603200001 | 0.014396821 | 0.9790209790209791 | 0.9971990873847638 |
| **residualize** | semantic relatedness judg. | TMS$rIPS | Word$frequency$(neg.$first w.) | 0.03027162105222912 | 0.035416841223050874 | 0.03284423113764 | 0.79130006 | 0.43156843156843155 | 0.5814816762185183 |
| **residualize** | semantic relatedness judg. | vertex | Word$frequency$(neg.$second v | 0.044857933534515795 | 0.0499670138814402 | 0.047412473707978 | 1.1503658 | 0.25374625374625376 | 0.3897542457542458 |
| **residualize** | semantic relatedness judg. | TMS$rSATL | Word$frequency$(neg.$second v | -0.0047728510580919295 | 0.0002573039956799278 | -0.002257773531206001 | -0.055639736 | 0.951048951048951 | 0.9928894634776987 |
| **residualize** | semantic relatedness judg. | TMS$rIPS | Word$frequency$(neg.$second v | 0.027775293169256567 | 0.032900675227443436 | 0.03033798419835 | 0.73374736 | 0.47952047952047955 | 0.6284500482452701 |
| **residualize** | semantic relatedness judg. | vertex | Word$frequency$(neg. sum) | -0.0388932219774186 | -0.03377785479734139 | -0.03633553838738 | -0.8805232 | 0.3516483516483517 | 0.5019812900853793 |
| **residualize** | semantic relatedness judg. | TMS$rSATL | Word$frequency$(neg. sum) | -0.0041436231559999395 | 0.0011637768989679388 | -0.0014899231285160006 | -0.034799106 | 0.993006993006993 | 0.999000999000999 |
| **residualize** | semantic relatedness judg. | TMS$rIPS | Word$frequency$(neg. sum) | 0.04400295302741514 | 0.04909796996778486 | 0.0465504614976 | 1.1325684 | 0.25374625374625376 | 0.3897542457542458 |
| **residualize** | semantic relatedness judg. | vertex | Word$length$(first w.) | -0.0695981500507601 | -0.0645523989035499 | -0.067075274477155 | -1.6478691 | 0.0999000999000999 | 0.1751672984549697 |
| **residualize** | quantity priming | TMS$rSATL | Word$length$(first w.) | -0.004045863213305059 | 0.0012902861500777585 | -0.00137778853161365 | -0.03200669 | 0.993006993006993 | 0.999000999000999 |
| **residualize** | quantity priming | TMS$rIPS | Word$length$(first w.) | 0.03166836696353712 | 0.03669373369844888 | 0.034181050330993 | 0.84314793 | 0.4095904095904096 | 0.56245533825605 |
| **residualize** | quantity priming | vertex | Word$length$(second w.) | -0.035568396223889964 | -0.03040044897112203 | -0.032984422597505995 | -0.7911828 | 0.44955044955044954 | 0.6014891032312635 |
| **residualize** | quantity priming | TMS$rSATL | Word$length$(second w.) | -0.11381632711703205 | -0.10861082603396793 | -0.11121357657549999 | -2.6483865 | 0.00999000999000999 | 0.02062453675356901 |
| **residualize** | quantity priming | TMS$rIPS | Word$length$(second w.) | 0.017676552635144276 | 0.022661053107395725 | 0.02016880287127 | 0.501585 | 0.6173826173826173 | 0.7526188097616668 |
| **residualize** | quantity priming | vertex | Word$length$(sum) | -0.04590751501987191 | -0.040736166670720074 | -0.04332184084529599 | -1.0384585 | 0.3096903096903097 | 0.45563631770528323 |
| **residualize** | quantity priming | TMS$rSATL | Word$length$(sum) | -0.11241354504449201 | -0.10742886124090799 | -0.1099212031427 | -2.7335682 | 0.005994005994005994 | 0.012577586348078151 |
| **residualize** | quantity priming | TMS$rIPS | Word$length$(sum) | 0.023668688321565247 | 0.028598383346587757 | 0.0261335358340765 | 0.65714955 | 0.5074925074925075 | 0.6517629527662973 |
| **residualize** | quantity priming | vertex | wac$PPMI | -0.05392054309795559 | -0.048855302952688404 | -0.051387923025321997 | -1.2576133 | 0.2177822177822178 | 0.33995273019663264 |
| **residualize** | quantity priming | TMS$rSATL | wac$PPMI | -0.01075317475123462 | -0.005595055812701381 | -0.008174115281968 | -0.1964425 | 0.8571428571428571 | 0.9297820823244551 |

**Figure C16**

*Full stats against null hypothesis #2*

| | | | | | | | | | |
|---|---|---|---|---|---|---|---|---|---|
| **residualize** | quantity priming | TMS$rIPS | wac$PPMI | 0.010735788363755591 | 0.01582828853898041 | 0.013282038451368 | 0.32331043 | 0.7572427572427572 | 0.8625693832794424 |
| **residualize** | quantity priming | vertex | GPT2$surprisal | -0.03459309716719036 | -0.029450056653521628 | -0.032021576910355995 | -0.7718072 | 0.4515484515484515 | 0.602064602064602 |
| **residualize** | quantity priming | TMS$rSATL | GPT2$surprisal | -0.004360061788224574 | 0.0008922155685685755 | -0.0017339231098279996 | -0.040923078 | 0.961038961038961 | 0.9928894634776987 |
| **residualize** | quantity priming | TMS$rIPS | GPT2$surprisal | -0.004453116986822619 | 0.0007661939791146195 | -0.0018434615038539998 | -0.043783147 | 0.967032967032967 | 0.9928894634776987 |
| **residualize** | quantity priming | vertex | Word$frequency$(neg.$first w.) | 0.08976844546507684 | 0.09517497189416316 | 0.09247170867962 | 2.120199 | 0.025974025974025976 | 0.04937636620804938 |
| **residualize** | quantity priming | TMS$rSATL | Word$frequency$(neg.$first w.) | -0.04541254160069664 | -0.039972244801283356 | -0.04269239320099 | -0.9727785 | 0.35564435564435565 | 0.5058053058053058 |
| **residualize** | quantity priming | TMS$rIPS | Word$frequency$(neg.$first w.) | -0.013896473592703424 | -0.008455554297984575 | -0.011176013945344 | -0.2546248 | 0.8151848151848152 | 0.8969368740142379 |
| **residualize** | quantity priming | vertex | Word$frequency$(neg.$second w | -0.04428675632599907 | -0.03894128523395293 | -0.041614020779976 | -0.9650277 | 0.3336663336663337 | 0.48533284896921264 |
| **residualize** | quantity priming | TMS$rSATL | Word$frequency$(neg.$second w | -0.048014712672837634 | -0.04258715717014236 | -0.045300934921489996 | -1.0346392 | 0.3016983016983017 | 0.447305590162733 |
| **residualize** | quantity priming | TMS$rIPS | Word$frequency$(neg.$second w | 0.006257972262176051 | 0.011656434421587949 | 0.008957203341882 | 0.20567827 | 0.8471528471528471 | 0.9241667423485606 |
| **residualize** | quantity priming | vertex | Word$frequency$(neg. sum) | 0.056043131963376534 | 0.06121732956774747 | 0.058630230765562 | 1.4046388 | 0.16183816183816183 | 0.2633298904485345 |
| **residualize** | quantity priming | TMS$rSATL | Word$frequency$(neg. sum) | -0.062034361174638425 | -0.056683946601697566 | -0.059359153888167995 | -1.3752649 | 0.1918081918081918 | 0.30561969151180773 |
| **residualize** | quantity priming | TMS$rIPS | Word$frequency$(neg. sum) | -0.010568781516001113 | -0.004946602948378887 | -0.00775769223219 | -0.1710464 | 0.8891108891108891 | 0.9431452525375177 |
| **residualize** | quantity priming | vertex | Word$length$(first w.) | 0.09017197883279522 | 0.0955495910938048 | 0.09286078496330001 | 2.1405675 | 0.03796203796203796 | 0.0700837623914547 |
| **residualize** | social priming | TMS$rSATL | Word$length$(first w.) | -0.04319361334143902 | -0.03777079582230096 | -0.04048220458186999 | -0.9253909 | 0.3696303696303696 | 0.5180221238615399 |
| **residualize** | social priming | TMS$rIPS | Word$length$(first w.) | -0.015205148430231876 | -0.009617150304028127 | -0.012411149367130002 | -0.2753226 | 0.7912087912087912 | 0.8832098134423715 |
| **residualize** | social priming | vertex | Word$length$(second w.) | 0.00362653651972273 | 0.00886891721709927 | 0.006247726868411 | 0.14773369 | 0.8671328671328671 | 0.9374894768153195 |
| **residualize** | social priming | TMS$rSATL | Word$length$(second w.) | 0.02220741338106162 | 0.02783495288933838 | 0.0250211831352 | 0.551157 | 0.5914085914085914 | 0.7325835454867713 |
| **residualize** | social priming | TMS$rIPS | Word$length$(second w.) | -0.010679999479707027 | -0.005166970014772974 | -0.00792348474724 | -0.17816073 | 0.8431568431568431 | 0.9224279993510762 |
| **residualize** | social priming | vertex | Word$length$(sum) | 0.017882027883683115 | 0.023521781322476884 | 0.02070190460308 | 0.45502603 | 0.6513486513486514 | 0.779183433389041 |
| **residualize** | social priming | TMS$rSATL | Word$length$(sum) | 0.009242823489074712 | 0.014818820570767291 | 0.012030822029921001 | 0.26746002 | 0.8211788211788211 | 0.9009504780933352 |
| **residualize** | social priming | TMS$rIPS | Word$length$(sum) | -0.014917739523594238 | -0.009268069568337760 | -0.012092904545966 | -0.26533446 | 0.8111888111888111 | 0.8951048951048951 |
| **residualize** | social priming | vertex | wac$PPMI | 0.10282989340095816 | 0.10886579906552182 | 0.10584784623323999 | 2.1738305 | 0.029970029970029972 | 0.056692076396509895 |
| **residualize** | social priming | TMS$rCereb | wac$PPMI | 0.045590302823483725 | 0.05153662042293228 | 0.048563461623208 | 1.0123894 | 0.2957042957042957 | 0.4418305429978581 |
| **residualize** | social priming | vertex | GPT2$surprisal | 0.0864007198157467 | 0.0922494340303333 | 0.08932507692304 | 1.893211 | 0.07192807192807193 | 0.12846688195525405 |
| **residualize** | social priming | TMS$rCereb | GPT2$surprisal | 0.054013835220523373 | 0.060110472506096635 | 0.057062153863310004 | 1.1602294 | 0.26573426573426573 | 0.4033278973990437 |
| **residualize** | social priming | vertex | Word$frequency$(neg.$first w.) | -0.02000060958391925 | -0.013363741964010752 | -0.016682175773965 | -0.31158432 | 0.7872127872127872 | 0.8813111087163565 |
| **residualize** | social priming | TMS$rCereb | Word$frequency$(neg.$first w.) | -0.02204485560365208 | -0.01541962975425792 | -0.018732242678955 | -0.35048962 | 0.6953046953046953 | 0.8140152530396433 |
| **residualize** | social priming | vertex | Word$frequency$(neg.$second w | -0.0003419037186107093 | 0.006456576705150708 | 0.0030573364932699994 | 0.05574648 | 0.955044955044955 | 0.9928894634776987 |
| **residualize** | social priming | TMS$rCereb | Word$frequency$(neg.$second w | -0.004656304271186223 | 0.002511736561078223 | -0.001072283855054 | -0.018543664 | 0.971028971028971 | 0.9943336663336663 |
| **residualize** | social priming | vertex | Word$frequency$(neg. sum) | -0.023988342602234335 | -0.01734296508359366 | -0.020665653842913998 | -0.38549215 | 0.7072927072927073 | 0.825356461720098 |
| **residualize** | social priming | TMS$rCereb | Word$frequency$(neg. sum) | -0.02907396646149088 | -0.022438264404009124 | -0.025756115432750002 | -0.48114896 | 0.6193806193806194 | 0.7526650564625249 |
| **residualize** | social priming | vertex | Word$length$(first w.) | -0.03135834084930286 | -0.024805142509397134 | -0.02808174167935 | -0.53119844 | 0.5934065934065934 | 0.7326949577753435 |
| **residualize** | social priming | TMS$rCereb | Word$length$(first w.) | -0.04643848054598549 | -0.039353903034594524 | -0.04289619179029001 | -0.75056934 | 0.45554445554445555 | 0.6032036928588652 |
| **residualize** | social priming | vertex | Word$length$(second w.) | -0.0029467557448207226 | 0.0033339304977367226 | 0.00019358737645800006 | 0.0038208135 | 0.955044955044955 | 0.9928894634776987 |
| **residualize** | social priming | TMS$rCereb | Word$length$(second w.) | -0.021431047930577846 | -0.014712926201622158 | -0.0180719870661 | -0.33346027 | 0.7312687312687313 | 0.8382304262901278 |
| **residualize** | social priming | vertex | Word$length$(sum) | -0.02611899407549437 | -0.019741682523043234 | -0.022930338299268802 | -0.44571668 | 0.6413586413586414 | 0.7744708122066614 |
| **residualize** | social priming | TMS$rCereb | Word$length$(sum) | -0.04897628056527877 | -0.04229328641266123 | -0.04563478348897 | -0.84646887 | 0.3916083916083916 | 0.5428795031683119 |
| **bootstrap** | action feature judgment | sham | wac$PPMI | 0.13621006057939017 | 0.1415787173126098 | 0.13889438894599998 | 3.2070453 | 0.001998001998001998 | 0.004434871486894609 |
| **bootstrap** | action feature judgment | TMS$pIPL | wac$PPMI | 0.15132528498928258 | 0.15647269266771743 | 0.1538989888285 | 3.7062376 | 0.001998001998001998 | 0.004434871486894609 |
| **bootstrap** | action feature judgment | sham | GPT2$surprisal | -0.02534921291238661 | -0.02015612248313339 | -0.02275266769776 | -0.54311585 | 0.6013986013986014 | 0.7401828940290478 |
| **bootstrap** | action feature judgment | TMS$pIPL | GPT2$surprisal | -0.0079983357321367 | -0.0029108538480733 | -0.005454594790105001 | -0.13290633 | 0.945054945054945 | 0.9928894634776987 |
| **bootstrap** | action feature judgment | sham | Word$frequency$(neg. sum) | -0.026564843616154396 | -0.021589842662397607 | -0.024077343139276 | -0.5999312 | 0.5274725274725275 | 0.6729217626227594 |
| **bootstrap** | action feature judgment | TMS$pIPL | Word$frequency$(neg. sum) | -0.019556821215321585 | -0.01442363533801842 | -0.016990228276670002 | -0.41029698 | 0.6693306693306693 | 0.7932006455083379 |
| **bootstrap** | action feature judgment | sham | Word$length$(sum) | 0.02710856372533762 | 0.03222253213814439 | 0.029665547931741004 | 0.7190853 | 0.4835164835164835 | 0.6293909480350158 |
| **bootstrap** | action feature judgment | TMS$pIPL | Word$length$(sum) | 0.01844271669757884 | 0.023419054076717162 | 0.020930885387148 | 0.52139133 | 0.6073926073926074 | 0.7427986026712141 |
| **bootstrap** | sound feature judgment | sham | wac$PPMI | 0.14791605745089853 | 0.15295766817310152 | 0.15043686281200003 | 3.6988866 | 0.001998001998001998 | 0.004434871486894609 |
| **bootstrap** | sound feature judgment | TMS$pIPL | wac$PPMI | 0.12125233207669366 | 0.12643299107150635 | 0.1238426615741 | 2.9632707 | 0.001998001998001998 | 0.004434871486894609 |
| **bootstrap** | sound feature judgment | sham | GPT2$surprisal | 0.06548176012186796 | 0.070493707187952 | 0.06798773365490998 | 1.6815515 | 0.10589410589410589 | 0.18316818316818315 |
| **bootstrap** | sound feature judgment | TMS$pIPL | GPT2$surprisal | 0.10133042466556437 | 0.10634825264983565 | 0.10383933865770001 | 2.565265 | 0.015984015984015984 | 0.031638464628155345 |
| **bootstrap** | sound feature judgment | sham | Word$frequency$(neg. sum) | 0.025446357939132972 | 0.03062670938062703 | 0.02803653365988 | 0.67088974 | 0.48151848151848153 | 0.6289220983098535 |
| **bootstrap** | sound feature judgment | TMS$pIPL | Word$frequency$(neg. sum) | 0.050323081418289096 | 0.055329827323670915 | 0.052826454370980006 | 1.3079224 | 0.21578421578421578 | 0.3382087300454647 |
| **bootstrap** | sound feature judgment | sham | Word$length$(sum) | -0.030704284848596705 | -0.025702551231083295 | -0.02820341803984 | -0.698984 | 0.4915084915084915 | 0.6354857263948173 |
| **bootstrap** | sound feature judgment | TMS$pIPL | Word$length$(sum) | 0.015846102766849463 | 0.020660731751238535 | 0.018253417259044 | 0.46996707 | 0.6453546453546454 | 0.7768532408030842 |
| **bootstrap** | semantic production | vertex | wac$PPMI | 0.1974002213814982 | 0.20146394554650182 | 0.19943208346400002 | 6.083547 | 0.001998001998001998 | 0.004434871486894609 |
| **bootstrap** | semantic production | TMS$pIFG | wac$PPMI | 0.2519363674814221 | 0.2560825630385779 | 0.25400946526 | 7.594272 | 0.001998001998001998 | 0.004434871486894609 |
| **bootstrap** | semantic production | TMS$aIFG | wac$PPMI | 0.2561078151681851 | 0.2602299920118148 | 0.25816890358999994 | 7.763603 | 0.001998001998001998 | 0.004434871486894609 |
| **bootstrap** | semantic production | vertex | GPT2$surprisal | 0.11226177595320365 | 0.11651412727879633 | 0.11438795161599999 | 3.3345497 | 0.001998001998001998 | 0.004434871486894609 |
| **bootstrap** | semantic production | TMS$pIFG | GPT2$surprisal | 0.12485166672157993 | 0.12889486424242003 | 0.12687326548199998 | 3.8898351 | 0.001998001998001998 | 0.004434871486894609 |
| **bootstrap** | semantic production | TMS$aIFG | GPT2$surprisal | 0.12844066711784885 | 0.13263438982215114 | 0.13053752847 | 3.858529 | 0.001998001998001998 | 0.004434871486894609 |
| **bootstrap** | semantic production | vertex | Word$frequency$(neg.$visual) | 0.13856070569384743 | 0.14272699104015257 | 0.140643848367 | 4.184637 | 0.001998001998001998 | 0.004434871486894609 |
| **bootstrap** | semantic production | TMS$pIFG | Word$frequency$(neg.$visual) | 0.1530134601336137 | 0.15707334819438631 | 0.155043404164 | 4.733968 | 0.001998001998001998 | 0.004434871486894609 |
| **bootstrap** | semantic production | TMS$aIFG | Word$frequency$(neg.$visual) | 0.12132379870508363 | 0.12539086531891638 | 0.123357332012 | 3.7598433 | 0.001998001998001998 | 0.004434871486894609 |
| **bootstrap** | semantic production | vertex | Word$frequency$(neg.$uttered) | 0.16908104243171063 | 0.1732603184422894 | 0.171170680437 | 5.0770845 | 0.001998001998001998 | 0.004434871486894609 |
| **bootstrap** | semantic production | TMS$pIFG | Word$frequency$(neg.$uttered) | 0.20065342357122612 | 0.20476944426877386 | 0.20271143392 | 6.1050158 | 0.001998001998001998 | 0.004434871486894609 |
| **bootstrap** | semantic production | TMS$aIFG | Word$frequency$(neg.$uttered) | 0.16769819773023967 | 0.17171494104776028 | 0.16970656938899997 | 5.2373385 | 0.001998001998001998 | 0.004434871486894609 |
| **bootstrap** | semantic production | vertex | Word$frequency$(neg. sum) | 0.1633285740074006 | 0.1673759932285994 | 0.165352283618 | 5.064285 | 0.001998001998001998 | 0.004434871486894609 |
| **bootstrap** | semantic production | TMS$pIFG | Word$frequency$(neg. sum) | 0.20377116855551203 | 0.207880892260488 | 0.20582603040800002 | 6.208315 | 0.001998001998001998 | 0.004434871486894609 |
| **bootstrap** | semantic production | TMS$aIFG | Word$frequency$(neg. sum) | 0.1448986267982632 | 0.14882739416573676 | 0.14686301048199998 | 4.6338525 | 0.001998001998001998 | 0.004434871486894609 |
| **bootstrap** | semantic production | vertex | Word$length$(visual) | 0.08723162706770204 | 0.09132765910849797 | 0.0892796430881 | 2.7019367 | 0.011988011988011988 | 0.024228403175771594 |
| **bootstrap** | semantic production | TMS$pIFG | Word$length$(visual) | 0.07515402749435976 | 0.07951117322625226 | 0.07733260036030601 | 2.2001212 | 0.025974025974025976 | 0.04937636620804938 |
| **bootstrap** | semantic production | TMS$aIFG | Word$length$(visual) | 0.09435792154913274 | 0.09872698243226723 | 0.09654245199069998 | 2.739153 | 0.003996003996003996 | 0.008524808524808525 |
| **bootstrap** | semantic production | vertex | Word$length$(uttered) | 0.16364245960691348 | 0.1677388744730865 | 0.16569066704 | 5.0139523 | 0.001998001998001998 | 0.004434871486894609 |
| **bootstrap** | semantic production | TMS$pIFG | Word$length$(uttered) | 0.14894439874903703 | 0.15312803176496298 | 0.151036215257 | 4.4752116 | 0.001998001998001998 | 0.004434871486894609 |
| **bootstrap** | semantic production | TMS$aIFG | Word$length$(uttered) | 0.11410767336875993 | 0.11808614153724006 | 0.116096907453 | 3.6173525 | 0.001998001998001998 | 0.004434871486894609 |
| **bootstrap** | semantic production | vertex | Word$length$(sum) | 0.1795829196387453 | 0.1836289299332547 | 0.181605924786 | 5.5640254 | 0.001998001998001998 | 0.004434871486894609 |
| **bootstrap** | semantic production | TMS$pIFG | Word$length$(sum) | 0.1416967471086187 | 0.1460732913273813 | 0.143885019218 | 4.075401 | 0.001998001998001998 | 0.004434871486894609 |
| **bootstrap** | semantic production | TMS$aIFG | Word$length$(sum) | 0.1407643809068969 | 0.14482116988310306 | 0.142792775395 | 4.3632483 | 0.001998001998001998 | 0.004434871486894609 |
| **bootstrap** | picture naming w/ interf | vertex | wac$PPMI | 0.2553085340648052 | 0.2604960732951948 | 0.25790230368 | 6.1628265 | 0.001998001998001998 | 0.004434871486894609 |
| **bootstrap** | picture naming w/ interf | TMS$pMTG | wac$PPMI | 0.29090685901491914 | 0.29601438866508084 | 0.29346062384 | 7.1223774 | 0.001998001998001998 | 0.004434871486894609 |
| **bootstrap** | picture naming w/ interf | vertex | GPT2$surprisal | 0.27100700955869467 | 0.27636658838930533 | 0.273686798974 | 6.3300815 | 0.001998001998001998 | 0.004434871486894609 |
| **bootstrap** | picture naming w/ interf | TMS$pMTG | GPT2$surprisal | 0.30454431234869067 | 0.3097373598313093 | 0.30714083609 | 7.3316426 | 0.001998001998001998 | 0.004434871486894609 |
| **bootstrap** | picture naming w/ interf | vertex | Word$frequency$(neg.$visual) | 0.07286970030123491 | 0.0782173930734851 | 0.07554354668736 | 1.7511243 | 0.08391608391608392 | 0.1484966646256969 |
| **bootstrap** | picture naming w/ interf | TMS$pMTG | Word$frequency$(neg.$visual) | 0.05442983308109419 | 0.0598106737283658 | 0.057120253404729995 | 1.3159096 | 0.16383616383616384 | 0.26545606292441737 |

#### Figure C17

*Full stats against null hypothesis #3*

| | | | | | | | | | |
|---|---|---|---|---|---|---|---|---|---|
| **bootstrap** | picture naming w/ interf | vertex | Word$frequency$(neg.$uttered) | 0.05576301469360768 | 0.06102835521598233 | 0.058395684954795 | 1.3748026 | 0.15984015984015984 | 0.26118562288775055 |
| **bootstrap** | picture naming w/ interf | TMS$pMTG | Word$frequency$(neg.$uttered) | 0.06379714839592542 | 0.06915061033275459 | 0.06647387936434 | 1.5392259 | 0.11388611388611389 | 0.19523333809048093 |
| **bootstrap** | picture naming w/ interf | vertex | Word$frequency$(neg. sum) | 0.04319742100426359 | 0.04867099775114442 | 0.045934209377704004 | 1.0402821 | 0.27972027972027974 | 0.41958041958041964 |
| **bootstrap** | picture naming w/ interf | TMS$pMTG | Word$frequency$(neg. sum) | 0.030434046128468197 | 0.03591524521029982 | 0.03317464566938401 | 0.7502687 | 0.43756243756243757 | 0.5874964196642519 |
| **bootstrap** | picture naming w/ interf | vertex | Word$length$(visual) | 0.043138267658765606 | 0.04835170417160038 | 0.045744985915182994 | 1.087691 | 0.26573426573426573 | 0.4033278973990437 |
| **bootstrap** | picture naming w/ interf | TMS$pMTG | Word$length$(visual) | 0.01346541522356407 | 0.018722679007886527 | 0.016094047115725298 | 0.37948236 | 0.7092907092907093 | 0.825356461720098 |
| **bootstrap** | picture naming w/ interf | vertex | Word$length$(uttered) | 0.03738051985976713 | 0.04263493762141288 | 0.040007728740590005 | 0.94385517 | 0.34965034965034963 | 0.5009915457676651 |
| **bootstrap** | picture naming w/ interf | TMS$pMTG | Word$length$(uttered) | 0.033814875226985036 | 0.03903955082791097 | 0.036427213027448 | 0.86427647 | 0.42157842157842157 | 0.5720357381134766 |
| **bootstrap** | picture naming w/ interf | vertex | Word$length$(sum) | 0.05615529324459866 | 0.061480285718825335 | 0.058817789481712 | 1.369228 | 0.1838161838161838 | 0.2953364627004794 |
| **bootstrap** | picture naming w/ interf | TMS$pMTG | Word$length$(sum) | 0.03576725865448574 | 0.04114575940286025 | 0.038456509028672994 | 0.8863284 | 0.36163836163836166 | 0.5124322172292652 |
| **bootstrap** | semantic relatedness judg. | vertex | wac$PPMI | 0.21131207682388886 | 0.21688223131611112 | 0.21409715407 | 4.764636 | 0.001998001998001998 | 0.004434871486894609 |
| **bootstrap** | semantic relatedness judg. | TMS$rCereb | wac$PPMI | 0.15711172416598526 | 0.1625306607870907 | 0.15982119247653798 | 3.6560016 | 0.001998001998001998 | 0.004434871486894609 |
| **bootstrap** | semantic relatedness judg. | vertex | GPT2$surprisal | 0.23634362886023705 | 0.2415609093277629 | 0.23895226909399997 | 5.6774464 | 0.001998001998001998 | 0.004434871486894609 |
| **bootstrap** | semantic relatedness judg. | TMS$rCereb | GPT2$surprisal | 0.20992789764397318 | 0.21491879439402684 | 0.212423346019 | 5.2760596 | 0.001998001998001998 | 0.004434871486894609 |
| **bootstrap** | semantic relatedness judg. | vertex | Word$frequency$(neg.$first w.) | -0.058353923463306916 | -0.05337106004287908 | -0.055862491753093 | -1.3897202 | 0.1918081918081918 | 0.30561969151180773 |
| **bootstrap** | semantic relatedness judg. | TMS$rCereb | Word$frequency$(neg.$first w.) | -0.054514956620245456 | -0.04943568724101455 | -0.05197532193063 | -1.2684752 | 0.2137862137862138 | 0.3364504348110906 |
| **bootstrap** | semantic relatedness judg. | vertex | Word$frequency$(neg.$second v | 0.19884220590891002 | 0.20445418626709003 | 0.20164819608800003 | 4.454144 | 0.001998001998001998 | 0.004434871486894609 |
| **bootstrap** | semantic relatedness judg. | TMS$rCereb | Word$frequency$(neg.$second v | 0.2082504315052365 | 0.21342850308076353 | 0.21083946729300002 | 5.0474257 | 0.001998001998001998 | 0.004434871486894609 |
| **bootstrap** | semantic relatedness judg. | vertex | Word$frequency$(neg. sum) | 0.0719526346215247 | 0.0770733650512153 | 0.07451299983637 | 1.803791 | 0.07592407592407592 | 0.13497613497613498 |
| **bootstrap** | semantic relatedness judg. | TMS$rCereb | Word$frequency$(neg. sum) | 0.08132625116463392 | 0.0862746718459181 | 0.08380046150527601 | 2.0992582 | 0.03396603396603397 | 0.06331532545124778 |
| **bootstrap** | semantic relatedness judg. | vertex | Word$length$(first w.) | -0.026188037264938256 | -0.020952903465803745 | -0.023570470365371 | -0.55811864 | 0.5594405594405595 | 0.7043448354923765 |
| **bootstrap** | semantic relatedness judg. | TMS$rCereb | Word$length$(first w.) | -0.03121341170629253 | -0.026104597351427474 | -0.028659004528860003 | -0.6953878 | 0.47952047952047955 | 0.6284500482452701 |
| **bootstrap** | semantic relatedness judg. | vertex | Word$length$(second w.) | 0.17391811658805637 | 0.17874767622394364 | 0.176332896406 | 4.525972 | 0.001998001998001998 | 0.004434871486894609 |
| **bootstrap** | semantic relatedness judg. | TMS$rCereb | Word$length$(second w.) | 0.15907500100481298 | 0.16407125500118702 | 0.161573128003 | 4.0087657 | 0.001998001998001998 | 0.004434871486894609 |
| **bootstrap** | semantic relatedness judg. | vertex | Word$length$(sum) | 0.10492514448578369 | 0.1100897215822163 | 0.107507433034 | 2.5804164 | 0.015984015984015984 | 0.031638464628155345 |
| **bootstrap** | semantic relatedness judg. | TMS$rCereb | Word$length$(sum) | 0.09497829104969506 | 0.10007277831490492 | 0.09752553468229999 | 2.3730338 | 0.02197802197802198 | 0.04240985145507759 |
| **bootstrap** | quantity priming | vertex | wac$PPMI | 0.11577015878741309 | 0.12011026532658692 | 0.117940212057 | 3.3685856 | 0.001998001998001998 | 0.004434871486894609 |
| **bootstrap** | quantity priming | TMS$rSATL | wac$PPMI | 0.046128448675443245 | 0.05012642108110875 | 0.048127434878276 | 1.492241 | 0.12987012987012986 | 0.21872863978127133 |
| **bootstrap** | quantity priming | TMS$rIPS | wac$PPMI | 0.028385924593678346 | 0.03254748119226166 | 0.030466702892970002 | 0.9075189 | 0.36363636363636365 | 0.5133689839572192 |
| **bootstrap** | quantity priming | vertex | GPT2$surprisal | 0.04459470400882605 | 0.04862126640474594 | 0.046607985206785996 | 1.4348681 | 0.13586413586413587 | 0.2268340355296877 |
| **bootstrap** | quantity priming | TMS$rSATL | GPT2$surprisal | 0.06340568898956903 | 0.06776164468407098 | 0.06558366683682 | 1.8663725 | 0.061938061938061936 | 0.11166298490242152 |
| **bootstrap** | quantity priming | TMS$rIPS | GPT2$surprisal | 0.05466626643219036 | 0.05871351617580964 | 0.056689891304 | 1.7363276 | 0.07192807192807193 | 0.12846688195525405 |
| **bootstrap** | quantity priming | vertex | Word$frequency$(neg.$first w.) | -0.100447546995202 | -0.09635663253079799 | -0.098402089763 | -2.9817412 | 0.005994005994005994 | 0.012577586348078151 |
| **bootstrap** | quantity priming | TMS$rSATL | Word$frequency$(neg.$first w.) | -0.04273020467815999 | -0.03872752892332001 | -0.04072886680074 | -1.261357 | 0.1958041958041958 | 0.310697566895914 |
| **bootstrap** | quantity priming | TMS$rIPS | Word$frequency$(neg.$first w.) | -1.0231980171902712e-05 | 0.004069020394879903 | 0.0020293942073539998 | 0.061669696 | 0.935064935064935 | 0.9891596007298487 |
| **bootstrap** | quantity priming | vertex | Word$frequency$(neg.$second v | 0.025708681174587647 | 0.029835556287682356 | 0.027772118731135 | 0.8342068 | 0.3776223776223776 | 0.5272981563890654 |
| **bootstrap** | quantity priming | TMS$rSATL | Word$frequency$(neg.$second v | -0.005448068221897884 | -0.001351506505942116 | -0.00339978736392 | -0.10287701 | 0.8771228771228772 | 0.9382038574239131 |
| **bootstrap** | quantity priming | TMS$rIPS | Word$frequency$(neg.$second v | 0.01861492692806109 | 0.02292987966953891 | 0.0207724032988 | 0.596756 | 0.5594405594405595 | 0.7043448354923765 |
| **bootstrap** | quantity priming | vertex | Word$frequency$(neg. sum) | -0.07538326032283238 | -0.07119344068256764 | -0.07328835050270001 | -2.1683316 | 0.03396603396603397 | 0.06331532545124778 |
| **bootstrap** | quantity priming | TMS$rSATL | Word$frequency$(neg. sum) | -0.039894127590637714 | -0.03573199375234227 | -0.03781306067148999 | -1.1261904 | 0.25774225774225773 | 0.3943148484981154 |
| **bootstrap** | quantity priming | TMS$rIPS | Word$frequency$(neg. sum) | 0.013168978641918341 | 0.01733180655324366 | 0.015250392597581 | 0.45412838 | 0.6493506493506493 | 0.779183433389041 |
| **bootstrap** | quantity priming | vertex | Word$length$(first w.) | -0.10350464760585025 | -0.09921421231300977 | -0.10135942995943001 | -2.9285245 | 0.003996003996003996 | 0.008524808524808525 |
| **bootstrap** | quantity priming | TMS$rSATL | Word$length$(first w.) | -0.04283799214686121 | -0.0384935073599188 | -0.040665749753390004 | -1.1603167 | 0.23176823176823177 | 0.3588669395121008 |
| **bootstrap** | quantity priming | TMS$rIPS | Word$length$(first w.) | 0.0007030998958465835 | 0.004867913973573416 | 0.00278550693471 | 0.08290766 | 0.939060939060939 | 0.9906576939543973 |
| **bootstrap** | quantity priming | vertex | Word$length$(second w.) | 0.10539511344222544 | 0.10960032460217456 | 0.1074977190222 | 3.16882 | 0.003996003996003996 | 0.008524808524808525 |
| **bootstrap** | quantity priming | TMS$rSATL | Word$length$(second w.) | 0.046520824786889306 | 0.05083534529127068 | 0.04867808503907999 | 1.398579 | 0.13986013986013987 | 0.2324947779493234 |
| **bootstrap** | quantity priming | TMS$rIPS | Word$length$(second w.) | 0.16623918048187492 | 0.1703729145881251 | 0.168306047535 | 5.047116 | 0.001998001998001998 | 0.004434871486894609 |
| **bootstrap** | quantity priming | vertex | Word$length$(sum) | 0.0816253124610521 | 0.08583342182614791 | 0.0837293671436 | 2.466476 | 0.011988011988011988 | 0.024228403175771594 |
| **bootstrap** | quantity priming | TMS$rSATL | Word$length$(sum) | 0.03412374193967113 | 0.03823545134925686 | 0.036179596644464 | 1.0907553 | 0.27972027972027974 | 0.41958041958041964 |
| **bootstrap** | quantity priming | TMS$rIPS | Word$length$(sum) | 0.16224163092227334 | 0.16628029410972667 | 0.164260962516 | 5.041767 | 0.001998001998001998 | 0.004434871486894609 |
| **bootstrap** | social priming | vertex | wac$PPMI | -0.022154382003487627 | -0.017922613660654375 | -0.020038497832071 | -0.5869882 | 0.5574425574425574 | 0.7043448354923765 |
| **bootstrap** | social priming | TMS$rSATL | wac$PPMI | 0.025669067739546525 | 0.03015269131028548 | 0.027910879524916003 | 0.77166784 | 0.43156843156843155 | 0.5814816762185183 |
| **bootstrap** | social priming | TMS$rIPS | wac$PPMI | 0.05068382803196562 | 0.05495517285304637 | 0.052819500442505996 | 1.5329068 | 0.12987012987012986 | 0.21872863978127133 |
| **bootstrap** | social priming | vertex | GPT2$surprisal | 0.018842551241801783 | 0.022962460687842214 | 0.020902505964822 | 0.6289219 | 0.5254745254745254 | 0.6726073926073926 |
| **bootstrap** | social priming | TMS$rSATL | GPT2$surprisal | 0.049288641100972065 | 0.053585927536507934 | 0.05143728431874 | 1.483781 | 0.13186813186813187 | 0.22112385431162726 |
| **bootstrap** | social priming | TMS$rIPS | GPT2$surprisal | 0.054623634759567156 | 0.058859918040532845 | 0.05674177640005 | 1.6603665 | 0.1038961038961039 | 0.18052535699594524 |
| **bootstrap** | social priming | vertex | Word$frequency$(neg.$first w.) | 0.0489151958581522 | 0.05311688077614782 | 0.05101603831715001 | 1.5051137 | 0.14585414585414586 | 0.24141375865513798 |
| **bootstrap** | social priming | TMS$rSATL | Word$frequency$(neg.$first w.) | -0.0837179610175455 | -0.07957089775365447 | -0.08164442938559999 | -2.4404614 | 0.01998001998001998 | 0.03914452894044731 |
| **bootstrap** | social priming | TMS$rIPS | Word$frequency$(neg.$first w.) | -0.0533990620167555 | -0.04897330187834449 | -0.051186181947549994 | -1.4336758 | 0.14985014985014986 | 0.24696333709209245 |
| **bootstrap** | social priming | vertex | Word$frequency$(neg.$second v | -0.032087821229919114 | -0.027655712378258156 | -0.029871766804088637 | -0.8354809 | 0.42157842157842157 | 0.5720357381134766 |
| **bootstrap** | social priming | TMS$rSATL | Word$frequency$(neg.$second v | -0.033472262179193496 | -0.029121923257986498 | -0.03129709271859 | -0.8917989 | 0.4015984015984016 | 0.5547258496898785 |
| **bootstrap** | social priming | TMS$rIPS | Word$frequency$(neg.$second v | 0.021442150073163732 | 0.025915319140028273 | 0.023678734606596002 | 0.65618944 | 0.4975024975024975 | 0.6410770437616076 |
| **bootstrap** | social priming | vertex | Word$frequency$(neg. sum) | 0.029715448586487316 | 0.034020063195616686 | 0.031867755891052 | 0.9177054 | 0.3436563436563437 | 0.49610539836103745 |
| **bootstrap** | social priming | TMS$rSATL | Word$frequency$(neg. sum) | -0.0862364546526502 | -0.08175714929674979 | -0.0839968019747 | -2.3245459 | 0.023976023976023976 | 0.046033966033966034 |
| **bootstrap** | social priming | TMS$rIPS | Word$frequency$(neg. sum) | -0.0349903593906432 | -0.03057116316044681 | -0.032780761275545 | -0.9195214 | 0.38961038961038963 | 0.5420666290231508 |
| **bootstrap** | social priming | vertex | Word$length$(first w.) | 0.05220222914366232 | 0.05643886200099768 | 0.05432054557233 | 1.5893859 | 0.11388611388611389 | 0.19523333809048093 |
| **bootstrap** | social priming | TMS$rSATL | Word$length$(first w.) | -0.08510244143918554 | -0.08099306084549444 | -0.08304775114233999 | -2.5051723 | 0.017982017982017984 | 0.0354107431030508 |
| **bootstrap** | social priming | TMS$rIPS | Word$length$(first w.) | -0.05422886240874873 | -0.04988820171945127 | -0.0520585320641 | -1.4866959 | 0.12987012987012986 | 0.21872863978127133 |
| **bootstrap** | social priming | vertex | Word$length$(second w.) | 0.1776494292878057 | 0.18187827125619432 | 0.179763850272 | 5.26947 | 0.001998001998001998 | 0.004434871486894609 |
| **bootstrap** | social priming | TMS$rSATL | Word$length$(second w.) | 0.19858926934450166 | 0.20284361791949834 | 0.200716443632 | 5.848385 | 0.001998001998001998 | 0.004434871486894609 |
| **bootstrap** | social priming | TMS$rIPS | Word$length$(second w.) | 0.18508984053486213 | 0.18941079703113792 | 0.18725031878300002 | 5.37191 | 0.001998001998001998 | 0.004434871486894609 |
| **bootstrap** | social priming | vertex | Word$length$(sum) | 0.17964238315696063 | 0.18371858170303934 | 0.18168048242999998 | 5.525086 | 0.001998001998001998 | 0.004434871486894609 |
| **bootstrap** | social priming | TMS$rSATL | Word$length$(sum) | 0.176562443784629 | 0.18073311476737103 | 0.178647779276 | 5.309796 | 0.001998001998001998 | 0.004434871486894609 |
| **bootstrap** | social priming | TMS$rIPS | Word$length$(sum) | 0.17049436033861978 | 0.17463056826338028 | 0.17256246430100003 | 5.171661 | 0.001998001998001998 | 0.004434871486894609 |

#### Figure C18

*Full stats for comparisons among conditions #1*

wac_corrected_comparisons_p-vals

| approach | dataset | condition_one | condition_two | model | ci_min | ci_max | t_value | raw_p | fdr_corrected_p |
|---|---|---|---|---|---|---|---|---|---|
| **residualize** | action feature judg. | sham | TMS$pIPL | wac$PPMI | -0.49 | -0.36 | -0.42530397405247744 | 0.000999000999000999 | 0.0032236670892133077 |
| **residualize** | action feature judg. | sham | TMS$pIPL | GPT2$surprisal | -0.47 | -0.34 | -0.40514574011721627 | 0.000999000999000999 | 0.0032236670892133077 |
| **residualize** | action feature judg. | sham | TMS$pIPL | Word$frequency$(neg. sum) | -0.2 | -0.07 | -0.1366880688823786 | 0.000999000999000999 | 0.0032236670892133077 |
| **residualize** | action feature judg. | sham | TMS$pIPL | Word$length$(sum) | 0.13 | 0.26 | 0.19678390362103132 | 0.000999000999000999 | 0.0032236670892133077 |
| **residualize** | sound feature judg. | sham | TMS$pIPL | wac$PPMI | 0.69 | 0.83 | 0.7628344581399927 | 0.000999000999000999 | 0.0032236670892133077 |
| **residualize** | sound feature judg. | sham | TMS$pIPL | GPT2$surprisal | -0.95 | -0.81 | -0.8788828630840638 | 0.000999000999000999 | 0.0032236670892133077 |
| **residualize** | sound feature judg. | sham | TMS$pIPL | Word$frequency$(neg. sum) | -0.61 | -0.48 | -0.5447002120573357 | 0.000999000999000999 | 0.0032236670892133077 |
| **residualize** | sound feature judg. | sham | TMS$pIPL | Word$length$(sum) | -1.13 | -0.97 | -1.052204278419338 | 0.000999000999000999 | 0.0032236670892133077 |
| **residualize** | picture naming w/ interf | vertex | TMS$pMTG | wac$PPMI | -0.98 | -0.84 | -0.9096126627371884 | 0.000999000999000999 | 0.0032236670892133077 |
| **residualize** | picture naming w/ interf | vertex | TMS$pMTG | GPT2$surprisal | -0.85 | -0.71 | -0.7795261177736709 | 0.02197802197802198 | 0.04240985145507759 |
| **residualize** | picture naming w/ interf | vertex | TMS$pMTG | Word$frequency$(neg.$visual) | 0.27 | 0.4 | 0.3387875965792799 | 0.000999000999000999 | 0.0032236670892133077 |
| **residualize** | picture naming w/ interf | vertex | TMS$pMTG | Word$frequency$(neg.$uttered) | -0.29 | -0.17 | -0.2303980134916195 | 0.000999000999000999 | 0.0032236670892133077 |
| **residualize** | picture naming w/ interf | vertex | TMS$pMTG | Word$frequency$(neg. sum) | 0.12 | 0.24 | 0.18075233116268422 | 0.000999000999000999 | 0.0032236670892133077 |
| **residualize** | picture naming w/ interf | vertex | TMS$pMTG | Word$length$(visual) | 0.63 | 0.77 | 0.7038616871658667 | 0.000999000999000999 | 0.0032236670892133077 |
| **residualize** | picture naming w/ interf | vertex | TMS$pMTG | Word$length$(uttered) | -0.07 | 0.05 | -0.009187882428674849 | 0.8561438561438561 | 0.9297820823244551 |
| **residualize** | picture naming w/ interf | vertex | TMS$pMTG | Word$length$(sum) | 0.28 | 0.41 | 0.3436346061047949 | 0.000999000999000999 | 0.0032236670892133077 |
| **residualize** | semantic production | vertex | TMS$pIFG | wac$PPMI | -1.55 | -1.38 | -1.4643706309874398 | 0.000999000999000999 | 0.0032236670892133077 |
| **residualize** | semantic production | vertex | TMS$aIFG | wac$PPMI | -1.51 | -1.34 | -1.4259521372754027 | 0.000999000999000999 | 0.0032236670892133077 |
| **residualize** | semantic production | vertex | TMS$pIFG | GPT2$surprisal | -0.41 | -0.28 | -0.3452253347070806 | 0.000999000999000999 | 0.0032236670892133077 |
| **residualize** | semantic production | vertex | TMS$aIFG | GPT2$surprisal | -0.19 | -0.07 | -0.1290639600216796 | 0.04595404595404595 | 0.08443231409738586 |
| **residualize** | semantic production | vertex | TMS$pIFG | Word$frequency$(neg.$visual) | -0.38 | -0.25 | -0.31838054949682815 | 0.000999000999000999 | 0.0032236670892133077 |
| **residualize** | semantic production | vertex | TMS$aIFG | Word$frequency$(neg.$visual) | 0.31 | 0.44 | 0.3755342088837919 | 0.000999000999000999 | 0.0032236670892133077 |
| **residualize** | semantic production | vertex | TMS$pIFG | Word$frequency$(neg.$uttered) | -0.85 | -0.71 | -0.7819761303877203 | 0.000999000999000999 | 0.0032236670892133077 |
| **residualize** | semantic production | vertex | TMS$aIFG | Word$frequency$(neg.$uttered) | -0.09 | 0.04 | -0.025579537881967544 | 0.3696303696303696 | 0.5180221238615399 |
| **residualize** | semantic production | vertex | TMS$pIFG | Word$frequency$(neg. sum) | -0.82 | -0.68 | -0.7521599023523816 | 0.000999000999000999 | 0.0032236670892133077 |
| **residualize** | semantic production | vertex | TMS$aIFG | Word$frequency$(neg. sum) | 0.35 | 0.48 | 0.4130713662882039 | 0.000999000999000999 | 0.0032236670892133077 |
| **residualize** | semantic production | vertex | TMS$pIFG | Word$length$(visual) | 0.11 | 0.24 | 0.1758088955264203 | 0.000999000999000999 | 0.0032236670892133077 |
| **residualize** | semantic production | vertex | TMS$aIFG | Word$length$(visual) | -0.46 | -0.33 | -0.3931804386247016 | 0.000999000999000999 | 0.0032236670892133077 |
| **residualize** | semantic production | vertex | TMS$pIFG | Word$length$(uttered) | 0.25 | 0.37 | 0.311119780678324 | 0.000999000999000999 | 0.0032236670892133077 |
| **residualize** | semantic production | vertex | TMS$aIFG | Word$length$(uttered) | 1.21 | 1.38 | 1.2984874026878384 | 0.000999000999000999 | 0.0032236670892133077 |
| **residualize** | semantic production | vertex | TMS$pIFG | Word$length$(sum) | 0.69 | 0.84 | 0.764817248279218 | 0.000999000999000999 | 0.0032236670892133077 |
| **residualize** | semantic production | vertex | TMS$aIFG | Word$length$(sum) | 0.65 | 0.79 | 0.7184816092715638 | 0.000999000999000999 | 0.0032236670892133077 |
| **residualize** | quantity priming | vertex | TMS$rSATL | wac$PPMI | 1.98 | 2.2 | 2.0898405405307274 | 0.000999000999000999 | 0.0032236670892133077 |
| **residualize** | quantity priming | vertex | TMS$rIPS | wac$PPMI | 2.06 | 2.28 | 2.1686554059422196 | 0.000999000999000999 | 0.0032236670892133077 |
| **residualize** | quantity priming | vertex | TMS$rSATL | GPT2$surprisal | 0.21 | 0.34 | 0.27246888163510724 | 0.000999000999000999 | 0.0032236670892133077 |
| **residualize** | quantity priming | vertex | TMS$rIPS | GPT2$surprisal | 0.02 | 0.14 | 0.07834347380697466 | 0.058941058941058944 | 0.10676116336493695 |
| **residualize** | quantity priming | vertex | TMS$rSATL | Word$frequency$(neg.$first w.) | -1.69 | -1.51 | -1.6008240936438618 | 0.000999000999000999 | 0.0032236670892133077 |
| **residualize** | quantity priming | vertex | TMS$rIPS | Word$frequency$(neg.$first w.) | -2.54 | -2.29 | -2.413948965557334 | 0.000999000999000999 | 0.0032236670892133077 |
| **residualize** | quantity priming | vertex | TMS$rSATL | Word$frequency$(neg.$second w.) | 1.13 | 1.3 | 1.2138828144608174 | 0.000999000999000999 | 0.0032236670892133077 |
| **residualize** | quantity priming | vertex | TMS$rIPS | Word$frequency$(neg.$second w.) | 0.35 | 0.48 | 0.4134099794627867 | 0.000999000999000999 | 0.0032236670892133077 |
| **residualize** | quantity priming | vertex | TMS$rSATL | Word$frequency$(neg. sum) | -0.9 | -0.76 | -0.8283048720197022 | 0.000999000999000999 | 0.0032236670892133077 |
| **residualize** | quantity priming | vertex | TMS$rIPS | Word$frequency$(neg. sum) | -2.12 | -1.9 | -2.0115788797582472 | 0.000999000999000999 | 0.0032236670892133077 |
| **residualize** | quantity priming | vertex | TMS$rSATL | Word$length$(first w.) | -1.66 | -1.47 | -1.5674758097888508 | 0.000999000999000999 | 0.0032236670892133077 |
| **residualize** | quantity priming | vertex | TMS$rIPS | Word$length$(first w.) | -2.62 | -2.37 | -2.4913938474523807 | 0.000999000999000999 | 0.0032236670892133077 |
| **residualize** | quantity priming | vertex | TMS$rSATL | Word$length$(second w.) | 1.77 | 1.97 | 1.8687080432383705 | 0.000999000999000999 | 0.0032236670892133077 |
| **residualize** | quantity priming | vertex | TMS$rIPS | Word$length$(second w.) | -1.38 | -1.21 | -1.2971397788838255 | 0.000999000999000999 | 0.0032236670892133077 |
| **residualize** | quantity priming | vertex | TMS$rSATL | Word$length$(sum) | 1.53 | 1.72 | 1.6246935623913903 | 0.000999000999000999 | 0.0032236670892133077 |
| **residualize** | quantity priming | vertex | TMS$rIPS | Word$length$(sum) | -1.8 | -1.61 | -1.7033904132356292 | 0.000999000999000999 | 0.0032236670892133077 |
| **residualize** | social priming | vertex | TMS$rSATL | wac$PPMI | -1.12 | -0.97 | -1.0473933646208144 | 0.000999000999000999 | 0.0032236670892133077 |
| **residualize** | social priming | vertex | TMS$rIPS | wac$PPMI | -1.67 | -1.48 | -1.577621261624924 | 0.000999000999000999 | 0.0032236670892133077 |
| **residualize** | social priming | vertex | TMS$rSATL | GPT2$surprisal | -0.79 | -0.65 | -0.7219426133860828 | 0.000999000999000999 | 0.0032236670892133077 |
| **residualize** | social priming | vertex | TMS$rIPS | GPT2$surprisal | -0.79 | -0.65 | -0.7216403914880227 | 0.000999000999000999 | 0.0032236670892133077 |
| **residualize** | social priming | vertex | TMS$rSATL | Word$frequency$(neg.$first w.) | 2.94 | 3.24 | 3.087845034523482 | 0.000999000999000999 | 0.0032236670892133077 |
| **residualize** | social priming | vertex | TMS$rIPS | Word$frequency$(neg.$first w.) | 2.25 | 2.49 | 2.3677120789631725 | 0.000999000999000999 | 0.0032236670892133077 |
| **residualize** | social priming | vertex | TMS$rSATL | Word$frequency$(neg.$second w.) | 0.02 | 0.15 | 0.08480305598501157 | 0.2007992007992008 | 0.3173123173123173 |
| **residualize** | social priming | vertex | TMS$rIPS | Word$frequency$(neg.$second w.) | -1.25 | -1.09 | -1.1663628709822471 | 0.000999000999000999 | 0.0032236670892133077 |
| **residualize** | social priming | vertex | TMS$rSATL | Word$frequency$(neg. sum) | 2.64 | 2.91 | 2.7776326006550107 | 0.000999000999000999 | 0.0032236670892133077 |
| **residualize** | social priming | vertex | TMS$rIPS | Word$frequency$(neg. sum) | 1.43 | 1.61 | 1.5224265775232593 | 0.000999000999000999 | 0.0032236670892133077 |
| **residualize** | social priming | vertex | TMS$rSATL | Word$length$(first w.) | 2.91 | 3.21 | 3.0593146934553723 | 0.000999000999000999 | 0.0032236670892133077 |
| **residualize** | social priming | vertex | TMS$rIPS | Word$length$(first w.) | 2.26 | 2.5 | 2.378475467163757 | 0.000999000999000999 | 0.0032236670892133077 |
| **residualize** | social priming | vertex | TMS$rSATL | Word$length$(second w.) | -0.49 | -0.36 | -0.42770489952543816 | 0.000999000999000999 | 0.0032236670892133077 |
| **residualize** | social priming | vertex | TMS$rIPS | Word$length$(second w.) | 0.26 | 0.39 | 0.3263933903110363 | 0.000999000999000999 | 0.0032236670892133077 |
| **residualize** | social priming | vertex | TMS$rSATL | Word$length$(sum) | 0.13 | 0.25 | 0.19157408964895048 | 0.000999000999000999 | 0.0032236670892133077 |
| **residualize** | social priming | vertex | TMS$rIPS | Word$length$(sum) | 0.65 | 0.79 | 0.7198333683520242 | 0.000999000999000999 | 0.0032236670892133077 |
| **residualize** | semantic relatedness judg. | vertex | TMS$rCereb | wac$PPMI | 1.1 | 1.27 | 1.1846375793531807 | 0.000999000999000999 | 0.0032236670892133077 |
| **residualize** | semantic relatedness judg. | vertex | TMS$rCereb | GPT2$surprisal | 0.6 | 0.74 | 0.6691293248644998 | 0.000999000999000999 | 0.0032236670892133077 |
| **residualize** | semantic relatedness judg. | vertex | TMS$rCereb | Word$frequency$(neg.$first w.) | -0.02 | 0.1 | 0.03830492058500293 | 0.41058941058941056 | 0.56245533825605 |
| **residualize** | semantic relatedness judg. | vertex | TMS$rCereb | Word$frequency$(neg.$second w.) | 0.01 | 0.14 | 0.07324343557527857 | 0.08791208791208792 | 0.15485431999193466 |
| **residualize** | semantic relatedness judg. | vertex | TMS$rCereb | Word$frequency$(neg. sum) | 0.03 | 0.16 | 0.09497787808425667 | 0.053946053946053944 | 0.09864421292992721 |
| **residualize** | semantic relatedness judg. | vertex | TMS$rCereb | Word$length$(first w.) | 0.21 | 0.33 | 0.268974664771396 | 0.000999000999000999 | 0.0032236670892133077 |
| **residualize** | semantic relatedness judg. | vertex | TMS$rCereb | Word$length$(second w.) | 0.28 | 0.41 | 0.3480028768622444 | 0.000999000999000999 | 0.0032236670892133077 |

#### Figure C19

*Full stats for comparisons among conditions #2*

| | | | | | | | | | |
|---|---|---|---|---|---|---|---|---|---|
| **residualize** | semantic relatedness judg. | vertex | TMS$rCereb | Word$length$(sum) | 0.37 | 0.5 | 0.43066287937064746 | 0.000999000999000999 | 0.0032236670892133077 |
| **bootstrap** | action feature judg. | sham | TMS$pIPL | wac$PPMI | -0.42 | -0.29 | -0.35348737313264494 | 0.000999000999000999 | 0.0032236670892133077 |
| **bootstrap** | action feature judg. | sham | TMS$pIPL | GPT2$surprisal | -0.48 | -0.35 | -0.41692345286054266 | 0.000999000999000999 | 0.0032236670892133077 |
| **bootstrap** | action feature judg. | sham | TMS$pIPL | Word$frequency$(neg. sum) | -0.24 | -0.11 | -0.17371681359039087 | 0.000999000999000999 | 0.0032236670892133077 |
| **bootstrap** | action feature judg. | sham | TMS$pIPL | Word$length$(sum) | 0.15 | 0.28 | 0.21448662663693127 | 0.000999000999000999 | 0.0032236670892133077 |
| **bootstrap** | sound feature judg. | sham | TMS$pIPL | wac$PPMI | 0.58 | 0.71 | 0.6446117889223292 | 0.000999000999000999 | 0.0032236670892133077 |
| **bootstrap** | sound feature judg. | sham | TMS$pIPL | GPT2$surprisal | -0.96 | -0.81 | -0.8857601589286543 | 0.000999000999000999 | 0.0032236670892133077 |
| **bootstrap** | sound feature judg. | sham | TMS$pIPL | Word$frequency$(neg. sum) | -0.67 | -0.54 | -0.6029210507076723 | 0.000999000999000999 | 0.0032236670892133077 |
| **bootstrap** | sound feature judg. | sham | TMS$pIPL | Word$length$(sum) | -1.25 | -1.09 | -1.1725164949817355 | 0.000999000999000999 | 0.0032236670892133077 |
| **bootstrap** | picture naming w/ interf | vertex | TMS$pIFG | wac$PPMI | -1.74 | -1.55 | -1.6472164540836678 | 0.000999000999000999 | 0.0032236670892133077 |
| **bootstrap** | picture naming w/ interf | vertex | TMS$aIFG | wac$PPMI | -1.88 | -1.68 | -1.7779996049580187 | 0.000999000999000999 | 0.0032236670892133077 |
| **bootstrap** | picture naming w/ interf | vertex | TMS$pIFG | GPT2$surprisal | -0.44 | -0.31 | -0.37283370594610893 | 0.000999000999000999 | 0.0032236670892133077 |
| **bootstrap** | picture naming w/ interf | vertex | TMS$aIFG | GPT2$surprisal | -0.54 | -0.41 | -0.47379951772386714 | 0.000999000999000999 | 0.0032236670892133077 |
| **bootstrap** | picture naming w/ interf | vertex | TMS$pIFG | Word$frequency$(neg.$visual) | -0.5 | -0.37 | -0.4337242485521215 | 0.000999000999000999 | 0.0032236670892133077 |
| **bootstrap** | picture naming w/ interf | vertex | TMS$aIFG | Word$frequency$(neg.$visual) | 0.45 | 0.59 | 0.520233188043064 | 0.002997002997002997 | 0.006538915629824721 |
| **bootstrap** | picture naming w/ interf | vertex | TMS$pIFG | Word$frequency$(neg.$uttered) | -1.02 | -0.87 | -0.9421635948706404 | 0.000999000999000999 | 0.0032236670892133077 |
| **bootstrap** | picture naming w/ interf | vertex | TMS$aIFG | Word$frequency$(neg.$uttered) | -0.02 | 0.11 | 0.04425725635553081 | 0.34965034965034963 | 0.5009915457676651 |
| **bootstrap** | semantic production | vertex | TMS$pIFG | Word$frequency$(neg. sum) | -1.31 | -1.15 | -1.2294799493952118 | 0.000999000999000999 | 0.0032236670892133077 |
| **bootstrap** | semantic production | vertex | TMS$aIFG | Word$frequency$(neg. sum) | 0.51 | 0.64 | 0.5743482386916938 | 0.01098901098901099 | 0.022445639466916065 |
| **bootstrap** | semantic production | vertex | TMS$pIFG | Word$length$(visual) | 0.29 | 0.41 | 0.3500516518942549 | 0.000999000999000999 | 0.0032236670892133077 |
| **bootstrap** | semantic production | vertex | TMS$aIFG | Word$length$(visual) | -0.28 | -0.15 | -0.2124936368397289 | 0.6923076923076923 | 0.812985179957657 |
| **bootstrap** | semantic production | vertex | TMS$pIFG | Word$length$(uttered) | 0.37 | 0.5 | 0.4385422442310122 | 0.002997002997002997 | 0.006538915629824721 |
| **bootstrap** | semantic production | vertex | TMS$aIFG | Word$length$(uttered) | 1.43 | 1.61 | 1.5217499143185884 | 0.000999000999000999 | 0.0032236670892133077 |
| **bootstrap** | semantic production | vertex | TMS$pIFG | Word$length$(sum) | 1.03 | 1.19 | 1.1089269160716058 | 0.000999000999000999 | 0.0032236670892133077 |
| **bootstrap** | semantic production | vertex | TMS$aIFG | Word$length$(sum) | 1.11 | 1.27 | 1.186976758970205 | 0.000999000999000999 | 0.0032236670892133077 |
| **bootstrap** | semantic production | vertex | TMS$pMTG | wac$PPMI | -0.93 | -0.78 | -0.8558500311199692 | 0.000999000999000999 | 0.0032236670892133077 |
| **bootstrap** | semantic production | vertex | TMS$pMTG | GPT2$surprisal | -0.86 | -0.71 | -0.7854756227292347 | 0.000999000999000999 | 0.0032236670892133077 |
| **bootstrap** | semantic production | vertex | TMS$pMTG | Word$frequency$(neg.$visual) | 0.36 | 0.49 | 0.4255235907203821 | 0.000999000999000999 | 0.0032236670892133077 |
| **bootstrap** | semantic production | vertex | TMS$pMTG | Word$frequency$(neg.$uttered) | -0.25 | -0.13 | -0.18850488125004508 | 0.01098901098901099 | 0.022445639466916065 |
| **bootstrap** | semantic production | vertex | TMS$pMTG | Word$frequency$(neg. sum) | 0.23 | 0.35 | 0.28862305170246716 | 0.000999000999000999 | 0.0032236670892133077 |
| **bootstrap** | semantic production | vertex | TMS$pMTG | Word$length$(visual) | 0.63 | 0.77 | 0.7017101105995593 | 0.000999000999000999 | 0.0032236670892133077 |
| **bootstrap** | semantic production | vertex | TMS$pMTG | Word$length$(uttered) | 0.02 | 0.15 | 0.08466793092697913 | 0.12887112887112886 | 0.21872863978127133 |
| **bootstrap** | semantic production | vertex | TMS$pMTG | Word$length$(sum) | 0.41 | 0.54 | 0.4713819479972672 | 0.000999000999000999 | 0.0032236670892133077 |
| **bootstrap** | quantity priming | vertex | TMS$rCereb | wac$PPMI | 1.14 | 1.31 | 1.223780215429802 | 0.000999000999000999 | 0.0032236670892133077 |
| **bootstrap** | quantity priming | vertex | TMS$rCereb | GPT2$surprisal | 0.58 | 0.71 | 0.6438185286471736 | 0.013986013986013986 | 0.028118478380258487 |
| **bootstrap** | quantity priming | vertex | TMS$rCereb | Word$frequency$(neg.$first w.) | -0.16 | -0.03 | -0.09572432923530212 | 0.056943056943056944 | 0.10363096619020791 |
| **bootstrap** | quantity priming | vertex | TMS$rCereb | Word$frequency$(neg.$second w.) | -0.27 | -0.15 | -0.21091134633699013 | 0.000999000999000999 | 0.0032236670892133077 |
| **bootstrap** | quantity priming | vertex | TMS$rCereb | Word$frequency$(neg. sum) | -0.29 | -0.17 | -0.22852799490509543 | 0.000999000999000999 | 0.0032236670892133077 |
| **bootstrap** | quantity priming | vertex | TMS$rCereb | Word$length$(first w.) | 0.06 | 0.18 | 0.12189132326556665 | 0.007992007992007992 | 0.01658881658881659 |
| **bootstrap** | quantity priming | vertex | TMS$rCereb | Word$length$(second w.) | 0.31 | 0.44 | 0.37217511155211347 | 0.000999000999000999 | 0.0032236670892133077 |
| **bootstrap** | quantity priming | vertex | TMS$rCereb | Word$length$(sum) | 0.18 | 0.3 | 0.24109825225349366 | 0.02097902097902098 | 0.04089311703524901 |
| **bootstrap** | quantity priming | vertex | TMS$rSATL | wac$PPMI | 1.96 | 2.18 | 2.0730148920979534 | 0.000999000999000999 | 0.0032236670892133077 |
| **bootstrap** | quantity priming | vertex | TMS$rIPS | wac$PPMI | 2.42 | 2.68 | 2.549034403037493 | 0.000999000999000999 | 0.0032236670892133077 |
| **bootstrap** | quantity priming | vertex | TMS$rSATL | GPT2$surprisal | -0.63 | -0.49 | -0.5605142427688048 | 0.000999000999000999 | 0.0032236670892133077 |
| **bootstrap** | quantity priming | vertex | TMS$rIPS | GPT2$surprisal | -0.37 | -0.25 | -0.30942927961626177 | 0.000999000999000999 | 0.0032236670892133077 |
| **bootstrap** | quantity priming | vertex | TMS$rSATL | Word$frequency$(neg.$first w.) | -1.86 | -1.67 | -1.765655592297993 | 0.000999000999000999 | 0.0032236670892133077 |
| **bootstrap** | quantity priming | vertex | TMS$rIPS | Word$frequency$(neg.$first w.) | -3.19 | -2.9 | -3.0460520065611307 | 0.000999000999000999 | 0.0032236670892133077 |
| **bootstrap** | quantity priming | vertex | TMS$rSATL | Word$frequency$(neg.$second w.) | 0.86 | 1.01 | 0.9393033054431862 | 0.000999000999000999 | 0.0032236670892133077 |
| **bootstrap** | quantity priming | vertex | TMS$rIPS | Word$frequency$(neg.$second w.) | 0.14 | 0.27 | 0.20541632025394987 | 0.000999000999000999 | 0.0032236670892133077 |
| **bootstrap** | social priming | vertex | TMS$rSATL | Word$frequency$(neg. sum) | -1.13 | -0.98 | -1.052529615576613 | 0.000999000999000999 | 0.0032236670892133077 |
| **bootstrap** | social priming | vertex | TMS$rIPS | Word$frequency$(neg. sum) | -2.76 | -2.5 | -2.6266712613850065 | 0.000999000999000999 | 0.0032236670892133077 |
| **bootstrap** | social priming | vertex | TMS$rSATL | Word$length$(first w.) | -1.84 | -1.64 | -1.741708362129087 | 0.000999000999000999 | 0.0032236670892133077 |
| **bootstrap** | social priming | vertex | TMS$rIPS | Word$length$(first w.) | -3.2 | -2.9 | -3.051846147794543 | 0.000999000999000999 | 0.0032236670892133077 |
| **bootstrap** | social priming | vertex | TMS$rSATL | Word$length$(second w.) | 1.61 | 1.81 | 1.7106433418889968 | 0.000999000999000999 | 0.0032236670892133077 |
| **bootstrap** | social priming | vertex | TMS$rIPS | Word$length$(second w.) | -1.91 | -1.71 | -1.8069028141181096 | 0.000999000999000999 | 0.0032236670892133077 |
| **bootstrap** | social priming | vertex | TMS$rSATL | Word$length$(sum) | 1.33 | 1.5 | 1.4161335347128974 | 0.000999000999000999 | 0.0032236670892133077 |
| **bootstrap** | social priming | vertex | TMS$rIPS | Word$length$(sum) | -2.54 | -2.3 | -2.419298308334551 | 0.000999000999000999 | 0.0032236670892133077 |
| **bootstrap** | social priming | vertex | TMS$rSATL | wac$PPMI | -1.45 | -1.28 | -1.3627418842993826 | 0.000999000999000999 | 0.0032236670892133077 |
| **bootstrap** | social priming | vertex | TMS$rIPS | wac$PPMI | -2.24 | -2.01 | -2.1232121378161355 | 0.000999000999000999 | 0.0032236670892133077 |
| **bootstrap** | social priming | vertex | TMS$rSATL | GPT2$surprisal | -0.97 | -0.83 | -0.8987310395635648 | 0.000999000999000999 | 0.0032236670892133077 |
| **bootstrap** | social priming | vertex | TMS$rIPS | GPT2$surprisal | -1.14 | -0.99 | -1.06269195262092 | 0.000999000999000999 | 0.0032236670892133077 |
| **bootstrap** | social priming | vertex | TMS$rSATL | Word$frequency$(neg.$first w.) | 3.75 | 4.12 | 3.9374011143837255 | 0.000999000999000999 | 0.0032236670892133077 |
| **bootstrap** | social priming | vertex | TMS$rIPS | Word$frequency$(neg.$first w.) | 2.79 | 3.08 | 2.934475477107718 | 0.000999000999000999 | 0.0032236670892133077 |
| **bootstrap** | social priming | vertex | TMS$rSATL | Word$frequency$(neg.$second w.) | -0.02 | 0.1 | 0.040214118288537666 | 0.26873126873126874 | 0.4062708944598708 |
| **bootstrap** | social priming | vertex | TMS$rIPS | Word$frequency$(neg.$second w.) | -1.58 | -1.4 | -1.490082342758317 | 0.000999000999000999 | 0.0032236670892133077 |
| **bootstrap** | semantic relatedness judg. | vertex | TMS$rSATL | Word$frequency$(neg. sum) | 3.11 | 3.42 | 3.2679482264066837 | 0.000999000999000999 | 0.0032236670892133077 |
| **bootstrap** | semantic relatedness judg. | vertex | TMS$rIPS | Word$frequency$(neg. sum) | 1.73 | 1.94 | 1.8361733405752767 | 0.000999000999000999 | 0.0032236670892133077 |
| **bootstrap** | semantic relatedness judg. | vertex | TMS$rSATL | Word$length$(first w.) | 3.89 | 4.27 | 4.078080183387404 | 0.000999000999000999 | 0.0032236670892133077 |
| **bootstrap** | semantic relatedness judg. | vertex | TMS$rIPS | Word$length$(first w.) | 2.92 | 3.22 | 3.073072604224002 | 0.000999000999000999 | 0.0032236670892133077 |
| **bootstrap** | semantic relatedness judg. | vertex | TMS$rSATL | Word$length$(second w.) | -0.68 | -0.54 | -0.6120338026124811 | 0.002997002997002997 | 0.006538915629824721 |
| **bootstrap** | semantic relatedness judg. | vertex | TMS$rIPS | Word$length$(second w.) | -0.28 | -0.15 | -0.21696751345164028 | 0.5444555444555444 | 0.6922878446057253 |
| **bootstrap** | semantic relatedness judg. | vertex | TMS$rSATL | Word$length$(sum) | 0.03 | 0.15 | 0.091119450665283 | 0.21978021978021978 | 0.34168260888908664 |
| **bootstrap** | semantic relatedness judg. | vertex | TMS$rIPS | Word$length$(sum) | 0.21 | 0.34 | 0.2751168735778509 | 0.03596403596403597 | 0.06671589280284933 |
